# Theoretical properties of nearest-neighbor distance distributions and novel metrics for high dimensional bioinformatics data

**DOI:** 10.1101/857821

**Authors:** Bryan A. Dawkins, Trang T. Le, Brett A. McKinney

## Abstract

The performance of nearest-neighbor feature selection and prediction methods depends on the metric for computing neighborhoods and the distribution properties of the underlying data. The effects of the distribution and metric, as well as the presence of correlation and interactions, are reflected in the expected moments of the distribution of pairwise distances. We derive general analytical expressions for the mean and variance of pairwise distances for *L*_*q*_ metrics for normal and uniform random data with *p* attributes and *m* instances. We use extreme value theory to derive results for metrics that are normalized by the range of each attribute (max – min). In addition to these expressions for continuous data, we derive similar analytical formulas for a new metric for genetic variants (categorical data) in genome-wide association studies (GWAS). The genetic distance distributions account for minor allele frequency and transition/transversion ratio. We introduce a new metric for resting-state functional MRI data (rs-fMRI) and derive its distance properties. This metric is applicable to correlation-based predictors derived from time series data. Derivations assume independent data, but empirically we also consider the effect of correlation. These analytical results and new metrics can be used to inform the optimization of nearest neighbor methods for a broad range of studies including gene expression, GWAS, and fMRI data. The summary of distribution moments and detailed derivations provide a resource for understanding the distance properties for various metrics and data types.

## Introduction

Statistical models can deviate from expected behavior depending on whether certain properties of the underlying data are satisfied, such as being normally distributed. The expected behavior of nearest neighbor models is further influenced by the choice of metric, such as Euclidean or Manhattan. For random normal data (𝒩(0, 1)), for example, the variance of the pairwise distances of a Manhattan metric is proportional to the number of attributes (*p*) whereas the variance is constant for a Euclidean metric. Relief methods [1–3] and nearest-neighbor projected distance regression (NDPR) [4] use nearest neighbors to compute attribute importance scores and often use adaptive neighborhoods that rely on the mean and variance of the distance distribution. The ability of this class of methods to identify association effects, like main effects or interaction effects, depends on parameters such as neighborhood radii or number of neighbors k [12, 13]. Thus, knowledge of the expected values for a given metric and data distribution may improve the performance of these feature selection methods by informing the selection of neighborhood parameters.

For continuous data, the metrics most commonly used in nearest neighbor methods are *L*_*q*_ with *q* = 1 (Manhattan) or *q* = 2 (Euclidean). For data from standard normal (𝒩(0, 1)) or standard uniform (𝒰(0, 1)) distributions, the asymptotic behavior of the *L*_*q*_ metrics is known. However, detailed derivations of these distance distribution asymptotics are not readily available in the literature. We provide detailed derivations of generalized expressions parameterized by metric *q*, attributes *p*, and samples *m*, and we extend the derivations to *L*_*q*_ metrics normalized by the range of the attributes using Extreme Value Theory (EVT). These range (max-min) normalized metrics are often used in Relief-based algorithms [3].

For genome-wide association study (GWAS) data, which is categorical, various metrics have been developed for feature selection and for computing similarity between individuals based on shared genetic variation. We build on the mathematical formalism for continuous data to derive the asymptotic properties of various categorical data metrics for GWAS. We derive asymptotic formulas for the mean and variance for three recently introduced GWAS metrics [5]. These metrics were developed for Relief-based feature selection to account for binary genotype differences (two levels), allelic differences (three levels), and transition/transversion differences (five levels). The mean and variance expressions we derive for these multi-level categorical data types are parameterized by the minor allele frequency and the transition/transversion ratio.

Resting-state fMRI (rs-fMRI) is a growing application area for machine learning and feature selection [6–9], which involves correlation data derived from time-series brain activity. For a given subject, a correlation or similar matrix is computed between brain regions of interest (ROIs) from their time series. Each time series represents functional activity of the ROI while the subject is not performing any task, and the ROI typically corresponds to a region with known function for emotion or cognition. Thus, the dataset consists of pairwise ROI correlations for each of the *m* subjects. Nearest-neighbor based feature selection was applied to rs-fMRI in the private evaporative cooling method [10], where the predictors were pairwise correlations between ROIs. The use of pairwise correlation predictors is a common practice because of convenience and differential connectivity between brain regions may be of biological importance [11]. However, one may be interested in the importance of attributes at the ROI level. Thus, in the current study we introduce a new metric to be used in NPDR [4] with resting state correlation matrices that provides attribute importance for ROIs. This metric is applicable to general time series derived correlation data, and we derive asymptotic estimates for the mean and variance of distance distributions for our new ts-corr based metric.

In Section 1, we introduce preliminary notation and apply the Central Limit Theorem (CLT) and the Delta Method to derive asymptotics for pairwise distances. In Section 2, we present general derivations for continuously distributed data sets with *m* instances and *p* attributes. We begin with the cases of standard normal (𝒩(0, 1)) and standard uniform (𝒰(0, 1)) data distributions, but we derive analytical expressions parameterized by *q, p*, and *m*. In Section 2.4 we use Extreme Value Theory (EVT) to derive attribute range-normalized (max-min) versions of *L*_*q*_ metrics. In Section 3, we extend the derivations to categorical data with a binomial distribution for GWAS data with multiple metric types. In Section 4, we present a new time series correlation-based distance metric, with a particular emphasis on rs-fMRI data, and we derive the corresponding asymptotic distance distribution results. Lastly, in Section 6, we demonstrate the effect of correlation in the attribute space on distance distributional properties.

## 1 Limit distribution for *L*_*q*_ on null data

In the application of nearest-neighbor distance-based methods to continuous data, the distance between instances (*i, j* ∈ ℐ, |ℐ| = *m*) in the data set *X*^*m*×*p*^ of *m* instances and *p* attributes (or features) is calculated in the space of all attributes (*a* ∈ 𝒜, |𝒜| = *p*) using a metric such as

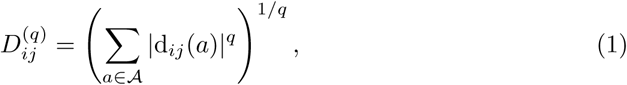

which is typically Manhattan (*q* = 1) in Relief-based methods but may also be Euclidean (*q* = 2). We use the terms “feature” and “attribute” interchangeably for the remainder of this work. The quantity d_*ij*_(*a*), known as a “diff” in Relief literature, is the projection of the distance between instances *i* and *j* onto the attribute *a* dimension. The function d_*ij*_(*a*) supports any type of attributes (e.g., numeric and categorical). For example, the projected difference between two instances *i* and *j* for a continuous numeric (d^num^) attribute *a* may be

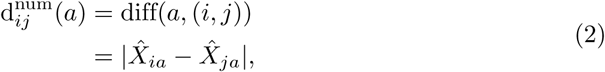

where 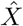 represents the standardized data matrix *X*. We use a simplified d_*ij*_(*a*) notation in place of the diff(*a*, (*i, j*)) notation that is customary in Relief-based methods. In NPDR, we omit the division by max(*a*) − min(*a*) used by Relief to constrain scores to the interval from −1 to 1, where max(*a*) = max_*k*∈ℐ_ {*X*_*ka*_} and min(*a*) = min *k* ∈ ℐ{*X*_*ka*_}. The numeric 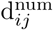(*a*) projection is simply the absolute difference between row elements *i* and *j* of the data matrix *X*^*m*×*p*^ for the attribute column *a*.

All derivations in the following sections are applicable to nearest-neighbor distance-based methods in general, which includes not only NPDR, but also Relief-based algorithms. Each of these methods uses a distance metric (Eq. 1) to compute neighbors for each instance *I* ∈ ℐ. Therefore, our derivations of asymptotic distance distributions are applicable to all methods that compute neighbors in order to weight features. The predictors used by NPDR, however, are the one-dimensional projected distances between two instances *i, j* ∈ ℐ (Eq. 2). Hence, all asymptotic estimates we derive for diff metrics (Eq. 2) are particularly relevant to NPDR. Since the standard distance metric (Eq. 1) is a function of the one-dimensional projection (Eq. 2), asymptotic estimates derived for this projection (Eq. 2) are implicitly relevant to older nearest-neighbor distance-based methods like Relief-based algorithms. We proceed in the following section by applying the Classical Central Limit Theorem and the Delta Method to derive the limit distribution of pairwise distances on any data distribution that is induced by the standard distance metric (Eq. 1).

### 1.1 Asymptotic normality of pairwise distances

Suppose that 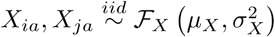 for two fixed and distinct instances *i, j* ∈ ℐ and a fixed attribute *a* ∈ 𝒜. ℱ_*X*_ represents any data distribution with mean *µ*_*X*_ and variance 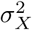.

It is clear that |*X*_*ia*_ − *X*_*ja*_|^*q*^ = |d_*ij*_(*a*)|^*q*^ is another random variable. Let 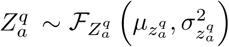 be the random variable such that

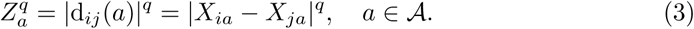

Furthermore, the collection 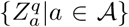 is a random sample of size *p* of mutually independent random variables. Hence, the sum of 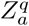 over all *a* ∈ 𝒜 is asymptotically normal by the Classical Central Limit Theorem (CCLT). More explicitly, this implies that

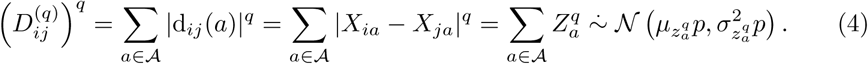

Consider the smooth function *g*(*z*) = *z*^1*/q*^ that is continuously differentiable for *z* > 0. Assuming that 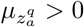, the Delta Method [14] can be applied to show that

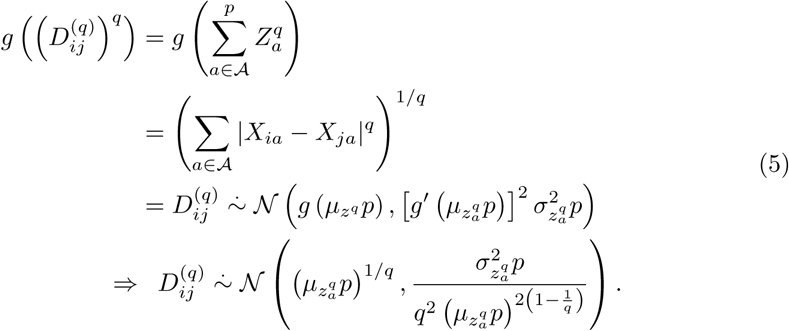

Therefore, the distance between two fixed, distinct instances *i* and *j* given by Eq. 1 is asymptotically normal. Specifically, when *q* = 2, the distribution of 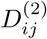 asymptotically approaches 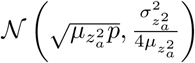. When *p* is small, however, we observe empirically that a closer estimate of the sample mean is

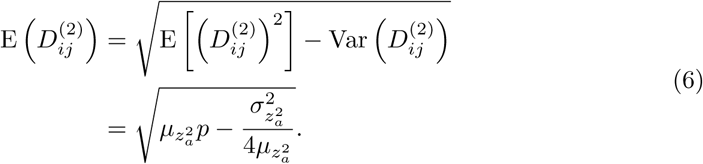

We estimate rate of convergence to normality for Euclidean (*q* = 2) and Manhattan (*q* = 1) metrics by comparing the distribution of pairwise distances in simulated data to a Gaussian (Fig. 1). We compute the distance between all pairs of instances in simulated datasets of uniformly distributed random data. We simulate data with fixed *m* = 100 instances, and, by varying the number of attributes (*p* = 10, 100, 10000), we observe rapid convergence to Gaussian. For *p* as low as 10 attributes, Gaussian is a good approximation. The number of attributes in bioinformatics data is typically quite large, at least on the order of 10^3^. The Euclidean metric has stronger convergence to a Gaussian than Manhattan. This may be due to Euclidean’s use of the square root, which is a common transformation of data in statistics. Normality was assessed using the Shapiro-Wilk test.

**Fig 1.**
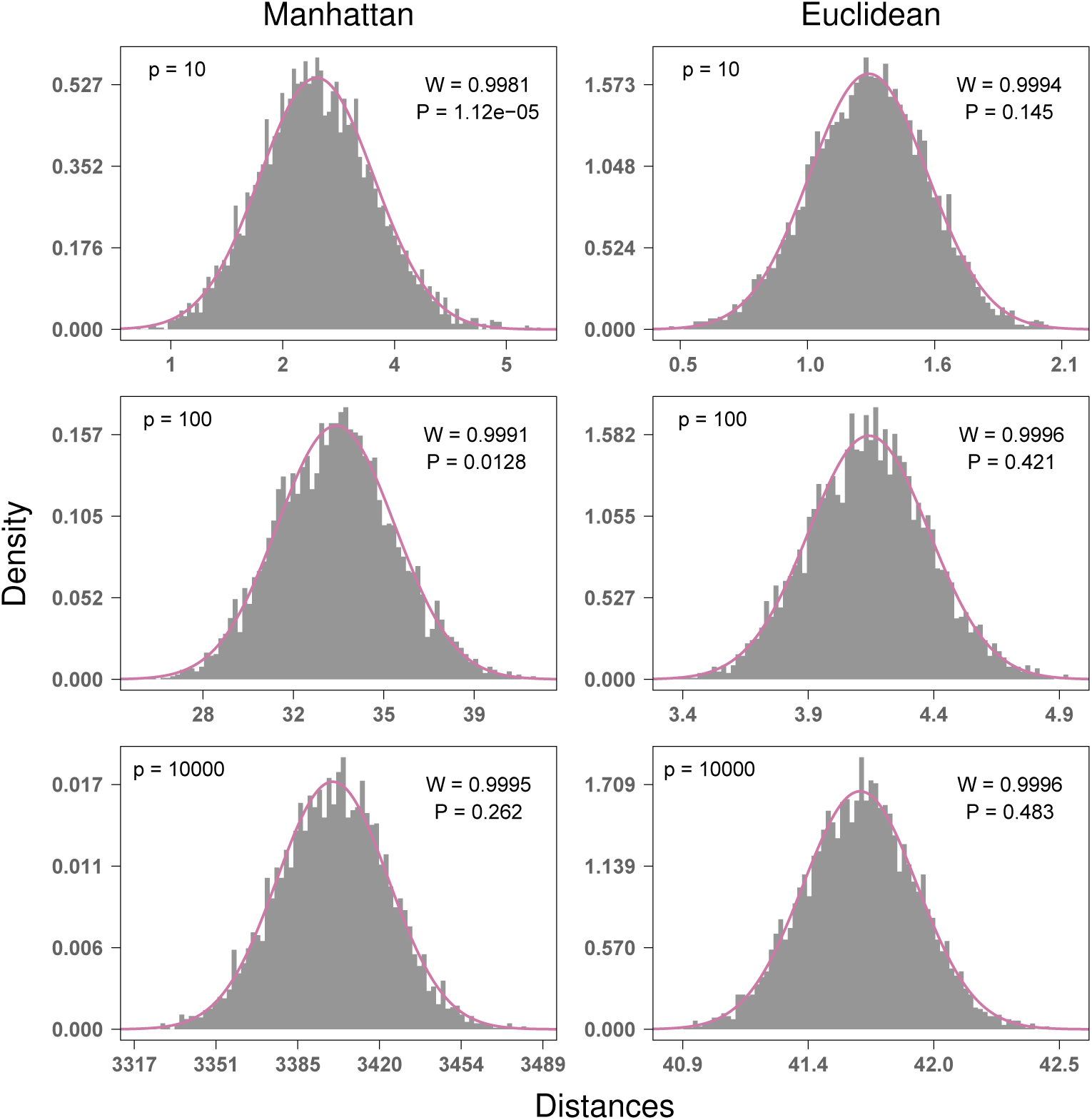
Convergence to Gaussian for Manhattan and Euclidean distances for simulated standard uniform data with *m* = 100 instances and *p* = 10, 100, and 10000 attributes. Convergence to Gaussian occurs rapidly with increasing *p*, and Gaussian is a good approximation for *p* as low as 10 attributes. The number of attributes in bioinformatics data is typically much larger, at least on the order of 10^3^. The Euclidean metric has stronger convergence to normal than Manhattan. P values from Shapiro-Wilk test, where the null hypothesis is a Gaussian distribution.

To show asymptotic normality of distances, we did not specify whether the data distribution ℱ_*X*_ was discrete or continuous. This is because asymptotic normality is a general phenomenon in high attribute dimension *p* for any data distribution ℱ_*X*_ satisfying the assumptions we have made. Therefore, the simulated distances we have shown (Fig. 1) has an analogous representation for discrete data, as well as all other continuous data distributions.

For distance based learning methods, all pairwise distances are used to determine relative importances for attributes. The collection of all distances above the diagonal in an *m* × *m* distance matrix does not satisfy the independence assumption used in the previous derivations. This is because of the redundancy that is inherent to the distance matrix calculation. However, this collection is still asymptotically normal with mean and variance approximately equal to those we have previously given (Eq. 5). In the next section, we assume actual data distributions in order to define more specific general formulas for standard *L*_*q*_ and max-min normalized *L*_*q*_ metrics. We also derive asymptotic moments for a new discrete metric in GWAS data and a new metric for time series correlation-based data, such as, resting-state fMRI.

For distance based learning methods, all pairwise distances are used to determine relative importances for attributes. The collection of all distances above the diagonal in an *m* × *m* distance matrix does not satisfy the independence assumption used in the previous derivations. This is because of the redundancy that is inherent to the distance matrix calculation. However, this collection is still asymptotically normal with mean and variance approximately equal to those we have previously given (Eq. 5). In the next section, we assume actual data distributions in order to define more specific general formulas for standard *L*_*q*_ and max-min normalized *L*_*q*_ metrics. We also derive asymptotic moments for a new discrete metric in GWAS data and a new metric for time series correlation-based data, such as, resting-state fMRI.

## 2 *L*_*q*_ metric moments for continuous data distributions

In this section, we begin by deriving general formulas for asymptotic means and variances of the *L*_*q*_ distance (Eq. 1) for standard normal and standard uniform data. With our general formulas for continuous data, we compute moments associated with Manhattan (*L*_1_) and Euclidean (*L*_2_) metrics. We then consider the max-min normalized version of the *L*_*q*_ distance, where the magnitude difference (Eq. 2) is divided by the range of each attribute *a*. Using Extreme Value Theory (EVT), we derive formulas for the moments of attribute range in standard normal and standard uniform data. Transitioning into discrete data distributions relevant to GWAS, we derive asymptotic moments for two well known metrics and one new metric. In addition, we derive distance asymptotics for time series correlation-based data, such as, resting-state fMRI.

### 2.1 Distribution of |d_*ij*_(*a*)|^*q*^ = |*X*_*ia*_ − *X*_*ja*_|^*q*^

Suppose that 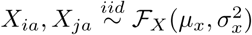 and define 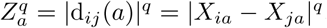, where *a* ∈ 𝒜 and |𝒜| = *p*. In order to find the distribution of 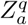, we will use the following theorem given in [15].

#### Theorem 2.1

*Let f* (*x*) *be the value of the probability density of the continuous random variable X at x. If the function given by y* = *u*(*x*) *is differentiable and either increasing or decreasing for all values within the range of X for which f* (*x*) ≠ 0, *then, for these values of x, the equation y* = *u*(*x*) *can be uniquely solved for x to give x* = *w*(*y*), *and for the corresponding values of y the probability density of Y* = *u*(*X*) *is given by*

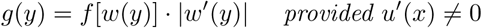

*Elsewhere, g*(*y*) = 0.

We have the following cases that result from solving for *X*_*ja*_ in the equation given by 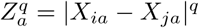.

i. Suppose that 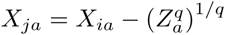. Based on the iid assumption for *X*_*ia*_ and *X*_*ja*_, it follows from Thm. 2.1 that the joint density function *g*^(1)^ of *X*_*ia*_ and 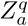 is given by

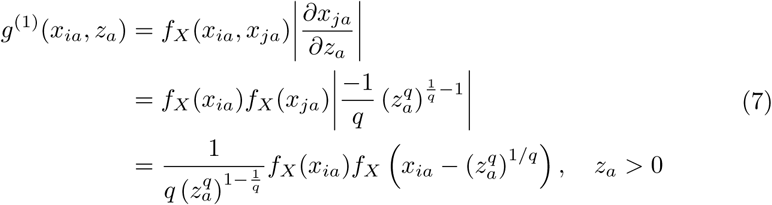 The density function 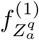 of 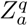 is then defined as

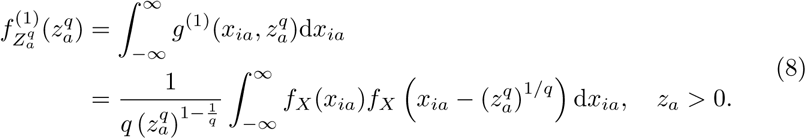
ii. Suppose that 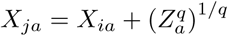. Based on the iid assumption for *X*_*ia*_ and *X*_*ja*_, it follows from Thm. 2.1 that the joint density function *g*^(2)^ of *X*_*ia*_ and *Z*_*a*_ is given by

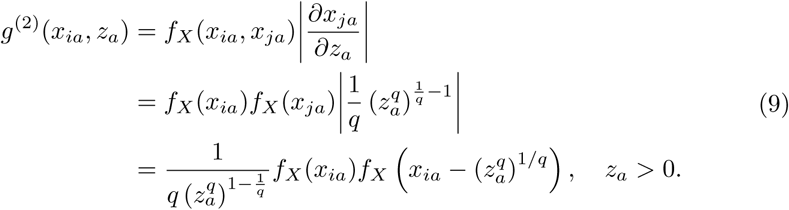 The density function 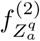 of 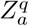 is then defined as

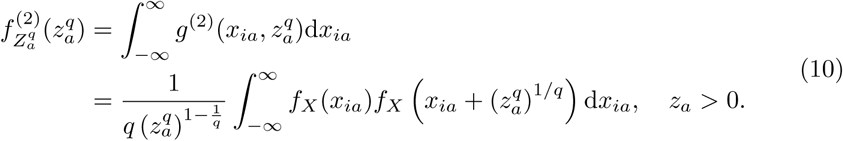

Let 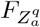 denote the distribution function of the random variable 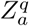. Furthermore, we define the events *E*^(1)^ and *E*^(2)^ as

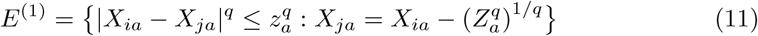

and

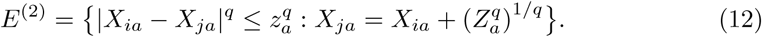

Then it follows from fundamental rules of probability that

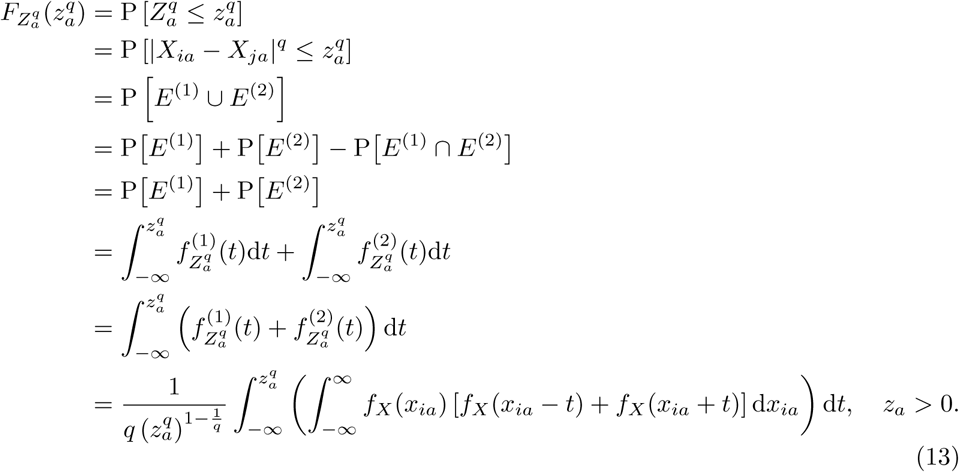

It follows directly from the previous result (Eq. 13) that the density function of the random variable 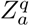 is given by

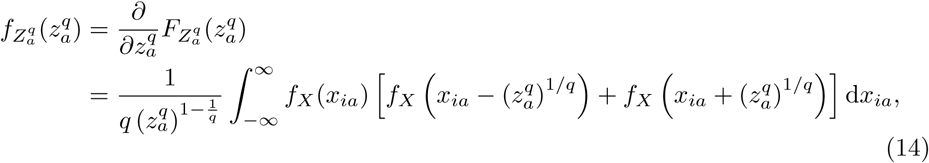

where *z*_*a*_ *>* 0.

Using the previous result (Eq. 14), we can compute the mean and variance of the random variable 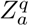 as

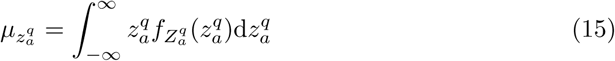

and

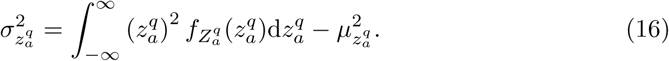

It follows immediately from the mean (Eq. 15) and variance (Eq. 16) and the Classical Central Limit Theorem (CCLT) that

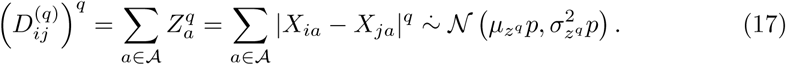

Applying the convergence result we derived previously (Eq. 5), the distribution of 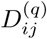 is given by

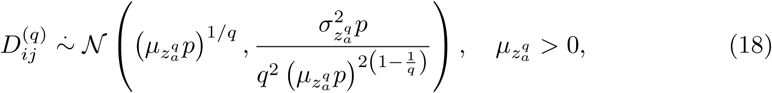

where we have an improved estimate of the mean for *q* = 2 (Eq. 6).

#### 2.1.1 Standard normal data

If 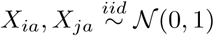, then the marginal density functions with respect to *X* for *X*_*ia*_, 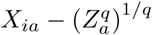, and 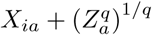 are defined as

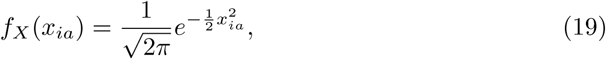

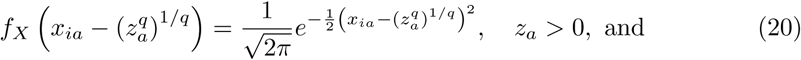

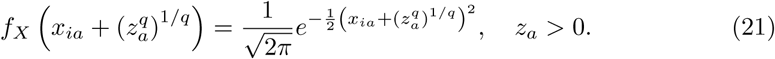

Substituting these marginal densities (Eqs. 19-21) into the general density function for 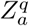 (Eq. 14) and completing the square on *x*_*ia*_ in the exponents, we have

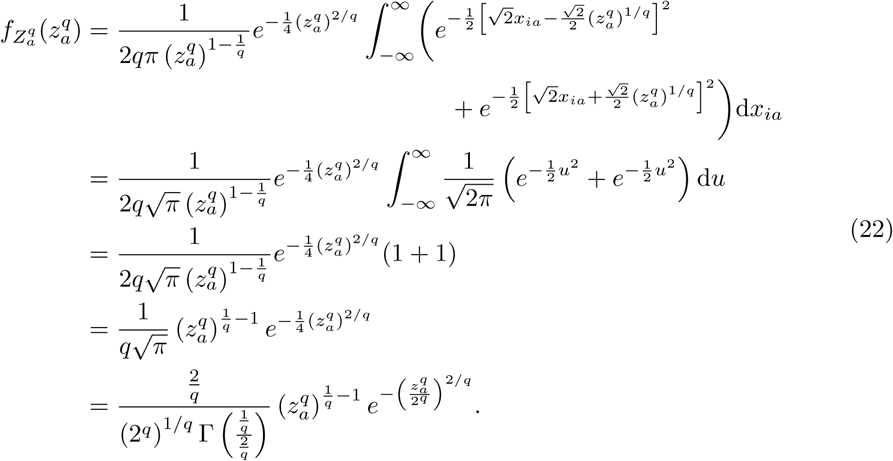

The density function given previously (Eq. 22) is a Generalized Gamma density with parameters 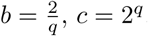, and 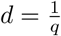. This distribution has mean and variance given by

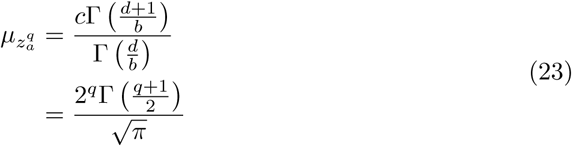

and

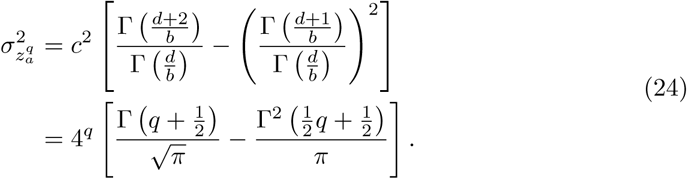

By linearity of the expected value and variance operators under the iid assumption, the mean (Eq. 23) and variance (Eq. 24) of the random variable 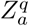 allow the *p*-dimensional mean and variance of the 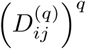 distribution to be computed directly as

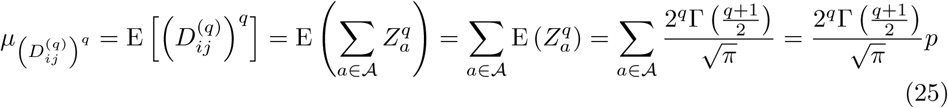

and

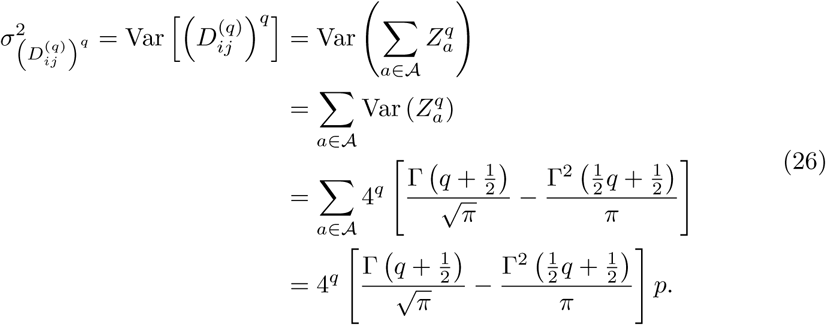

Therefore, the asymptotic distribution of 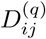 for standard normal data is

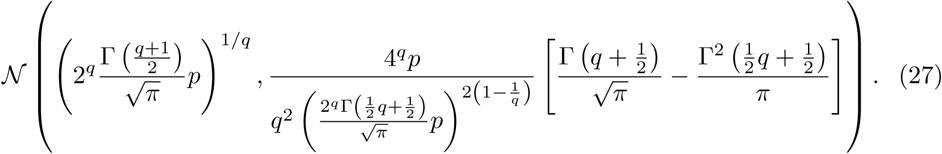

We provide a summary table of the moment estimates (Eq. 27) for the *L*_*q*_ metric on standard normal data (Tab. 1). The summary is organized by data type, type of statistic (mean or variance), and corresponding asymptotic formula. In the next section, we derive the *L*_*q*_ distance distribution on standard uniform data in a similar fashion.

**Table 1.** Summary of distance distribution derivations for standard normal (𝒩(0, 1)) and standard uniform (𝒰(0, 1)) data. Asymptotic estimates are given for both standard (Eq. 1) and max-min normalized (Eq. 55) q-metrics. These estimates are relevant for all *q* ∈ ℕ and *p* ≫ 1 for which the normality assumption of distances holds.

| q-Metric | Data | Stat | Formula (Eq. #) |
| --- | --- | --- | --- |
| standard<br>(Eq. 1) | $\mathcal{N}(0, 1)$ | mean | $\left( \frac{2^q \Gamma(\frac{q+1}{2}) p}{\sqrt{\pi}} \right)^{1/q} \quad (25)$ |
| | $\mathcal{N}(0, 1)$ | variance | $\frac{4^q p}{q^2 \left( \frac{2^q \Gamma(\frac{1}{2} q + \frac{1}{2})}{\sqrt{\pi}} p \right)^{2(1-\frac{1}{q})}} \left[ \frac{\Gamma(q+\frac{1}{2})}{\sqrt{\pi}} - \frac{\Gamma^2(\frac{1}{2} q + \frac{1}{2})}{\pi} \right] \quad (26)$ |
| | $\mathcal{U}(0, 1)$ | mean | $\left( \frac{2p}{(q+2)(q+1)} \right)^{1/q} \quad (35)$ |
| | $\mathcal{U}(0, 1)$ | variance | $\frac{p}{q^2 \left( \frac{2p}{(q+2)(q+1)} \right)^{2(1-\frac{1}{q})}} \left[ \frac{1}{(q+1)(2q+1)} - \left( \frac{2}{(q+2)(q+1)} \right)^2 \right] \quad (36)$ |
| max-min<br>normalized<br>(Eq. 55) | $\mathcal{N}(0, 1)$ | mean | $\frac{\mu_{D_{ij}}^{(q)}}{2\mu_{\max}^{(1)}(m)} \quad (89)$<br>where $\mu_{D_{ij}}^{(q)}$ and $\mu_{\max}^{(1)}(m)$ are given by Eqs. 25 and 83 respectively. |
| | $\mathcal{N}(0, 1)$ | variance | $\frac{6\log(m)\sigma_{D_{ij}}^2{}^{(q)}}{\pi^2 + 24[\mu_{\max}^{(1)}(m)]^2 \log(m)} \quad (89)$<br>where $\sigma_{D_{ij}}^2{}^{(q)}$ and $\mu_{\max}^{(1)}(m)$ are given by Eqs. 25 and 83 respectively. |
| | $\mathcal{U}(0, 1)$ | mean | $\frac{(m+1)\mu_{D_{ij}}^{(q)}}{m-1} \quad (97)$<br>where $\mu_{D_{ij}}^{(q)}$ is given by Eq. 35 |
| | $\mathcal{U}(0, 1)$ | variance | $\frac{(m+2)(m+1)^2 \sigma_{D_{ij}}^2{}^{(q)}}{m^3 - m + 2} \quad (97)$<br>where $\sigma_{D_{ij}}^2{}^{(q)}$ is given by Eq. 36 |

#### 2.1.2 Standard uniform data

If 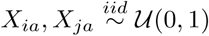, then the marginal density functions with respect to *X* for *X*_*ia*_, 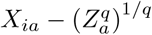, and 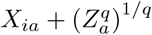 are defined as

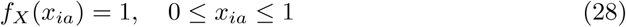

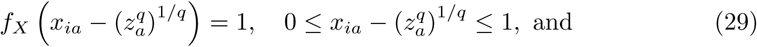

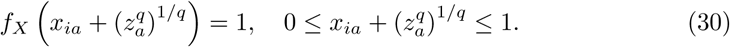

Substituting these marginal densities (Eqs. 28-30) into the more general density function for 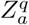 (Eq. 14), we have

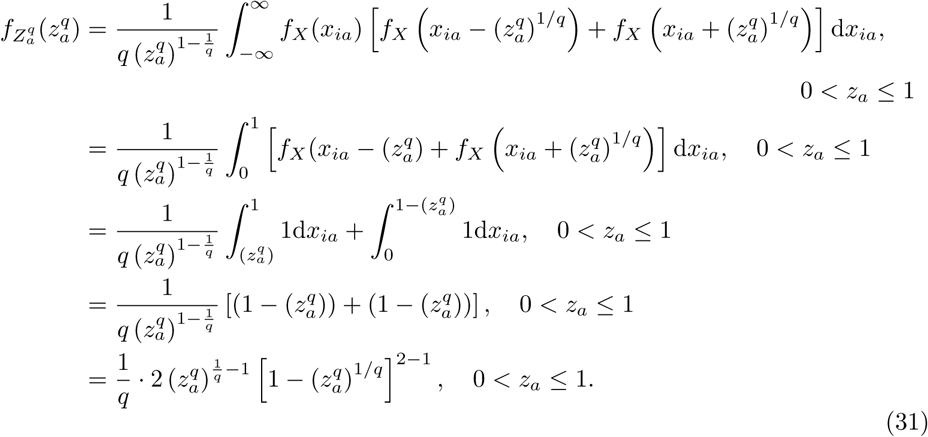

The previous density (Eq. 31) is a Kumaraswamy density with parameters 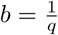 and *c* = 2 with moment generating function (MGF) given by

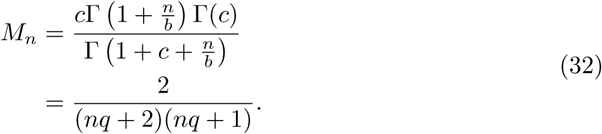

Using this MGF (Eq. 32), the mean and variance of 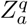 are computed as

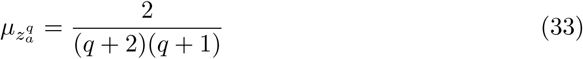

and

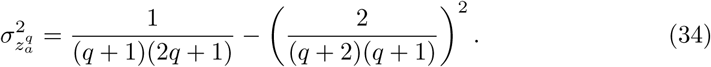

By linearity of the expected value and variance operators under the iid assumption, the mean (Eq. 33) and variance (Eq. 34) of the random variable 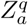 allow the *p*-dimensional mean and variance of the 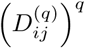 distribution to be computed directly as

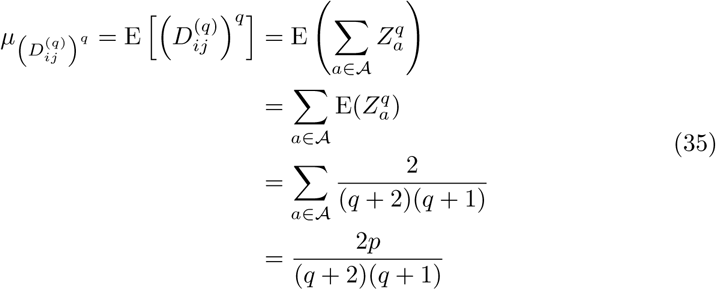

and

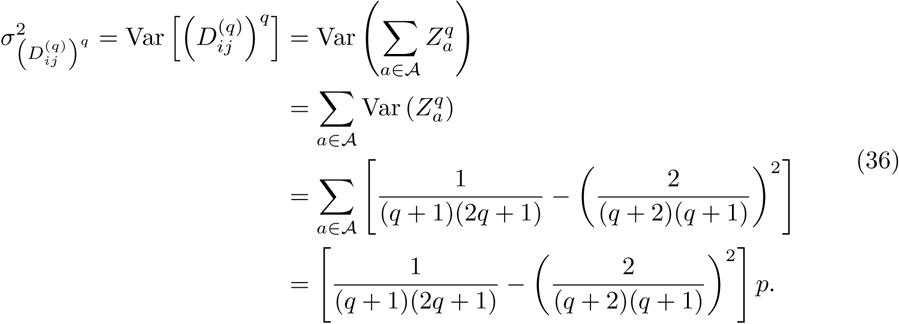

Therefore, the asymptotic distribution of 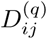 for standard uniform data is

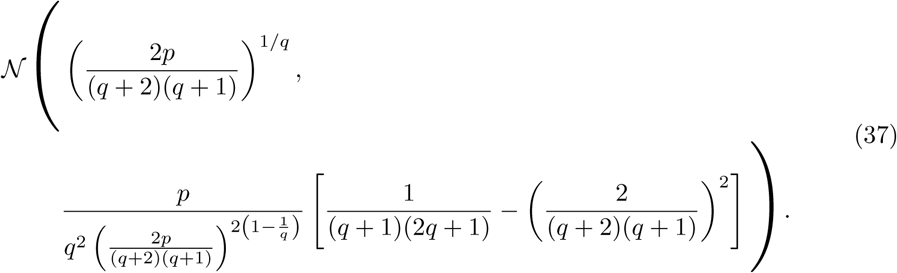

We provide a summary table of the moment estimates (Eq. 37) for the *L*_*q*_ metric on standard uniform data (Tab. 1). The summary is organized by data type, type of statistic (mean or variance), and corresponding asymptotic formula. In the next section, we use our general *L*_*q*_ distance distribution derivations to provide Manhattan (*q* = 1) and Euclidean (*q* = 2) asymptotic moments on both standard normal and standard uniform data. These are the most commonly applied metrics in the context of nearest-neighbor feature selection, so they are of particular interest.

### 2.2 Manhattan (*L*_1_)

With our general formulas for the asymptotic mean and variance (Eqs. 27 and 37) for any value of *q* ∈ ℕ, we can simply substitute a particular value of *q* in order to determine the asymptotic distribution of the corresponding distance *L*_*q*_ metric. We demonstrate this with the example of the Manhattan metric (*L*_1_) for standard normal and standard uniform data (Eq. 1, *q* = 1).

#### 2.2.1 Standard normal data

Substituting *q* = 1 into the asymptotic formula for the mean *L*_*q*_ distance (Eq. 27), we have the following for expected *L*_1_ distance between two independently sample instances *i, j* ∈ ℐ in standard normal data

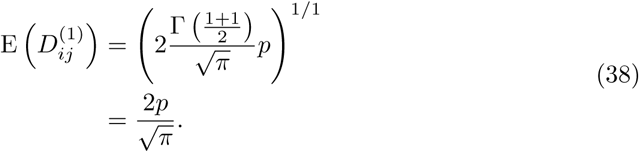

We see in the formula for the expected Manhattan distance (Eq. 38) that 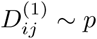 in the limit, which implies that this distance is unbounded as feature dimension *p* increases.

Substituting *q* = 1 into the formula for the asymptotic variance of 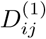 (Eq. 27) leads to the following

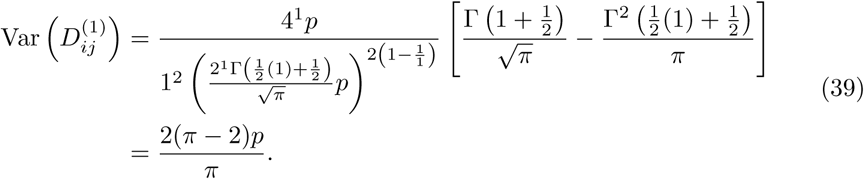

Similar to the mean (Eq. 38), the limiting variance of 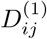 (Eq. 39) grows on the order of feature dimension *p*, which implies that points become more dispersed as the dimension increases. The moment estimates given in this section (Eqs. 38 and 39) are summarized in a table that is organized by metric, data type, statistic (mean or variance), and asymptotic formula (Tab. 2).

**Table 2.** Asymptotic estimates of means and variances for the standard *L*_1_ and *L*_2_ (*q* = 1 and *q* = 2 in Tab. 1) distance distributions. Estimates for both standard normal (𝒩 (0, 1)) and standard uniform (𝒰(0, 1)) data are given.

| $q$-Metric | Data | Stat | Formula (Eq. #) |
| --- | --- | --- | --- |
| standard<br>$L_1$ (Eq. 1) | $\mathcal{N}(0, 1)$ | mean | $\frac{2p}{\sqrt{\pi}} \quad (38)$ |
| | | variance | $\frac{2(\pi-2)p}{\pi} \quad (39)$ |
| | $\mathcal{U}(0, 1)$ | mean | $\frac{p}{3} \quad (40)$ |
| | | variance | $\frac{p}{18} \quad (41)$ |
| standard<br>$L_2$ (Eq. 1) | $\mathcal{N}(0, 1)$ | mean | $\sqrt{2p-1} \quad (48)$ |
| | | variance | $1 \quad (49)$ |
| | $\mathcal{U}(0, 1)$ | mean | $\sqrt{\frac{p}{6} - \frac{7}{120}} \quad (51)$ |
| | | variance | $\frac{7}{120} \quad (52)$ |

#### 2.2.2 Standard uniform data

Substituting *q* = 1 into the asymptotic formula of the mean (Eq. 37), we have the following for the expected *L*_1_ distance between two independently sampled instances *i, j* ∈ ℐ in standard uniform data

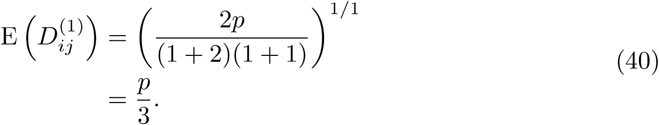

Once again, we see that the mean of 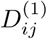 (Eq. 40) grows on the order of *p* just as in the case of standard normal data.

Substituting *q* = 1 into the formula of the asymptotic variance of 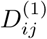 (Eq. 37) leads to the following

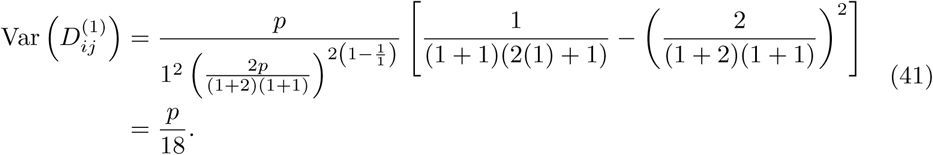

As in the case of the *L*_1_ metric on standard normal data, we have a variance (Eq. 41) that grows on the order of *p*. The distances between points in high-dimensional uniform data become more widely dispersed with this metric. The moment estimates given in this section (Eqs. 40 and 41) are summarized in a table that is organized by metric, data type, statistic (mean or variance), and asymptotic formula (Tab. 2).

#### 2.2.3 Distribution of one-dimensional projection of pairwise distance onto an attribute

In nearest-neighbor distance-based feature selection like NPDR and Relief-based algorithms, the one-dimensional projection of the pairwise distance onto an attribute (Eq. 2) is particularly fundamental to feature quality for association with an outcome. For instance, this distance projection is the predictor used to determine beta coefficients in NPDR. In particular, understanding distributional properties of the projected distances is necessary for defining pseudo P values for NPDR. In this section, we summarize the exact distribution of the one-dimensional projected distance onto an attribute *a* ∈ 𝒜. These results apply to continuous data, such as gene expression.

In previous sections, we derived the exact density function (Eq. 14) and moments (Eqs. 15 and 16) for the distribution of 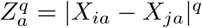. We then derived the exact density (Eq. 22) and moments (Eqs. 23 and 24) for standard normal data. Analogously, we formulated the exact density (Eq. 31) and moments (Eqs. 33 and 34) for standard uniform data. From these exact densities and moments, we simply substitute *q* = 1 to define the distribution of the one-dimensional projected distance onto an attribute *a* ∈ 𝒜.

Assuming data is standard normal, we substitute *q* = 1 into the density function of 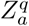 (Eq. 22) to arrive at the following density function

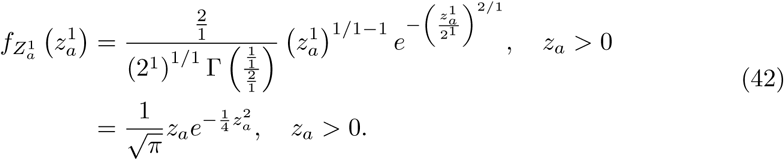

The mean corresponding to this Generalized Gamma density is computed by substituting *q* = 1 into the formula for the mean of 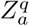 (Eq. 23). This result is given by

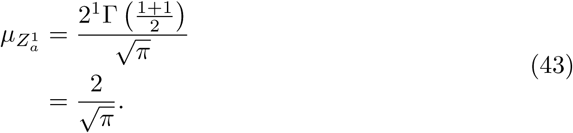

Substituting *q* = 1 into Eq. 24 for the variance, we have the following

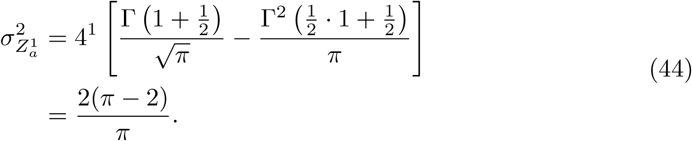

These last few results (Eqs. 42-44) provide us with the distribution for NPDR predictors when the data is from the standard normal distribution.

If we have standard uniform data, we substitute *q* = 1 into the density function of 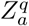 (Eq. 31) to obtain the following density function

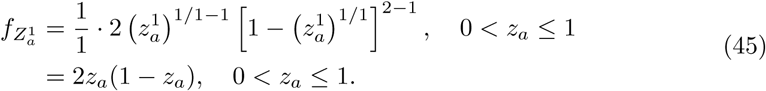

The mean corresponding to this Kumaraswamy density is computed by substituting *q* = 1 into the formula for the mean of 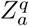 (Eq. 33). After substitution, we have the following result

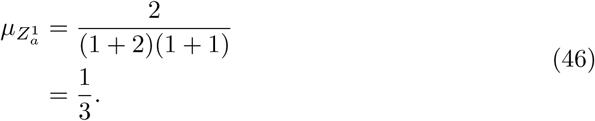

Substituting *q* = 1 into the formula for the variance of 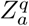 (Eq. 34), we have the following

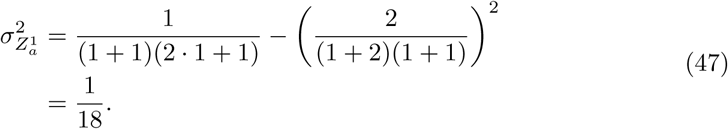

In the event that the data distribution is standard uniform, the density function (Eq. 45), the mean (Eq. 46, and the variance (Eq. 47) sufficiently define the distribution for NPDR predictors. The means (Eqs. 43 and 46) and variances (Eqs.44 and 47) come from the exact distribution of pairwise distances with respect to a single attribute *a* ∈ 𝒜. This is the distribution of the so-called “projection” of the pairwise distance onto a single attribute to which we have been referring, which is a direct implication from our more general derivations. In a similar manner, one can substitute any value of *q* ≥ 2 into the general densities of 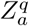 for standard normal (Eq. 22) and standard uniform (Eq. 31) to derive the associated density of 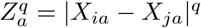 for the given data type.

### 2.3 Euclidean (*L*_2_)

Moment estimates for the Euclidean metric are obtained by substituting *q* = 2 into the asymptotic moment formulas for standard normal data (Eq. 27) and standard uniform data (Eq. 37). As in the case of the Manhattan metric in the previous sections, we initially proceed by deriving Euclidean distance moments in standard normal data.

#### 2.3.1 Standard normal data

Substituting *q* = 2 into the asymptotic formula of the mean (Eq. 27), we have the following for expected *L*_2_ distance between two independently sampled instances *i, j* ∈ ℐ in standard normal data

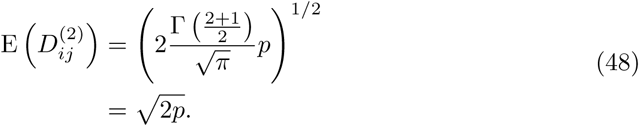

In the case of *L*_2_ on standard normal data, we see that the mean of 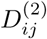 (Eq. 48) grows on the order of 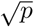. Hence, the Euclidean distance does not increase as quickly as the Manhattan distance on standard normal data.

Substituting *q* = 2 into the formula for the asymptotic variance of 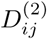 (Eq. 27) leads to the following

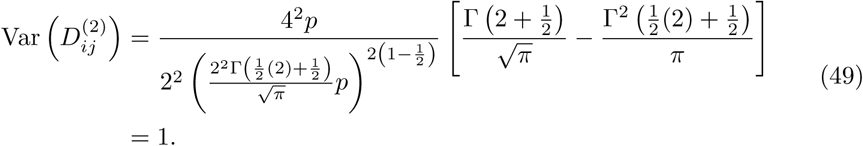

Surprisingly, the asymptotic variance (Eq. 49) is just 1. Regardless of data dimensions *m* and *p*, the variance of Euclidean distances on standard normal data tends to 1. Therefore, most instances are contained within a ball of radius 1 about the mean in high feature dimension *p*. This means that the Euclidean distance distribution on standard normal data is simply a horizontal shift to the right of the standard normal distribution.

For the case in which the number of attributes *p* is small, we have an improved estimate of the mean (Eq. 6). The lower dimensional estimate of the mean is given by

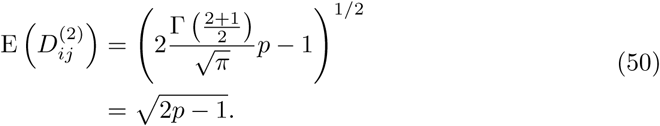

For high dimensional data sets like gene expression [16, 17], which typically contain thousands of genes (or features), it is clear that the magnitude of *p* will be sufficient to use the standard asymptotic estimate (Eq. 48) since 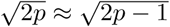 in that case. The moment estimates given in this section (Eqs. 50 and 49) are summarized in a table that is organized by metric, data type, statistic (mean or variance), and asymptotic formula (Tab. 2).

#### 2.3.2 Standard uniform data

Substituting *q* = 2 into the asymptotic formula of the mean (Eq. 37), we have the following for expected *L*_2_ distance between two independently sampled instances *i, j* ∈ ℐ in standard uniform data

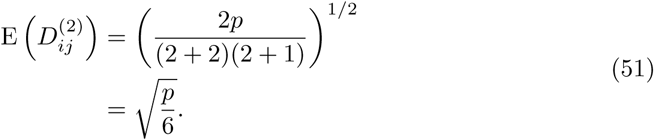

As in the case of standard normal data, the expected value of 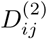 (Eq. 51) grows on the order of 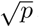.

Substituting *q* = 2 into the formula for the asymptotic variance of 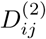 (Eq. 37) leads to the following

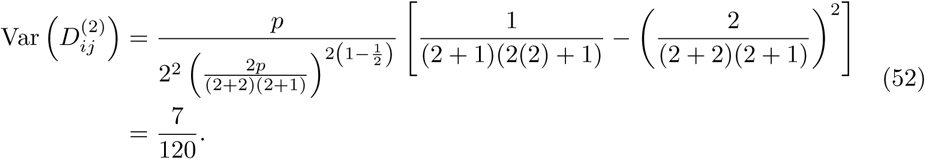

Once again, the variance of Euclidean distance surprisingly approaches a constant.

For the case in which the number of attributes *p* is small, we have an improved estimate of the mean (Eq. 6). The lower dimensional estimate of the mean is given by

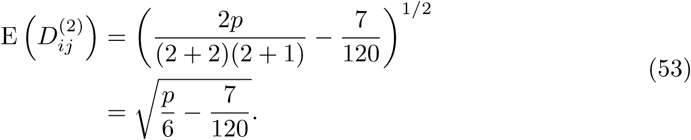

We summarize the moment estimates given in this section for standard *L*_*q*_ metrics (Eqs. 53 and 52) organized by metric, data type, statistic (mean or variance), and asymptotic formula (Table 2). In the next section, we extend these results for the standard *L*_*q*_ metric to derive asymptotics for the attribute range-normalized (max-min) *L*_*q*_ metric used frequently in Relief-based algorithms [1, 3] for scoring attributes. These derivations use extreme value theory to handle the maximum and minimum attributes for standard normal and standard uniform data.

### 2.4 Distribution of max-min normalized *L*_*q*_ metric

For Relief-based methods [1, 3], the standard numeric diff metric is given by

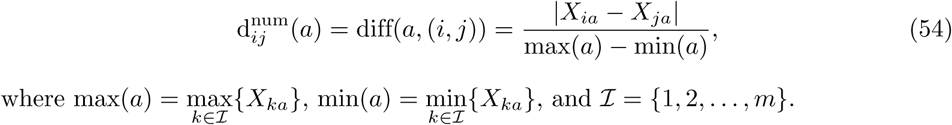

The pairwise distance using this max-min normalized diff metric is then computed as

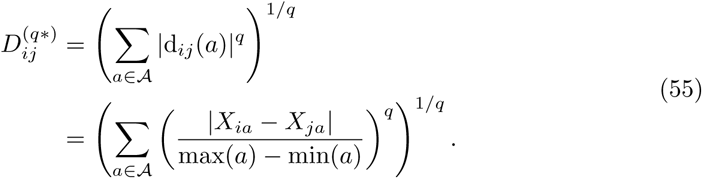

In order to determine moments of asymptotic max-min normalized distance (Eq. 54) distributions, we will first derive the asymptotic extreme value distributions of the attribute maximum and minimum. Although the exact distribution of the maximum or minimum requires an assumption about the data distribution, the Fisher-Tippett-Gnedenko Theorem is an important result that allows one to generally categorize the extreme value distribution for a collection of independent and identically distributed random variables into one of three distributional families. This theorem does not, however, tell us the exact distribution of the maximum that we require in order to determine asymptotic results for the max-min normalized distance (Eq. 55). We mention this theorem simply to provide some background on convergence of extreme values. Before stating the theorem, we first need the following definition

#### Definition 2.1

*A distribution* ℱ_*X*_ *is said to be* ***degenerate*** *if its density function f*_*X*_ *is the Dirac delta δ* (*x* − *c*_0_) *centered at a constant c*_0_ ∈ ℝ, *with corresponding distribution function F*_*X*_ *defined as*

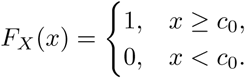

#### Theorem 2.2 (Fisher-Tippett-Gnedenko)

*Let* 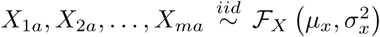 *and let* 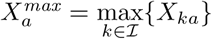. *If there exists two non-random sequences b*_*m*_ *>* 0 *and c*_*m*_ *such that*

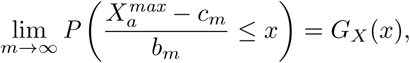

*where G*_*X*_ *is a non-degenerate distribution function, then the limiting distribution 𝒢*_*X*_ *is in the Gumbel, Fréchet, or Wiebull family*.

The three distribution families given in Theorem 2.2 are actually special cases of the Generalized Extreme Value Distribution. In the context of extreme values, Theorem 2.2 is analogous to the Central Limit Theorem for the distribution of sample mean. Although we will not explicitly invoke this theorem, it does tell us something very important about the asymptotic behavior of sample extremes under certain necessary conditions. For illustration of this general phenomenon of sample extremes, we derive the distribution of the maximum for standard normal data to show that the limiting distribution is in the Gumbel family, which is a well known result. In the case of standard uniform data, we will derive the distribution of the maximum and minimum directly. Regardless of data type, the distribution of the sample maximum can be derived as follows

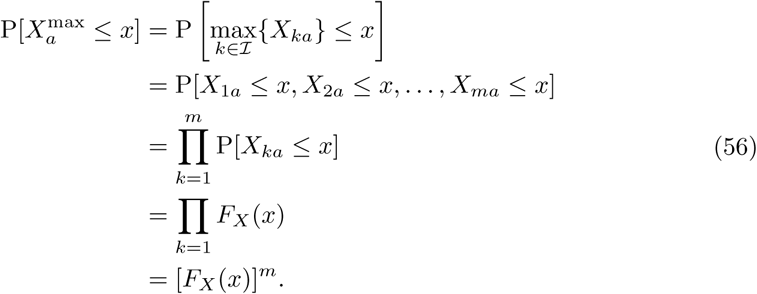

Using more precise notation, the distribution function of the sample maximum in standard normal data is

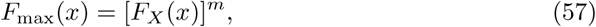

where *m* is the size of the sample from which the maximum is derived and *F*_*X*_ is the distribution function corresponding to the data sample. This means that the distribution of the sample maximum relies only on the distribution function of the data from which extremes are drawn *F*_*X*_ and the size of the sample *m*.

Differentiating the distribution function (Eq. 57) gives us the following density function for the distribution of the maximum

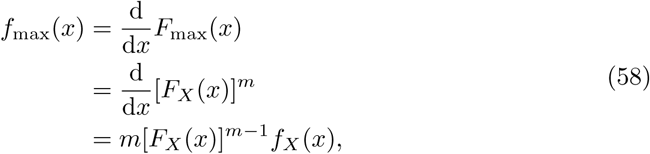

where *m* is the size of the sample from which the maximum is derived, *F*_*X*_ is the distribution function corresponding to the data sample, and *f*_*X*_ is the density function corresponding to the data sample. Similar to the distribution function for the sample maximum (Eq. 57), the density function (Eq 58) relies only on the distribution and density function of the data from which extremes are derived.

The distribution of the sample minimum, 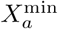, can be derived as follows

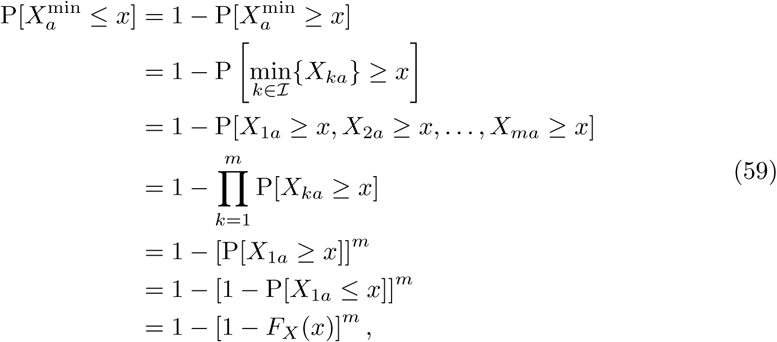

where *m* is the size of the sample from which the maximum is derived and *F*_*X*_ is the distribution function corresponding to the data sample. Therefore, the distribution of sample minimum also relies only on the distribution function of the data from which extremes are derived.

With more precise notation, we have the following expression for the distribution function of the minimum

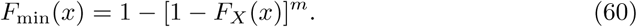

where *m* is the size of the sample from which the minimum is derived and *F*_*X*_ is the distribution function corresponding to the data sample.

Differentiating the distribution function (Eq. 60) gives us the following density function for the distribution of sample minimum

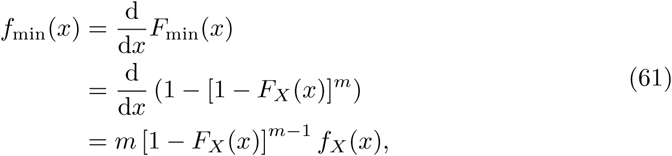

where *m* is the size of the sample from which the minimum is derived, *F*_*X*_ is the distribution function corresponding to the data sample, and *f*_*X*_ is the density function corresponding to the data sample. As in the case of the density function for sample maximum (Eq. 58), the density function for sample minimum relies only on the distribution *F*_*X*_ and density *f*_*X*_ functions of the data from which extremes are derived and the sample size *m*.

Given the densities of the distribution of sample maximum and minimum, we can easily compute the raw moments and variance. The first moment about the origin of the distribution of sample maximum is given by the following

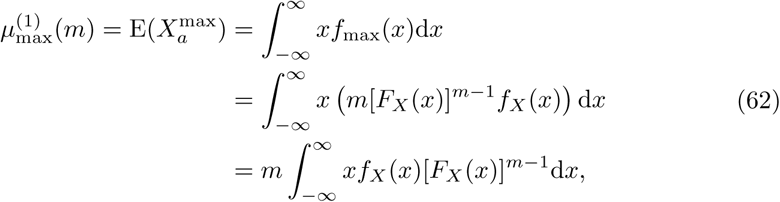

where *m* is the sample size, *F*_*X*_ is the distribution function, and *f*_*X*_ is the density function of the data from which the maximum is derived.

The second raw moment of the distribution of sample maximum is derived similarly as follows

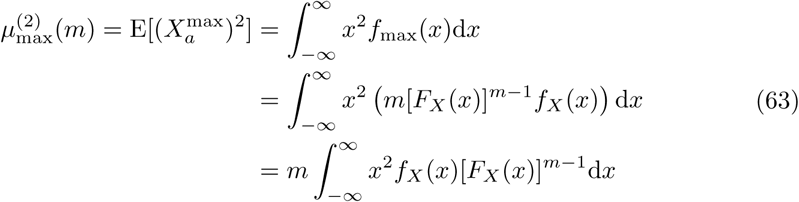

where *m* is the sample size, *F*_*X*_ is the distribution function, and *f*_*X*_ is the density function of the data from which the maximum is derived.

Using the first (Eq. 62) and second (Eq. 63) raw moments of the distribution of sample maximum, the variance is given by

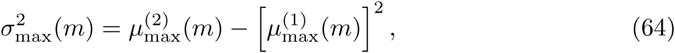

where *m* is the sample size of the data from which the maximum is derived and 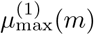 and 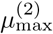 are the first and second raw moments, respectively, of the distribution of sample maximum.

Moving on to the distribution of sample minimum, the first raw moment is given by the following

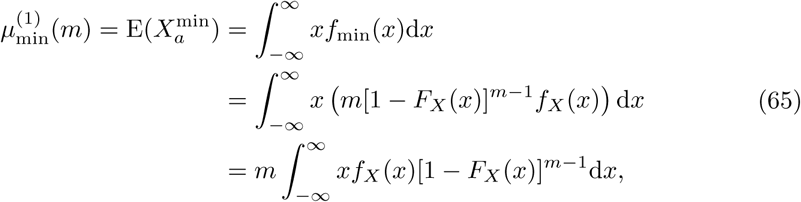

where *m* is the sample size, *F*_*X*_ is the distribution function, and *f*_*X*_ is the density function of the data from which the minimum is derived.

Similarly, the second raw moment of the distribution of sample minimum is given by the following

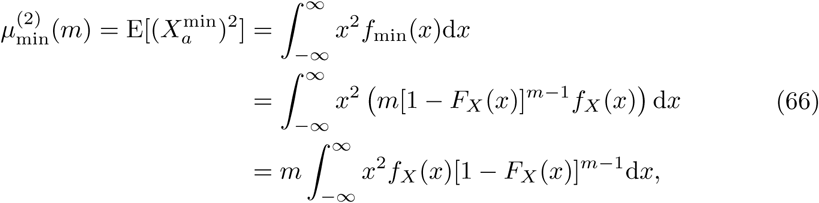

where *m* is the sample size, *F*_*X*_ is the distribution function, and *f*_*X*_ is the density function of the data from which the minimum is derived.

Using the first (Eq. 65) and second (Eq. 66) raw moments of the distribution of sample minimum, the variance is given by

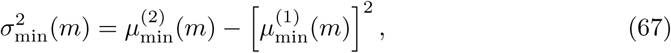

where *m* is the sample size of the data from which the maximum is derived and 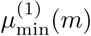 and 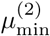 are the first and second raw moments, respectively, of the distribution of sample maximum.

Using the expected attribute maximum (Eq. 62) and minimum (Eq. 65) for sample size *m*, the following expected attribute range results from linearity of the expectation operator

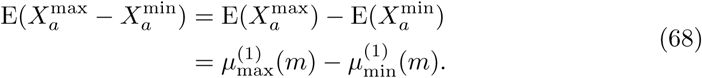

where 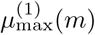 is the expected sample maximum (Eq. 62) and 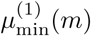 is the expected sample minimum.

For a data distribution whose density is an even function, the expected attribute range (Eq. 68) can be simplified to the following expression

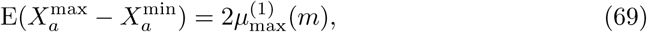

where *m* is the size of the sample from which the maximum is derived. Hence, the expected attribute range is simply twice the expected attribute maximum (Eq. 62). This result naturally applies to standard normal data, which is symmetric about its mean at 0 and without any skewness.

For large samples (*m >>* 1) from an exponential type distribution that has infinite support and all moments, the covariance between the sample maximum and minimum is approximately zero [18]. In this case, the variance of the attribute range of a sample of size *m* is given by the following

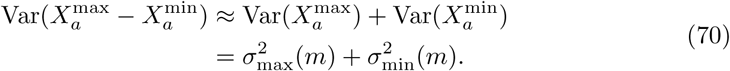

Under the assumption of zero skewness, infinite support and even density function, sufficiently large sample size *m*, and distribution of an exponential type for all moments, the variance of attribute range (Eq. 70) simplifies to the following

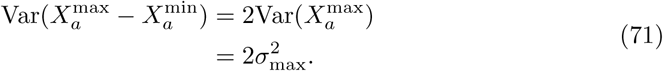

Let 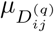 and 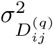 (Eq. 18) denote the mean and variance of the standard *L*_*q*_ distance metric (Eq. 1). Then the expected value of the max-min normalized distance (Eq. 55) distribution is given by the following

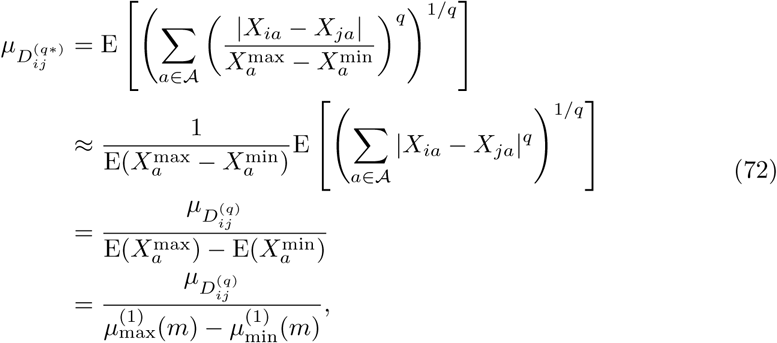

where *m* is the size of the sample from which extremes are derived, 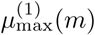 is the expected value of the sample maximum (Eq. 62), and 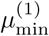 is the expected value of the sample minimum.

The variance of the max-min normalized distance (Eq. 55) distribution is given by the following

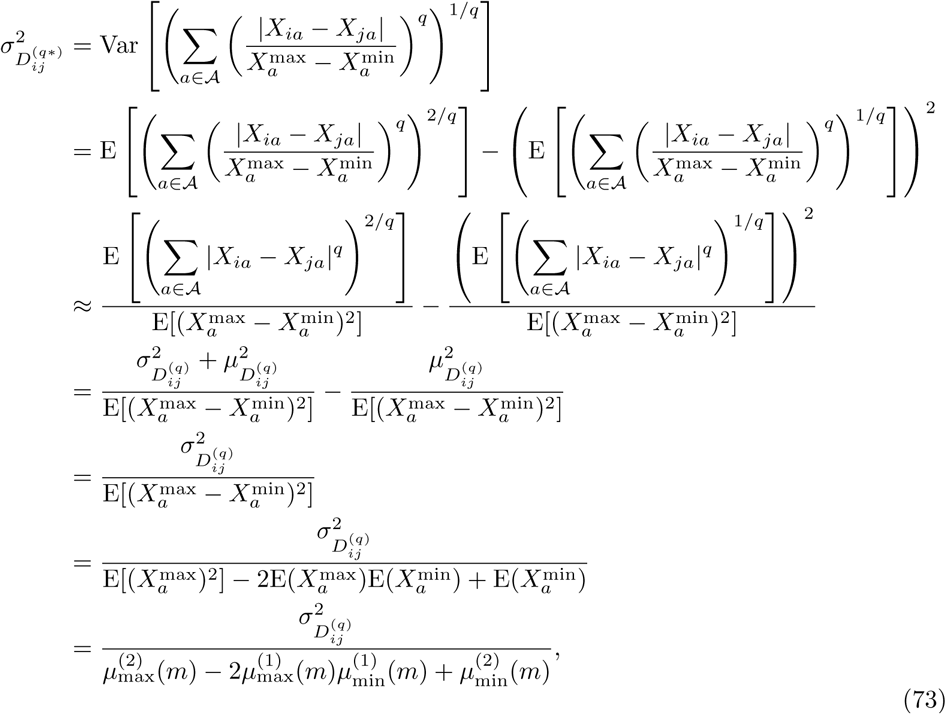

where *m* is the size of the sample from which extremes are derived, 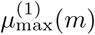 is the expected value of the sample maximum (Eq. 62), and 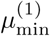 is the expected value of the sample minimum.

With the mean (Eq. 72) and variance (Eq. 73) of the max-min normalized distance (Eq. 55), we have the following generalized estimate for the asymptotic distribution of the max-min normalized distance distribution

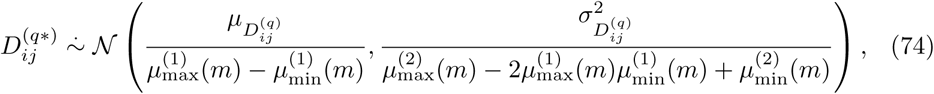

where *m* is the size of the sample from which extremes are derived, 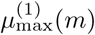 is the expected value of the sample maximum (Eq. 62), and 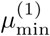 is the expected value of the sample minimum.

For data with zero skewness and support that is symmetric about 0, the expected sample maximum is the additive inverse of the expected sample minimum. This allows us to express the expected max-min normalized pairwise distance (Eq. 72) exclusively in terms of the expected sample maximum. This result is given by the following

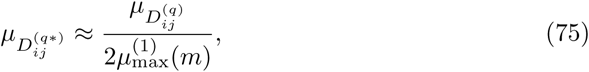

where *m* is the size of the sample from which the maximum is derived and 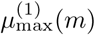 is the expected value of the sample maximum (Eq. 62).

A similar substitution gives us the following expression for the variance of the max-min normalized distance distribution

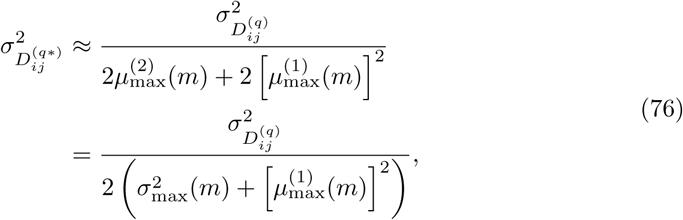

where *m* is the size of the sample from which extremes are derived, 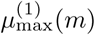 is the expected value of the sample maximum (Eq. 62), and 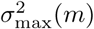 is the variance of the sample maximum (Eq. 64).

Therefore, the asymptotic distribution of the max-min normalized distance distribution (Eq. 74) becomes

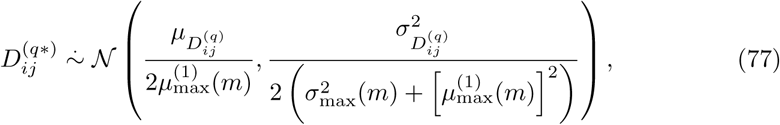

where *m* is the size of the sample from which extremes are derived, 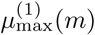 is the expected value of the sample maximum (Eq. 62), and 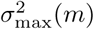 is the variance of the sample maximum (Eq. 64).

We have now derived asymptotic estimates of the moments of the max-min normalized *L*_*q*_ distance metric (Eq. 55) for any continuous data distribution. In the next two sections, we examine the max-min normalized *L*_*q*_ distance on standard normal and standard uniform data. As in previous sections in which we analyzed the standard *L*_*q*_ metric (Eq. 1), we will use the more general results for the max-min *L*_*q*_ metric to derive asymptotic estimates for normalized Manhattan (*q* = 1) and Euclidean (*q* = 2).

#### 2.4.1 Standard normal data

The standard normal distribution has zero skewness, even density function, infinite support, and all moments. This implies that the corresponding mean and variance of the distribution of sample range can be expressed exclusively in terms of the sample maximum. Given the nature of the density function of the sample maximum for sample size *m*, the integration required to determine the moments (Eqs. 62 and 63) is not possible. These moments can either be approximated numerically or we can use extreme value theory to determine the form of the asymptotic distribution of the sample maximum. Using the latter method, we will show that the asymptotic distribution of the sample maximum for standard normal data is in the Gumbel family. Let 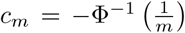 and 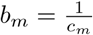, where **Φ** is the standard normal cumulative distribution function. Using Taylor’s Theorem, we have the following expansion

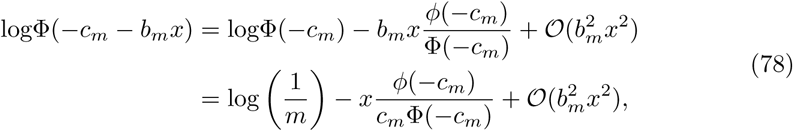

where *m* is the size of the sample from which the maximum is derived.

In order to simplify the right-hand side of this expansion (Eq. 78), we will use the well known Mills Ratio Bounds [19] given by the following

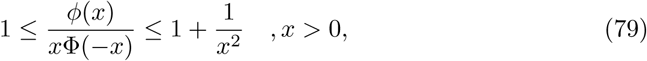

where **Φ** and *ϕ* once again represent the cumulative distribution function and density function, respecively, of the standard normal distribution.

The inequalities given above (Eq. 79) show that

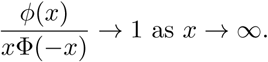

This further implies that

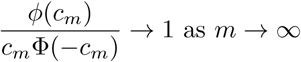

since

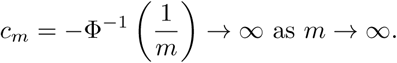

This gives us the following approximation of the right-hand side of the expansion (Eq. 78) given previously

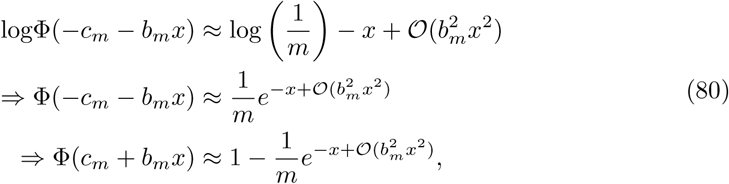

where *m* is the size of the sample from which the maximum is derived.

Using the approximation of expansion given previously (Eq. 80), we now derive the limit distribution for the sample maximum in standard normal data as

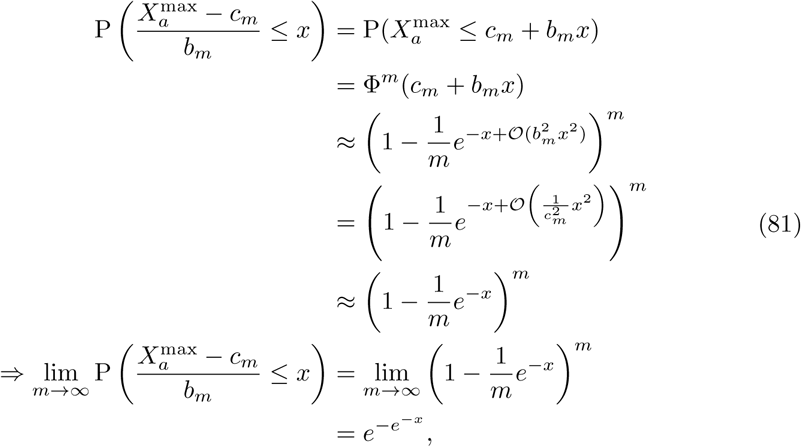

which is the cumulative distribution function of the standard Gumbel distribution. The mean of this distribution is given by the following

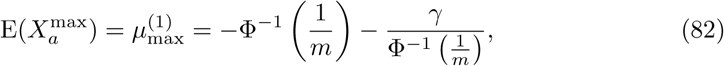

where *m* is the size of the sample from which the maximum is derived and *γ* is the well known Euler-Mascheroni constant. This constant has many equivalent definitions, one of which is given by

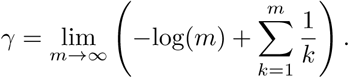

Perhaps a more convenient definition of the Euler-Mascheroni constant is simply

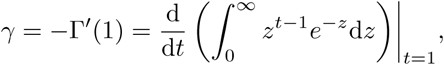

which is just the additive inverse of the first derivative of the gamma function evaluated at 1.

The median of the distribution of the maximum for standard normal data is given by

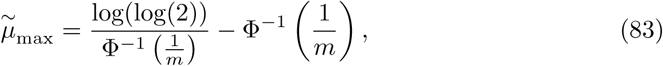

where *m* is the size of the sample from which the maximum is derived.

Finally, the variance of the asymptotic distribution of the sample maximum is given by

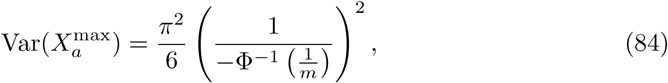

where *m* is the size of the sample from which the maximum is derived.

For typical sample sizes *m* in high-dimensional spaces, the variance estimate (Eq. 84) exceeds the variance of the sample maximum significantly. Using the fact that

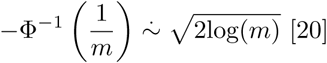

and

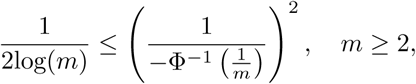

we can get a more accurate approximation of the variance with the following

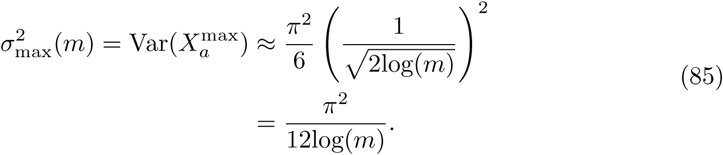

Therefore, the mean of the range of *m* iid standard normal random variables is given by

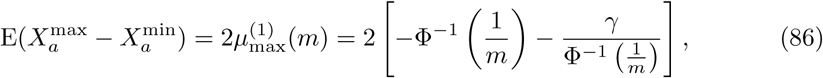

where *γ* is the Euler-Mascheroni constant.

It is well known that the sample extremes from the standard normal distribution are approximately uncorrelated for large sample size *m* [18]. This implies that we can approximate the variance of the range of *m* iid standard normal random variables with the following result

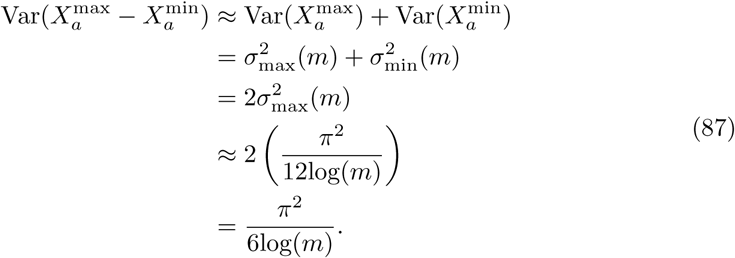

For the purpose of approximating the mean and variance of the max-min normalized distance distribution, we observe empirically that the formula for the median of the distribution of the attribute maximum (Eq. 83) yields more accurate results. More precisely, the approximation of the expected maximum (Eq. 82) overestimates the sample maximum slightly. The formula for the median of the sample maximum (Eq. 83) provides a more accurate estimate of this sample extreme. Therefore, the following estimate for the mean of the attribute range will be used instead

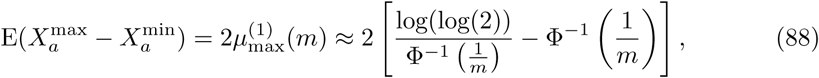

where *m* is the size of the sample from which extremes are derived.

We have already determined the mean and variance (Eq. 27) for the *L*_*q*_ metric (Eq. 1) on standard normal data. Using the expected value of the sample maximum (Eq. 88), the variance of the sample maximum (Eq. 87), and the general formulas for the mean and variance of the max-min normalized distance distribution (Eq. 77), this leads us to the following asymptotic estimate for the distribution of the max-min normalized distances for standard normal data

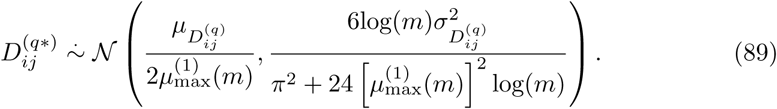

where *m* is the size of the sample from which the maximum is derived, 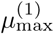 is the median of the sample maximum (Eq. 83), 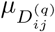 is the expected *L*_*q*_ pairwise distance (Eq. 25), and 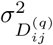 is the variance of the *L*_*q*_ pairwise distance (Eq. 26). The moments of the max-min normalized *L*_*q*_ distance metric in standard normal data (Eq. 89) are summarized in a table that is organized by metric, data type, statistic (mean or variance), and asymptotic formula (Table 1).

#### 2.4.2 Standard uniform data

Standard uniform data does not have an even density function. Due to the simplicity of the density function, however, we can derive the distribution of the maximum and minimum of a sample of size *m* explicitly. Using the general forms of the distribution functions of the maximum (Eq. 57) and minimum (Eq. 60), we have the following distribution functions for standard uniform data

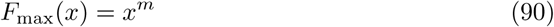

and

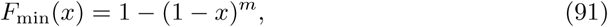

where *m* is the size of the sample from which extremes are derived.

Using the general forms of the density functions of the maximum (Eq. 58) and minimum (Eq. 61), we have the following density functions for standard uniform data

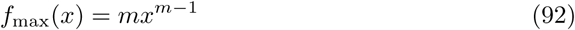

and

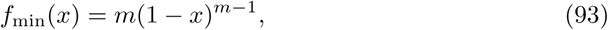

where *m* is the size of the sample from which extremes are derived.

Then the expected maximum and minimum are computed through straightforward integration as follows

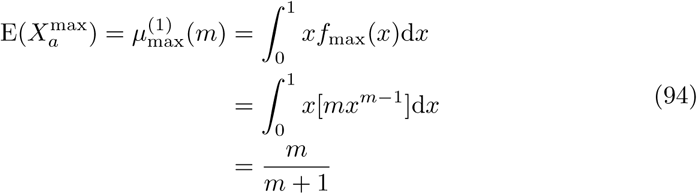

and

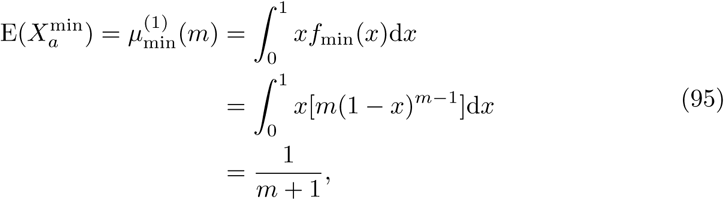

where *m* is the size of the sample from which extremes are derived.

We can compute the second moment about the origin of the sample range as follows

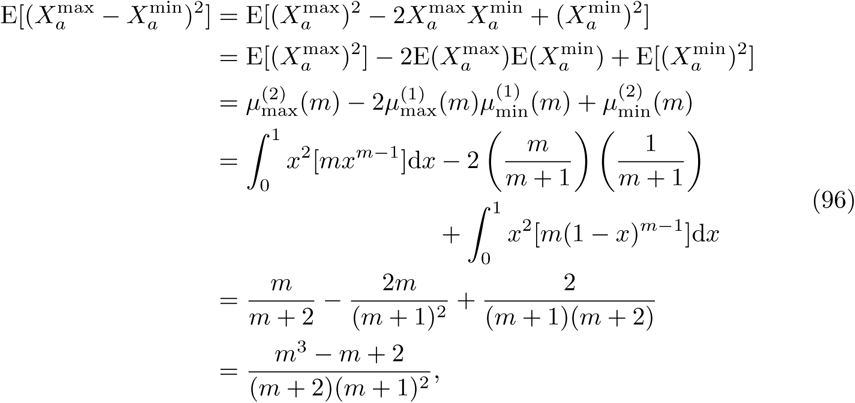

where *m* is the size of the sample from which extremes are derived.

Using the general asymptotic distribution of max-min normalized distances for any data type (Eq. 74) and the mean and variance (Eq. 37) of the standard *L*_*q*_ distance metric (Eq. 1), we have the following asymptotic estimate for the max-min normalized distance distribution for standard uniform data

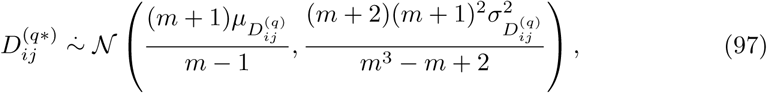

where *m* is the size of the sample from which extremes are derived, 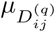 is the expected value (Eq. 35) of the *L*_*q*_ metric (Eq. 1) in standard uniform data, and 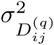 is the variance (Eq. 36) of the *L*_*q*_ metric (Eq. 1) in standard uniform data. The moments of the max-min normalized *L*_*q*_ distance metric in standard uniform data (Eq. 89) are summarized in a table that is organized by metric, data type, statistic (mean or variance), and asymptotic formula (Tab. 1).

### 2.5 Range-Normalized Manhattan (*q* = 1)

Using the general asymptotic results for mean and variance of max-min normalized distances in standard normal and standard uniform data (Eqs. 89 and 97) for any value of *q* ∈ ℕ, we can substitute a particular value of *q* in order to determine a more specified distribution for the normalized distance (*D*^(*q*\*)^, Eq. 55). The following results are for the max-min normalized Manhattan (*q* = 1), *D*^(1*)^, metric for both standard normal and standard uniform data.

#### 2.5.1 Standard normal data

Substituting *q* = 1 into the asymptotic formula for the expected max-min normalized distance (Eq. 89), we derive the expected normalized Manhattan distance in standard normal data as follows

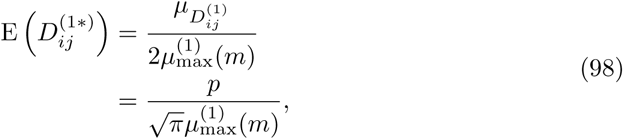

where 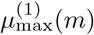 is the expected attribute maximum (Eq. 83), *m* is the size of the sample from which the maximum is derived, and *p* is the total number of attributes.

Similarly, the variance of 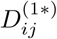 is given by

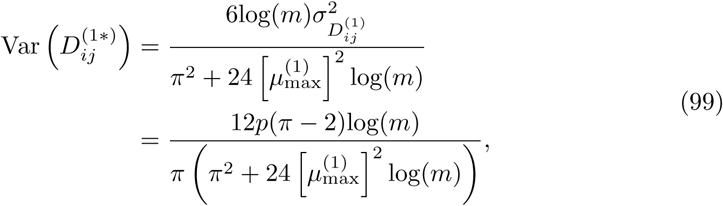

where 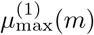 is the expected attribute maximum (Eq. 83), *m* is the size of the sample from which the maximum is derived, and *p* is the total number of attributes. Similar to the variance of the standard Manhattan distance, the variance of the max-min normalized Manhattan distance is on the order of *p* for fixed instance dimension *m*. For fixed *p*, the variance (Eq. 99) vanishes as *m* grows without bound. If we fix *m*, the same variance increases monotonically with increasing *p*. The moments derived in this section (Eqs. 98 and 99) are summarized in a table that is organized by metric, data type, statistic (mean or variance), and asymptotic formula (Table 3).

**Table 3.** Asymptotic estimates of means and variances for the max-min normalized *L*_1_ and *L*_2_ distance distributions commonly used in Relief-based algorithms. Estimates for both standard normal (𝒩(0, 1)) and standard uniform (𝒰(0, 1)) data are given. The cumulative distribution function of the standard normal distribution is represented by **Φ**. Furthermore, 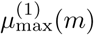 (Eq. 83) is the asymptotic median of the sample maximum from *m* standard normal random samples.

| **$q$-Metric** | **Data** | **Stat** | **Formula (Eq. #)** |
| --- | --- | --- | --- |
| max-min<br>normalized<br>$L_1$ (Eq. 55) | $\mathcal{N}(0, 1)$ | mean | $\frac{p}{\sqrt{\pi}\mu_{\max}^{(1)}(m)} \quad (98)$<br>where $\mu_{\max}^{(1)}(m) = \frac{\log(\log(2))}{\Phi^{-1}\left(\frac{1}{m}\right)} - \Phi^{-1}\left(\frac{1}{m}\right)$ |
| | | variance | $\frac{12p(\pi-2)\log(m)}{\pi\left(\pi^2+24\left[\mu_{\max}^{(1)}(m)\right]^2\log(m)\right)} \quad (99)$<br>where $\mu_{\max}^{(1)}(m) = \frac{\log(\log(2))}{\Phi^{-1}\left(\frac{1}{m}\right)} - \Phi^{-1}\left(\frac{1}{m}\right)$ |
| | $\mathcal{U}(0, 1)$ | mean | $\frac{(m+1)p}{3(m-1)} \quad (100)$ |
| | | variance | $\frac{(m+2)(m+1)^2p}{18(m^3-m+2)} \quad (100)$ |
| max-min<br>normalized<br>$L_2$ (Eq. 55) | $\mathcal{N}(0, 1)$ | mean | $\frac{\sqrt{2p-1}}{2\mu_{\max}^{(1)}(m)} \quad (102)$<br>where $\mu_{\max}^{(1)}(m) = \frac{\log(\log(2))}{\Phi^{-1}\left(\frac{1}{m}\right)} - \Phi^{-1}\left(\frac{1}{m}\right)$ |
| | | variance | $\frac{6\log(m)}{\pi^2+24\left[\mu_{\max}^{(1)}(m)\right]^2\log(m)} \quad (103)$<br>where $\mu_{\max}^{(1)}(m) = \frac{\log(\log(2))}{\Phi^{-1}\left(\frac{1}{m}\right)} - \Phi^{-1}\left(\frac{1}{m}\right)$ |
| | $\mathcal{U}(0, 1)$ | mean | $\sqrt{\frac{p}{6} - \frac{7}{120} \left(\frac{m+1}{m-1}\right)} \quad (104)$ |
| | | variance | $\frac{7(m+2)(m+1)^2}{120(m^3-m+2)} \quad (105)$ |

#### 2.5.2 Standard uniform data

Substituting *q* = 1 into the asymptotic formula for the expected max-min pairwise distance (Eq. 97), we derive the expected normalized Manhattan distance in standard uniform data as

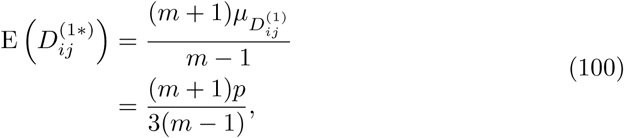

where *m* is the size of the sample from which extremes are derived and *p* is the total number attributes.

Similarly, the variance of 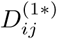 is given by

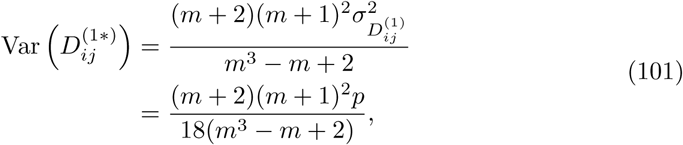

where *m* is the size of the sample from which extremes are derived and *p* is the total number of attributes. Interestingly, the variance of the max-min normalized Manhattan distance in standard uniform data approaches *p/*18 as *m* increases without bound for a fixed number of attributes *p*. This is the same asymptotic value to which the variance of the standard Manhattan distance (Eq. 40) converges. Therefore, large sample sizes make the variance of the normalized Manhattan distance approach the variance of the standard Manhattan distance in standard uniform data. The moments derived in this section (Eqs. 100 and 101) are summarized in a table that is organized by metric, data type, statistic (mean or variance), and asymptotic formula (Table 3).

### 2.6 Range-Normalized Euclidean (*q* = 2)

Analogous to the previous section, we use the asymptotic moment estimates for the max-min normalized metric (*D*^(*q*\*)^, Eq. 55) for standard normal (Eq. 89) and standard uniform (Eq. 97) data but specific to a range-normalized Euclidean metric (*q* = 2).

#### 2.6.1 Standard normal data

Substituting *q* = 2 into the asymptotic formula for the expected max-min normalized pairwise distance (Eq. 89), we derive the expected normalized Euclidean distance in standard normal data as

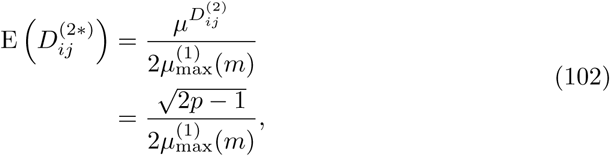

where 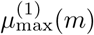 is the expected attribute maximum (Eq. 83), *m* is the size of the sample from which the maximum is derived, and *p* is the total number of attributes.

Similarly, the variance of 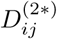 is given by

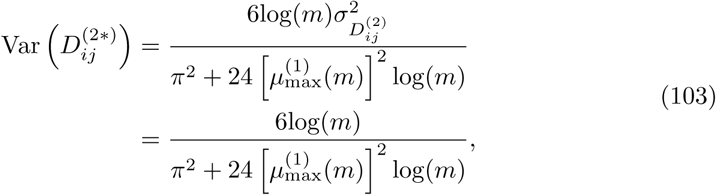

where 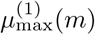 is the expected attribute maximum (Eq. 83) and *m* is the size of the sample from which the maximum is derived. It is interesting to note that the variance (Eq. 103) vanishes as the sample size *m* increases without bound, which means that all distances will be tightly clustered about the mean (Eq. 102). This is different than the variance of the standard *L*_2_ metric (Eq. 49), which is asymptotically equal to 1. This could imply that any two pairwise distances computed with the max-min normalized Euclidean metric in a large sample space *m* may be indistinguishable, which is another curse of dimensionality. The moments derived in this section (Eqs. 102 and 103) are summarized in a table that is organized by metric, data type, statistic (mean or variance), and asymptotic formula (Table 3).

#### 2.6.2 Standard uniform data

Substituting *q* = 2 into the asymptotic formula for the expected max-min normalized pairwise distance (Eq. 97), we derive the expected normalized Euclidean distance in standard uniform data as

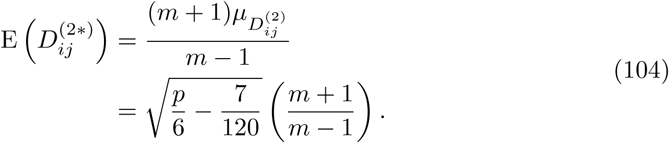

where *m* is the size of the sample from which extremes are derived and *p* is the total number of attributes.

Similarly, the variance of 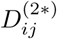 is given by

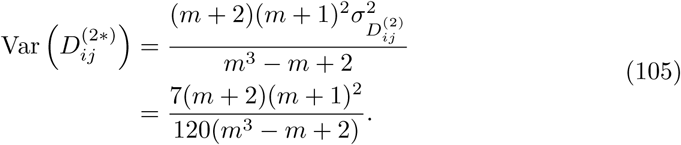

where *m* is the size of the sample from which extremes are derived. Similar to the variance of max-min normalized Manhattan distances in standard uniform data (Eq. 101), the variance of normalized Euclidean distances approaches the variance of the standard Euclidean distances in uniform data (Eq. 52) as *m* increases without bound. That is, the variance of the max-min normalized Euclidean distance (Eq. 105) approaches 7*/*120 as *m* grows larger. The moments derived in this section (Eqs. 104 and 105) are summarized in a table that is organized by metric, data type, statistic (mean or variance), and asymptotic formula (Table 3).

We summarize moment estimates in tables (Tables 1-3) that contain all of our asymptotic results for both standard and max-min normalized *L*_*q*_ metrics in each data type we have considered. This includes our most general results for any combination of sample size *m*, number of attributes *p*, type of metric *L*_*q*_, and data type (Table 1). From these more general derivations, we show the results of the standard *L*_1_ and *L*_2_ metrics for any combination of sample size *m*, number of attributes *p*, and data type (Table 2). Our last set of summarized results show asymptotics for the max-min normalized *L*_1_ and *L*_2_ metrics for any combination of sample size *m*, number of attributes *p*, and data type (Table 3). For both standard and max-min normalized *L*_2_ metrics (Tables 2 and 3), the low-dimensional improved estimates of sample means (Eqs. 50 and 53) are used because they perform well at both low and high attribute dimension *p*.

In the next section, we make a transition into discrete GWAS data. We will discuss some commonly known metrics and then a relatively new metric, which will lead us into novel asymptotic results for this data type.

## 3 GWAS distance distributions

In genome-wide association studies (GWAS), data is encoded by the minor allele at a particular locus *a*, which is just the second most common allele (adenine-A, thymine-T, cytosine-C, or guanine-G) associated with a given attribute *a* in the data set. Attributes in GWAS data are single nucleotide polymorphisms (SNPs), or mutations involving the substitution, deletion, or insertion of one nucleotide at some point in the DNA sequence of an organism. These are common mutations that can affect how an individual reacts to certain pathogens or the susceptibility for certain diseases. Feature selection in GWAS is typically concerned with finding interacting SNPs that are associated with disease susceptibility [21].

Consider a GWAS data set, which has the following encoding based on minor allele frequency

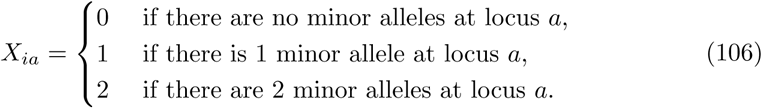

A minor allele at a particular locus *a* is the least frequent of the two alleles at that particular locus *a*. For random GWAS data sets, we can think *X*_*ia*_ as the number of successes in two Bernoulli trials. That is, *X*_*ia*_ ∼ ℬ(2, *f*_*a*_) where *f*_*a*_ is the probability of success. The success probability *f*_*a*_ is the probability of a minor allele occurring at *a*. Furthermore, the minor allele probabilities are assumed to be independent and identically distributed according to 𝒰(*l, u*), where *l* and *u* are the lower and upper bounds, respectively, of the sampling distribution’s support. Two commonly known types of distance metrics for GWAS data are the Genotype Mismatch (GM) and Allele Mismatch (AM) metrics. The GM and AM metrics are defined by

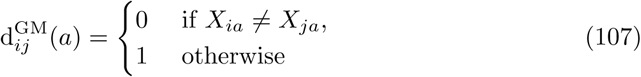

and

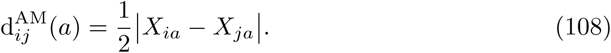

More informative metrics may include differences at the nucleotide level for each allele by considering differences in the rates of transition and transversion mutations (Fig. 2). One such discrete metric that accounts for transitions (Ti) and transversions (Tv) was introduced in [5] and can be written as

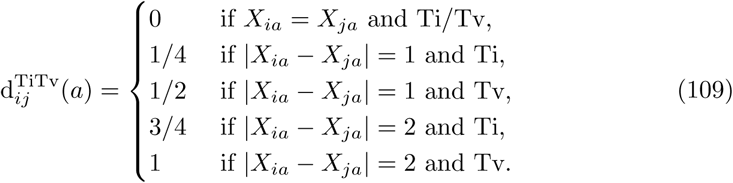

With these GWAS distance metrics, we then compute the pairwise distance between two instances *i, j* ∈ ℐ with

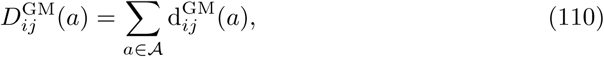

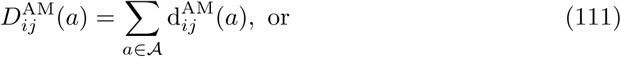

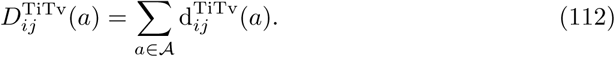

**Fig 2.**
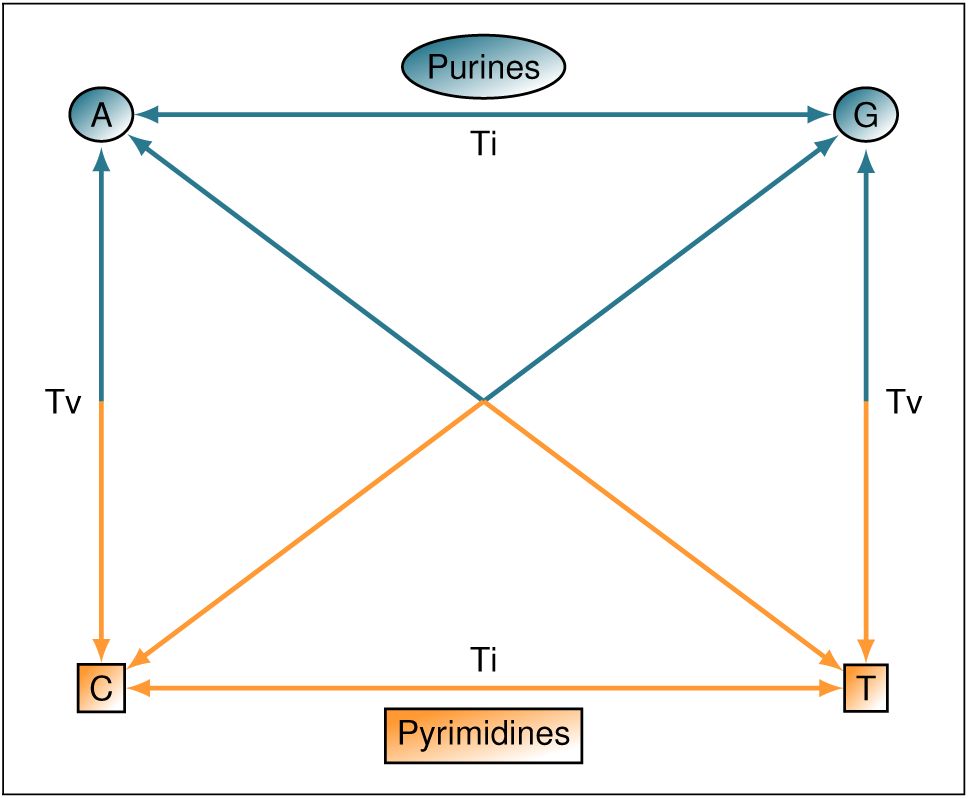
Purines (A and G) and pyrimidines (C and T) are shown. Transitions occur when a mutation involves purine-to-purine or pyrimidine-to-pyrimidine insertion. Transversions occur when a purine-to-pyrimidine or pyrimidine-to-purine insertion happens, which is a more extreme case. There are visibly more possibilities for transversions to occur than there are transitions, but there are about twice as many transitions in real data.

Assuming that all data entries *X*_*ia*_ are independent and identically distributed, we have already shown that the distribution of pairwise distances is asymptotically normal regardless of data distribution and value of *q*. Therefore, the distance distributions induced by each of the GWAS metrics (Eqs. 107-109) are asymptotically normal. With this Gaussian limiting behavior, we will proceed by deriving the mean and variance for each distance distribution induced by these three GWAS metrics.

### 3.1 GM distance distribution

The simplest distance metric in nearest-neighbor feature selection in GWAS data is the genotype-mismatch (GM) distance metric (Eq. 110). The GM attribute diff (Eq. 107) indicates only whether two genotypes are the same or not. There are many ways two genotypes could differ, but this metric does not record this information. We will now derive the moments for the GM distance (Eq. 110), which are sufficient for defining its corresponding asymptotic distribution.

The expected value of the GM attribute diff metric (Eq. 107) is given by the following

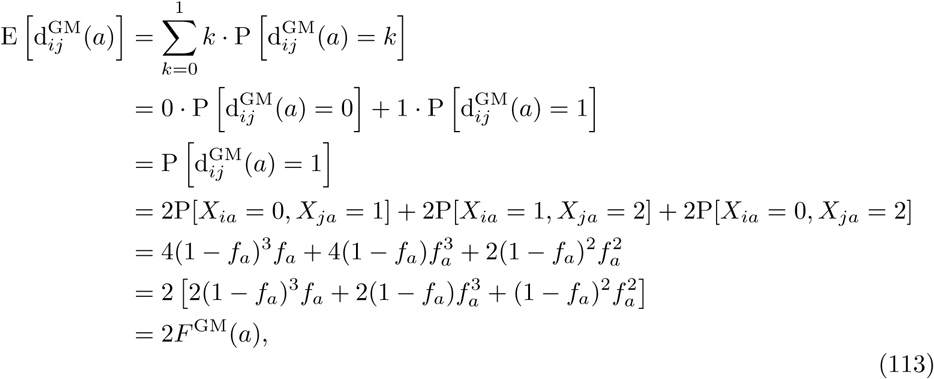

where 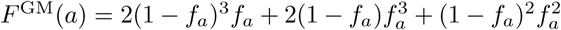 and *f*_*a*_ is the probability of a minor allele occurring at locus *a*.

Then the expected pairwise GM distance between instances *i, j* ∈ ℐ is given by

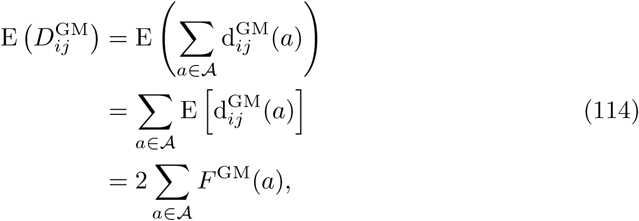

where 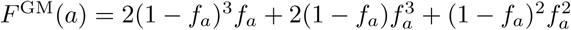 and *f*_*a*_ is the probability of a minor allele occurring at locus *a*. We see that the expected GM pairwise distance (Eq. 114) relies only on the minor allele probabilities *f*_*a*_ for all *a* ∈ 𝒜. In real data, we can easily determine these probabilities by dividing the total number of minor alleles at locus *a* by the twice the number of instances *m*. To be more explicit, this is just

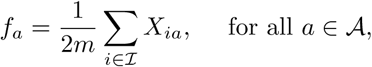

where *m* is the number of instances (or sample size). This is because each instance has two alleles, the minor and major alleles, at each locus. Therefore, the total number of alleles at locus *a* is 2*m*.

The second moment about the origin for the GM distance is computed as follows

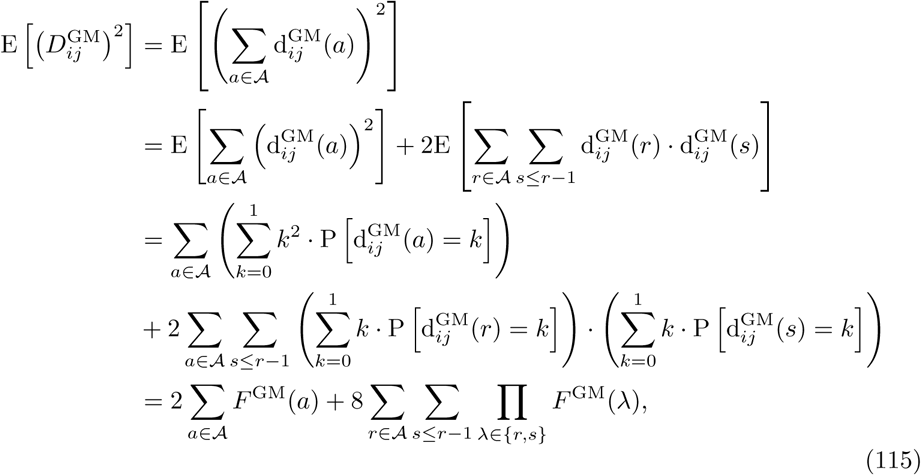

where 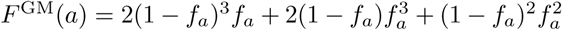 and *f*_*a*_ is the probability of a minor allele occurring at locus *a*.

Using the first (Eq. 114) and second (Eq. 115) raw moments of the GM distance, the variance is given by

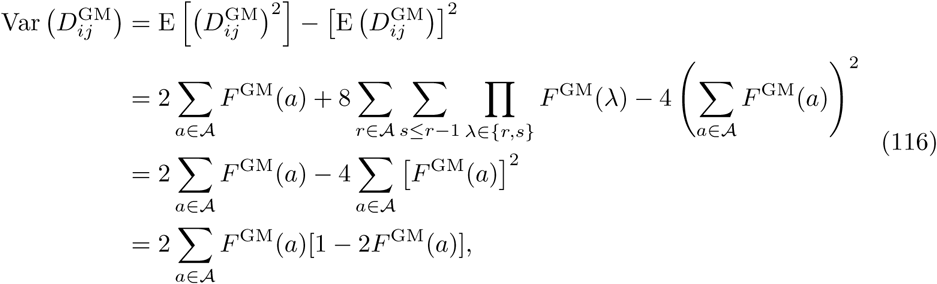

where 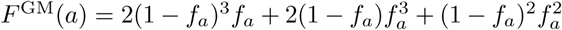 and *f*_*a*_ is the probability of a minor allele occurring at locus *a*. Hence, the variance of the asymptotic GM distance distribution also just depends on the minor allele probabilities *f*_*a*_ for all *a* ∈ 𝒜. This implies that the limiting GM distance distribution is fully determined by the minor allele probabilities, which are known in real data.

With the mean and variance estimates (Eqs. 114 and 116), the asymptotic GM distance distribution is given by the following

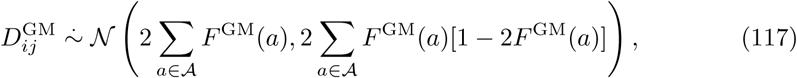

where 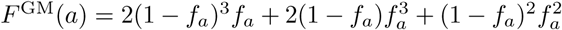 and *f*_*a*_ is the probability of a minor allele occurring at locus *a*. This GM distribution holds for random independent GWAS data with minor allele probabilities *f*_*a*_ and binomial samples *X*_*ia*_ ∼ ℬ (2, *f*_*a*_) for all *a* ∈ 𝒜. Next we consider the distance distribution for an AM metric, which incorporates differences at the allele level and contains more information than genotype differences.

### 3.2 AM distance distribution

As we have mentioned previously, the AM attribute diff metric (Eq. 108) is slightly more dynamic than the GM metric because the AM metric accounts for differences between the alleles of two genotypes. In this section, we derive moments of the AM distance metric (Eq. 111) that adequately define its corresponding asymptotic distribution.

The expected value of the AM attribute diff metric (Eq. 108) is given by the following

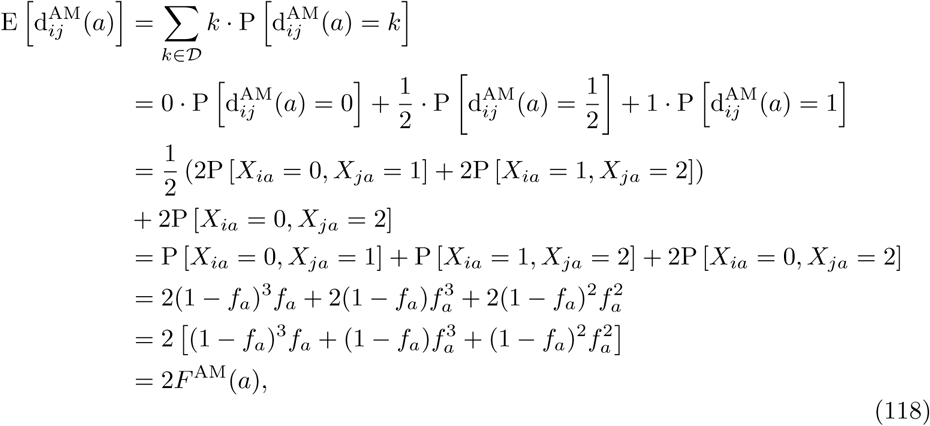

where 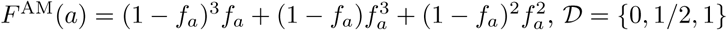 and *f*_*a*_ is the probability of a minor allele occurring at locus *a*.

Using the expected AM attribute diff (Eq. 118), the expected pairwise AM distance (Eq. 111) between instances *i, j* ∈ ℐ is given by

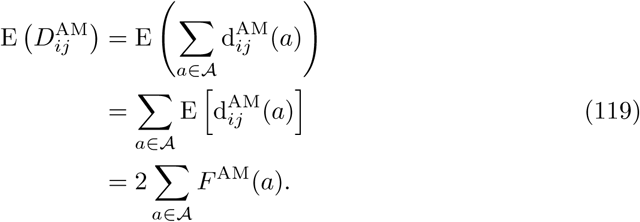

where 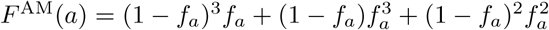 and *f*_*a*_ is the probability of a minor allele occurring at locus *a*. Similar to GM distances, the expected AM distance (Eq. 119) depends only on the minor allele probabilities *f*_*a*_ for all *a* ∈ 𝒜. This is to be expected because, although the AM metric is more informative, it still only accounts for simple differences between nucleotides of two instances *i, j* ∈ ℐ at some locus *a*.

The second moment about the origin for the AM distance is computed as follows

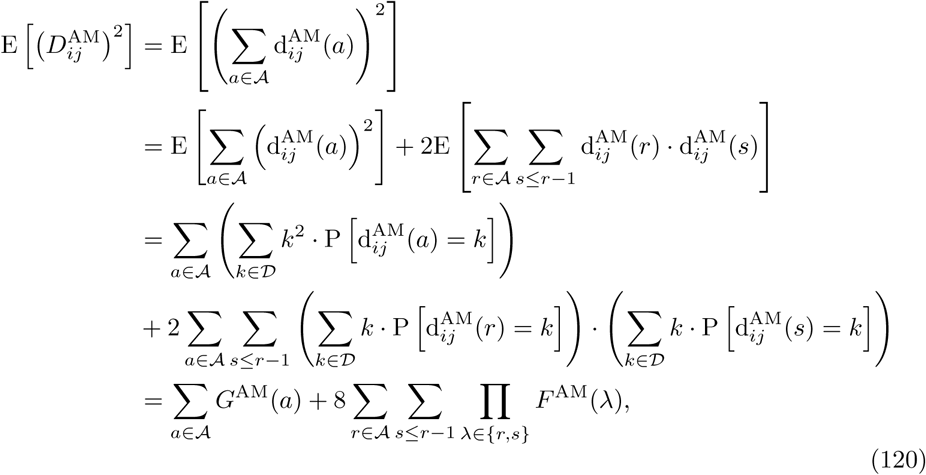

where 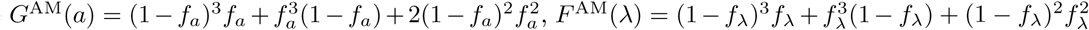, and *f*_*a*_ is the probability of a minor allele occurring at locus *a*.

Using the first (Eq. 119) and second (Eq. 120) raw moments of the asymptotic AM distance distribution, the variance is given by

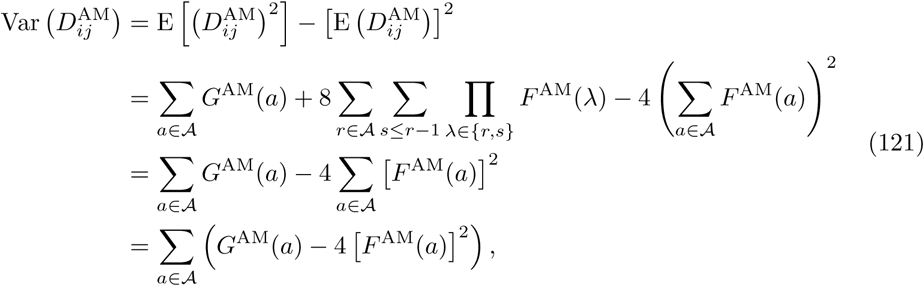

where 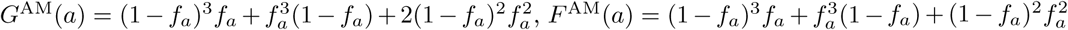, and *f*_*a*_ is the probability of a minor allele occurring at locus *a*. Similar to the mean (Eq. 119), the variance just depends on minor allele probabilities *f*_*a*_ for all *a* ∈ 𝒜.

With the mean (Eq. 119) and variance (Eq. 121) estimates of AM distances, the asymptotic AM distance distribution is given by the following

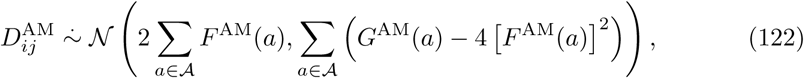

where 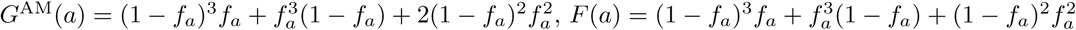, and *f*_*a*_ is the probability of a minor allele occurring at locus *a*.

This concludes our analysis of the AM metric in GWAS data when the independence assumption holds for minor allele probabilities *f*_*a*_ and binomial samples ℬ (2, *f*_*a*_) for all *a* ∈ 𝒜. In the next section, we derive more complex asymptotic results for the TiTv distance metric (Eq. 112).

### 3.3 TiTv distance distribution

The TiTv metric allows for one to account for both genotype mismatch, allele mismatch, transition, and transversion. However, this added dimension of information requires knowledge of the nucleotide makeup at a particular locus. A sufficient condition to compute the TiTv metric between instances *i, j* ∈ ℐ is that we know whether the nucleotides associated with a particular locus *a* are both purines (PuPu), purine and pyrimidine (PuPy), or both pyrimidines (PyPy). We illustrate all possibilities for transitions and transversions in a diagram (Fig. 2). Purines (A and G) and pyrimidines (C and T) are shown at the top and bottom, respectively. Transitions occur in the cases of PuPu and PyPy, while transversion occurs only with PuPy encoding.

This additional encoding is always given in a particular GWAS data set, which leads us to consider the probabilities of PuPu, PuPy, and PyPy. These will be necessary to determine asymptotics for the TiTv distance metric. Let *γ*_0_, *γ*_1_, and *γ*_2_ denote the probabilities of PuPu, PuPy, and PyPy, respectively, for the *p* loci of data matrix *X*. In real data, there are approximately twice as many transitions as there are transversions. That is, the probability of a transition P(Ti) is approximately twice the probability of transversion P(Tv). It is likely that any particular data set will not satisfy this criterion exactly. In this general case, we have P(Ti) being equal to some multiple *η* times P(Tv). In order to enforce this general constraint in simulated data, we define the following set of equalities

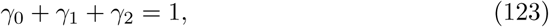

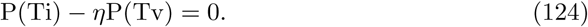

The sum-to-one constraint (Eq. 123) is natural in this context because there are only three possible genotype encodings at a particular locus, which are PuPu, PuPy, and PyPy. Solving the Ti/Tv ratio constraint (Eq. 124) for *η* gives

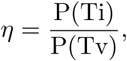

which is easily computed in a real data set by dividing the fraction of Ti out of the total *p* loci by the fraction of Tv out of the total *p* loci. We will use the simplified notation *η* = Ti/Tv to represent this factor for the remainder of this work.

Using this PuPu, PuPy, and PyPy encoding, the probability of a transversion occurring at any fixed locus *a* is given by the following

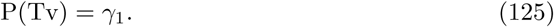

Using the sum-to-one constraint (Eqs. 123) and the probability of transversion (Eq. 124), the probability of a transition occuring at locus *a* is computed as follows

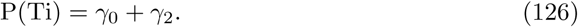

Also using the sum-to-one constraint (Eq. 123) and the Ti/Tv ratio constraint (Eq. 124), it is clear that we have 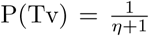 and 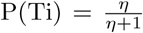. Without loss of generality, we then sample

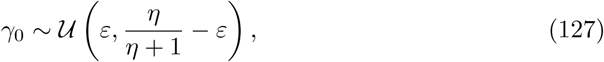

where *ε* is some small positive real number.

Then it immediately follows that we have

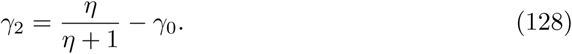

However, we can derive the mean and variance of the distance distribution induced by the TiTv metric without specifying any relationship between *γ*_0_, *γ*_1_, and *γ*_2_. We proceed by computing 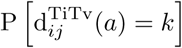 for each 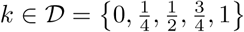. Let *y* represent a random sample of size *p* from {0, 1, 2}, where

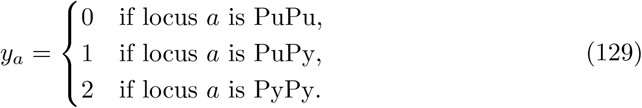

We derive 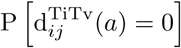 as follows

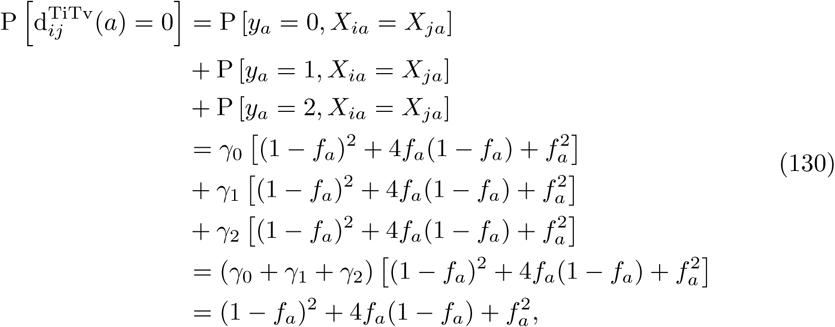

where *f*_*a*_ is the probability of a minor allele occurring at locus *a*.

We derive 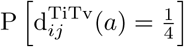 as follows

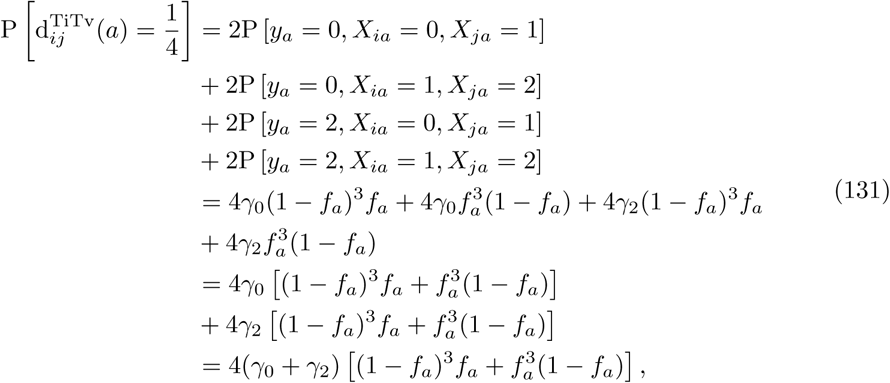

where *f*_*a*_ is the probability of a minor allele occurring at locus *a, γ*_0_ is the probability of PuPu occurring at any locus *a*, and *γ*_2_ is the probability of PyPy occurring at any locus *a*.

We derive 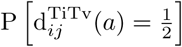 as follows

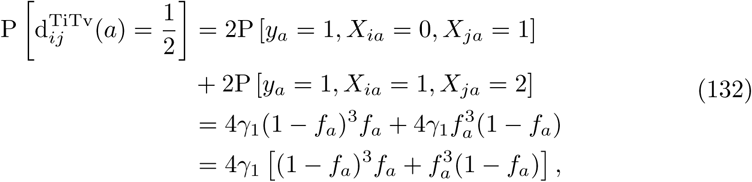

where *f*_*a*_ is the probability of a minor allele occurring at locus *a* and *γ*_1_ is the probability of PuPy occurring at any locus *a*.

We derive 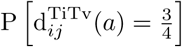 as follows

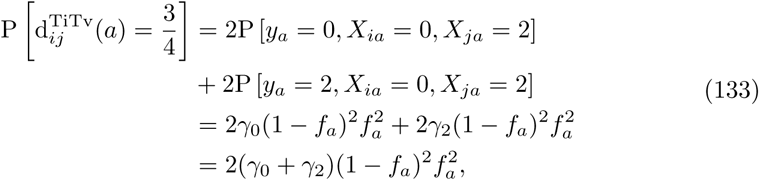

where *f*_*a*_ is the probability of a minor allele occurring at locus *a, γ*_0_ is the probability of PuPu occurring at any locus *a*, and *γ*_2_ is the probability of PyPy occurring at any locus *a*.

We derive 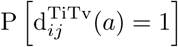 as follows

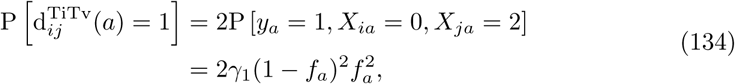

where *f*_*a*_ is the probability of a minor allele occurring at locus *a* and *γ*_1_ is the probability of PuPy occurring at any locus *a*.

Using the TiTv diff probabilities (Eqs. 130-134), we compute the expected TiTv distance between instances *i, j* ∈ ℐ as follows

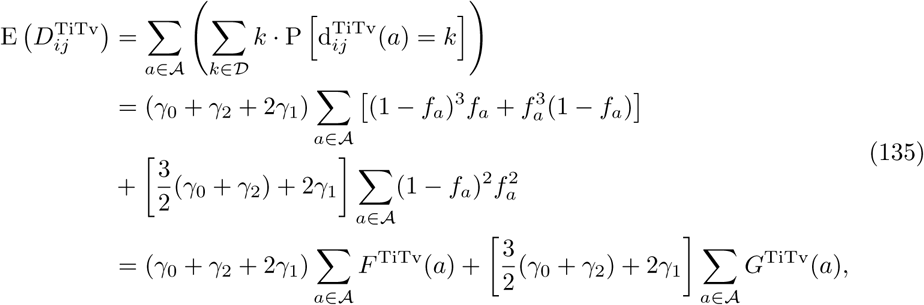

where 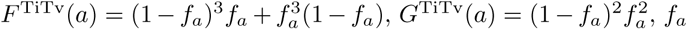 is the probability of a minor allele occurring at locus *a, γ*_0_ is the probability of PuPu occurring at any locus *a, γ*_1_ is the probability of PuPy occurring at any locus *a*, and *γ*_2_ is the probability of PyPy occurring at any locus *a*. In contrast to the expected GM and AM distances (Eqs. 114 and 119), the expected TiTv distance (Eq. 135) depends on minor allele probabilities *f*_*a*_ for all *a* ∈ 𝒜 and the genotype encoding probabilities *γ*_0_, *γ*_1_, and *γ*_2_.

The second moment about the origin for the TiTv distance is computed as follows

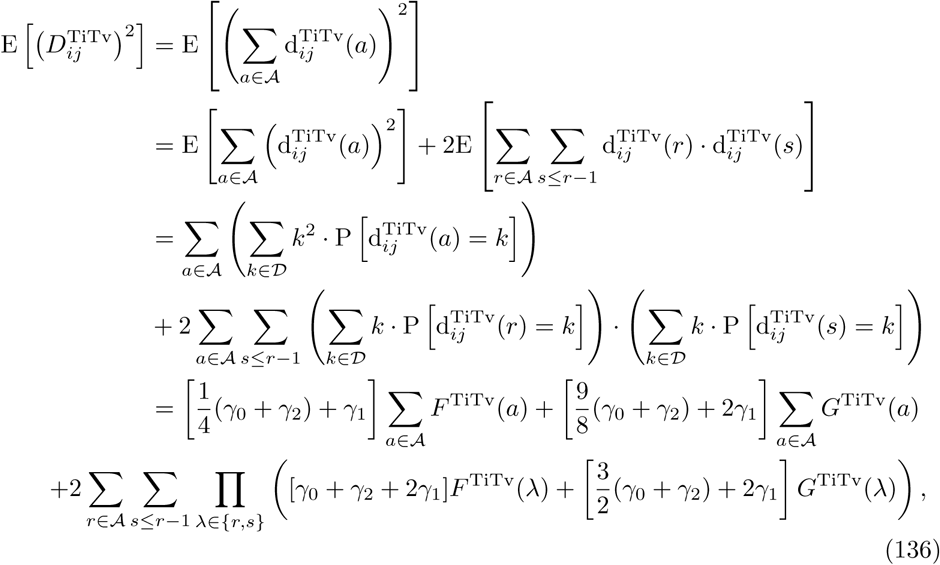

where 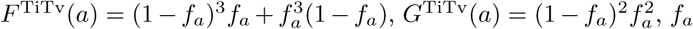 is the probability of a minor allele occurring at locus *a, γ*_0_ is the probability of PuPu occurring at any locus *a, γ*_1_ is the probability of PuPy occurring at any locus *a*, and *γ*_2_ is the probability of PyPy occurring at any locus *a*.

Using the first (Eq. 135) and second (Eq. 136) raw moments of the TiTv distance, the variance is given by

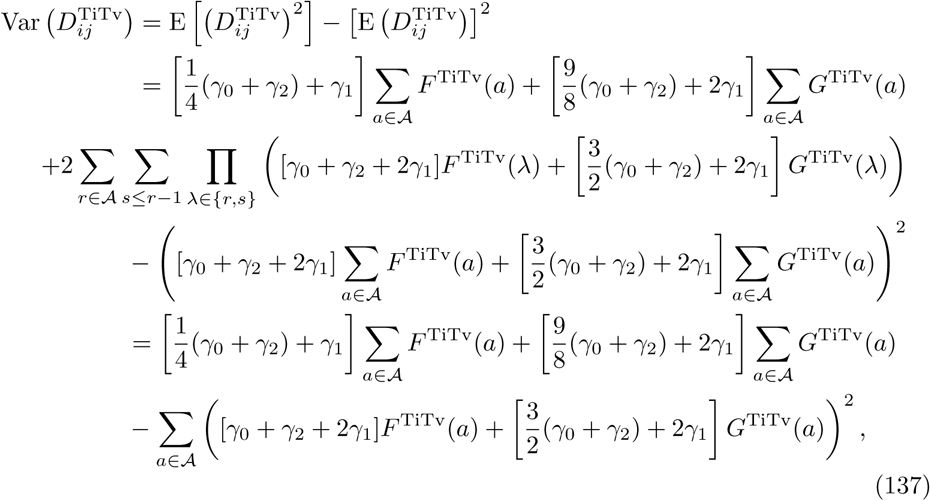

where 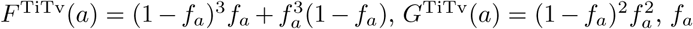 is the probability of a minor allele occurring at locus *a, γ*_0_ is the probability of PuPu occurring at any locus *a, γ*_1_ is the probability of PuPy occurring at any locus *a*, and *γ*_2_ is the probability of PyPy occurring at any locus *a*.

With the mean (Eq. 135) and variance (Eq. 137) estimates, the asymptotic TiTv distance distribution is given by the following

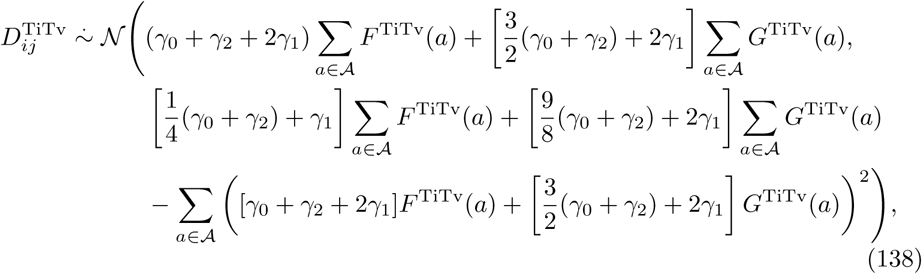

where 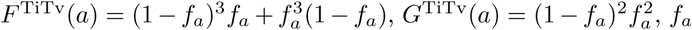 is the probability of a minor allele occurring at locus *a, γ*_0_ is the probability of PuPu occurring at any locus *a, γ*_1_ is the probability of PuPy occurring at any locus *a*, and *γ*_2_ is the probability of PyPy occurring at any locus *a*.

Given upper and lower bounds *l* and *u*, respectively, of the success probability sampling interval, the average success probability (or average MAF) is computed as follows

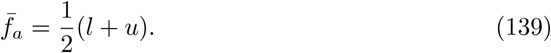

The maximum TiTv distance occurs at 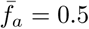 for any fixed Ti/Tv ratio *η* (Eq. 124), which is the inflection point about which the minor allele changes at locus *a* (Fig. 3). If few minor alleles are present 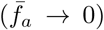, the predicted TiTv distance approaches 0. The same is true after the minor allele switches 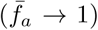. To explore how TiTv distance changes with increased minor allele frequency, we fixed the Ti/Tv ratio *η* and generated simulated TiTv distances for 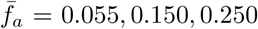, and 0.350 (Fig. 4A). For fixed *η*, TiTv distance increases significantly with increased 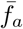. We similarly fixed the average minor allele frequency 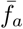 and generated simulated TiTv distances for *η* = Ti/Tv = 0.5, 1, 1.5, and 2 (Fig. 4C). The TiTv distance decreases slightly with increased *η* = Ti/Tv. As *η* → 0^+^, the data is approaching all Tv and no Ti, which means the TiTv distance is larger by definition. On the other hand, the TiTv distance decreases as *η* → 2^−^ because the data is approaching approximately twice as many Ti as there are Tv, which is typical for GWAS data in humans.

**Fig 3.**
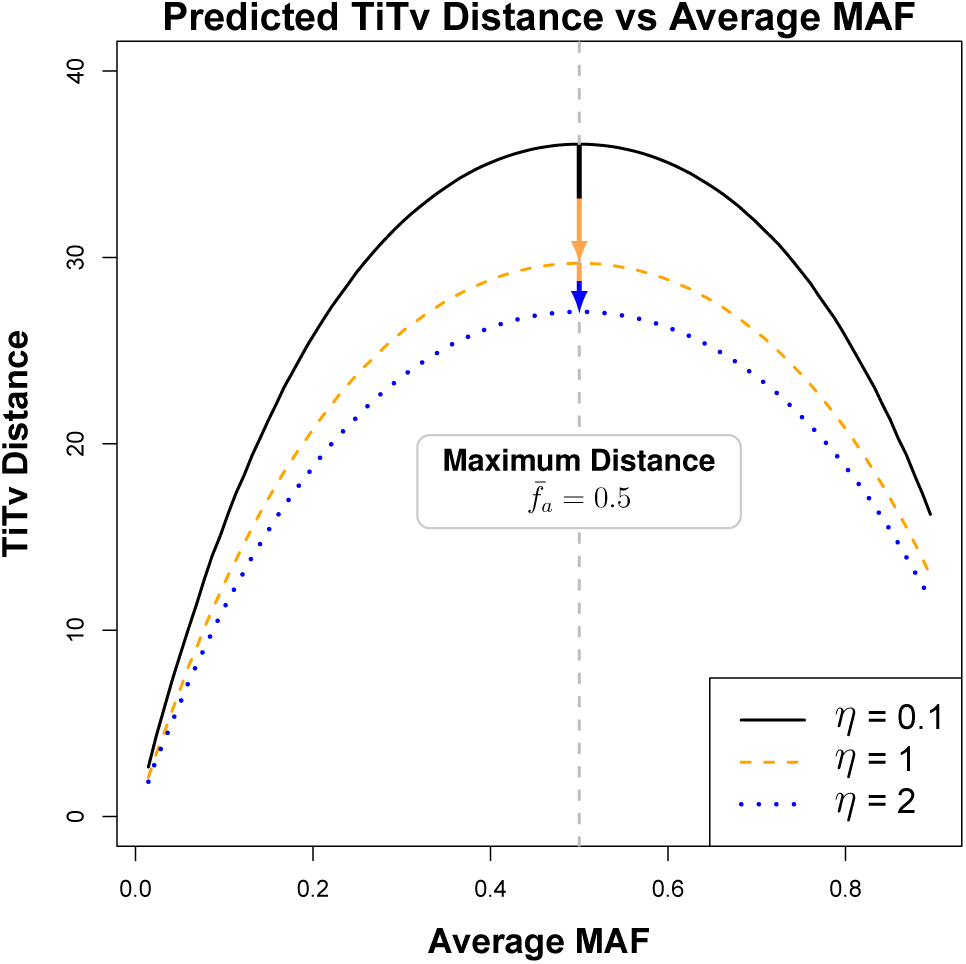
Predicted average TiTv distance as a function of average minor allele frequency 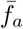 (see Eq. 139). Success probabilities *f*_*a*_ are drawn from a sliding window interval from 0.01 to 0.9 in increments of about 0.009 and *m* = *p* = 100. For *η* = 0.1, where *η* is the Ti/Tv ratio given by Eq. 123, Tv is ten times more likely than Ti and results in larger distance. Increasing to *η* = 1, Tv and Ti are equally likely and the distance is lower. In line with real data for *η* = 2, Tv is half as likely as Ti so the distances are relatively small.

**Fig 4.**
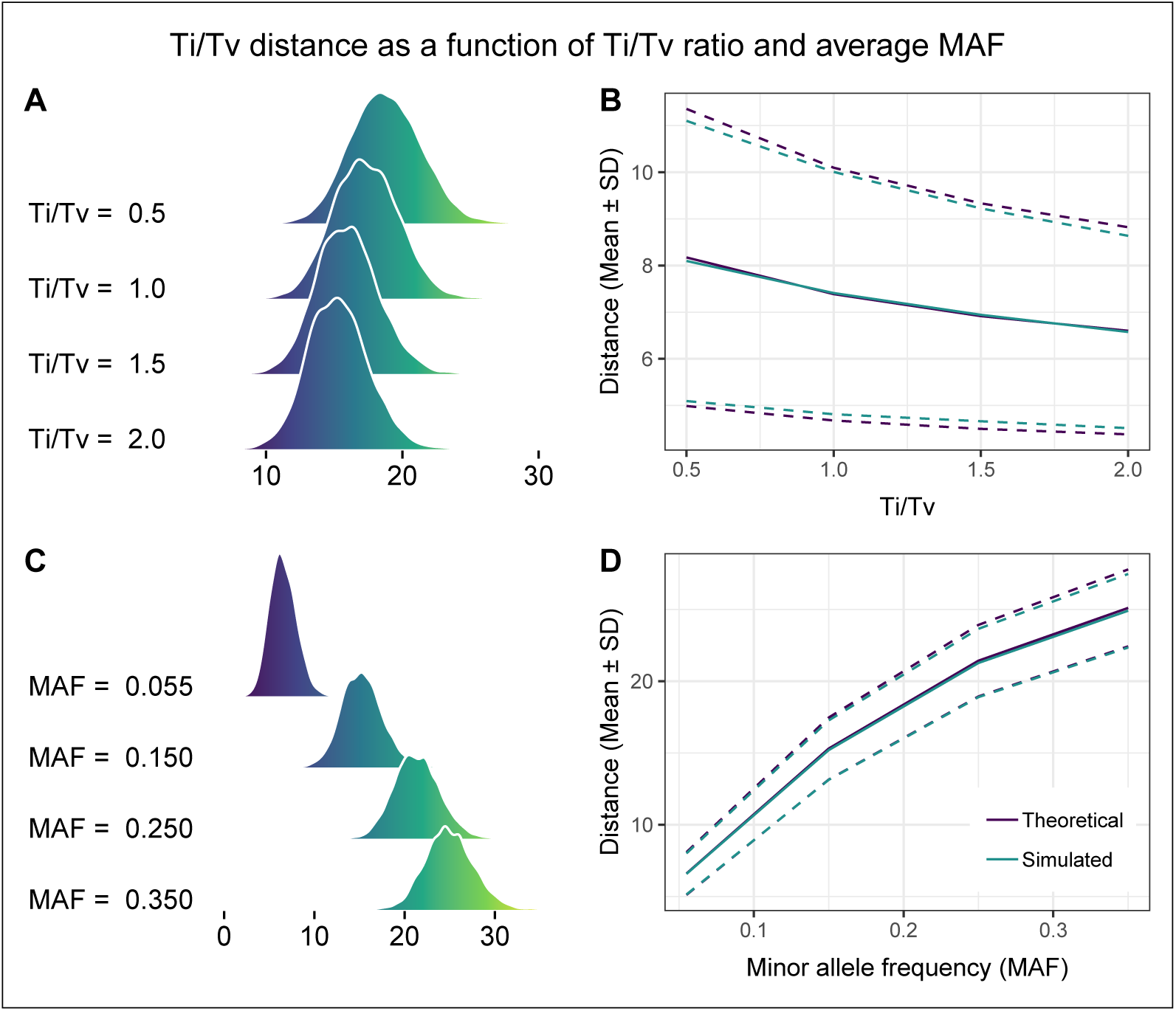
Density curves and moments of TiTv distance as a function of average MAF 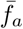, given by Eq. 139, and Ti/Tv ratio *η*, given by Eq. 124. We fix *m* = *p* = 100 for all simulated TiTv distances. (**A**) For fixed 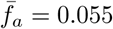, TiTv distance density is plotted as a function of increasing *η*. TiTv distance decreases as *η* increases. For *η* = Ti/Tv = 0.5, there are twice as many transversions as there are transitions. On the other hand, *η* = Ti/Tv = 2 indicates that there are half as many transversions as transitions. Since transversions encode a larger magnitude distance than transitions, this behavior is expected. (**B**) Simulated and predicted mean ± SD are shown as a function of increasing Ti/Tv ratio *η*. Distance decreases as Ti/Tv increases. Theoretical and simulated moments are approximately the same. (**C**) For fixed *η* = 2, TiTv distance density is plotted as a function of increasing 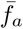. TiTv distance increases as 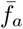 approaches maximum of 0.5, which means that there is about the same frequency of minor alleles as major alleles. (**D**) Simulated and predicted mean ± SD as a function of increasing average MAF 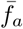. Distance increases as the number of minor alleles increases. Theoretical and simulated moments are approximately the same.

We also compared theoretical and sample moments as a function of *η* = Ti/Tv and 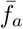 for the TiTv distance metric (Fig. 4B and D). We fixed 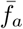 and computed the theoretical and simulated moments as a function of *η* (Fig. 4B). Theoretical average TiTv distance, given by Eq. 135, and simulated TiTv average distance are approximately equal as *η* increases. Theoretical standard deviation, given by Eq. 137, and simulated TiTv standard deviation differ slightly. We also fixed *η* and computed theoretical and sample moments as a function of 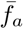 (Fig. 4D). In this case, there is approximate agreement with simulated and theoretical moments as 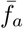 increases.

We summarize our moment estimates for GWAS distance metrics (Eqs. 110-112) (Table 4) organized by metric, statistic (mean or variance), and asymptotic formula. Next we consider the important case of distributions of GWAS distances projected onto a single attribute (Eqs. 107-109).

**Table 4.** Asymptotic estimates of means and variances of genotype mismatch (GM) (Eq. 110), allele mismatch (AM) (Eq. 111), and transition-transversion (TiTv) (Eq. 112) distance metrics in GWAS data (*p* ≫ 1). GWAS data *X*_*ia*_ ∼ ℬ(2, *f*_*a*_), where *f*_*a*_ for all *a* ∈ 𝒜 are the probabilities of a minor allele occurring at locus *a*. For the TiTv distance metric, we have the additional encoding that uses *γ*_0_ = P(PuPu), *γ*_1_ = P(PuPy), and *γ*_2_ = P(PyPy).

| GWAS-Metric | Stat | Formula (Eq. #) |
| --- | --- | --- |
| GM<br>(Eq. 110) | mean | $2 \sum_{a \in \mathcal{A}} F^{\text{GM}}(a) \quad (114)$<br>where $F^{\text{GM}}(a) = 2(1 - f_a)^3 f_a + 2f_a^3(1 - f_a) + (1 - f_a)^2 f_a^2$ |
| | variance | $2 \sum_{a \in \mathcal{A}} F^{\text{GM}}(a)[1 - 2F^{\text{GM}}(a)] \quad (116)$<br>where $F^{\text{GM}}(a) = 2(1 - f_a)^3 f_a + 2f_a^3(1 - f_a) + (1 - f_a)^2 f_a^2$ |
| AM<br>(Eq. 111) | mean | $2 \sum_{a \in \mathcal{A}} F^{\text{AM}}(a) \quad (119)$<br>where $F^{\text{AM}}(a) = (1 - f_a)^3 f_a + f_a^3(1 - f_a) + (1 - f_a)^2 f_a^2$ |
| | variance | $\sum_{a \in \mathcal{A}} [G^{\text{AM}}(a) - 4(F^{\text{AM}}(a))^2] \quad (121)$<br>where $F^{\text{AM}}(a) = (1 - f_a)^3 f_a + f_a^3(1 - f_a) + (1 - f_a)^2 f_a^2$ and<br>$G^{\text{AM}}(a) = (1 - f_a)^3 f_a + f_a^3(1 - f_a) + 2(1 - f_a)^2 f_a^2$ |
| TiTv<br>(Eq. 112) | mean | $(\gamma_0 + \gamma_2 + 2\gamma_1) \sum_{a \in \mathcal{A}} F^{\text{TiTv}}(a) + \left[\frac{3}{2}(\gamma_0 + \gamma_2) + 2\gamma_1\right] \sum_{a \in \mathcal{A}} G^{\text{TiTv}}(a) \quad (135)$<br>where $F^{\text{TiTv}}(a) = (1 - f_a)^3 f_a + f_a^3(1 - f_a)$ and $G^{\text{TiTv}}(a) = (1 - f_a)^2 f_a^2$ |
| | variance | $\left[\frac{1}{4}(\gamma_0 + \gamma_2) + \gamma_1\right] \sum_{a \in \mathcal{A}} F^{\text{TiTv}}(a) + \left[\frac{9}{8}(\gamma_0 + \gamma_2) + 2\gamma_1\right] \sum_{a \in \mathcal{A}} G^{\text{TiTv}}(a) + \sum_{a \in \mathcal{A}} \left( \left[\gamma_0 + \gamma_2 + 2\gamma_1\right] F^{\text{TiTv}}(a) + \left[\frac{3}{2}(\gamma_0 + \gamma_2) + 2\gamma_1\right] G^{\text{TiTv}}(a) \right)^2 \quad (137)$<br>where $F^{\text{TiTv}}(a) = (1 - f_a)^3 f_a + f_a^3(1 - f_a)$ and $G^{\text{TiTv}}(a) = (1 - f_a)^2 f_a^2$ |

### 3.4 Distribution of one-dimensional projection of GWAS distance onto a SNP

We previously derived the exact distribution of the one-dimensional projected distance onto an attribute in continuous data (Section 2.2.3), which is used as the predictor in NPDR to calculate relative attribute importance in the form of standardized beta coefficients. GWAS data and the metrics we have considered are discrete. Therefore, we derive the density function for each diff metric (Eqs. 107-88 109), which also serves as the probability distribution for each metric, respectively.

The support of the GM metric (Eq. 107) is simply {0, 1}, so we derive the probability, 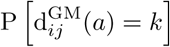, of this diff taking on each of these two possible values. First, the probability that the GM diff is equal to zero is given by

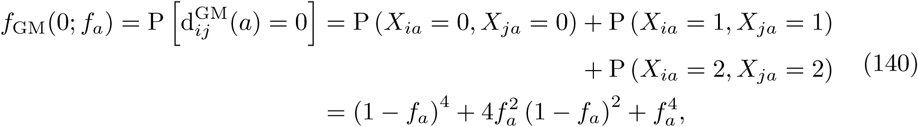

where *f*_*a*_ is the probability of a minor allele occurring at locus *a*.

Similarly, the probability that the GM diff is equal to 1 is derived as follows

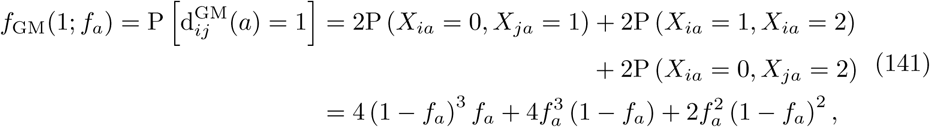

where *f*_*a*_ is the probability of a minor allele occurring at locus *a*.

This leads us to the probability distribution of the GM diff metric, which is the distribution of the one-dimensional GM distance projected onto a single SNP. This distribution is given by

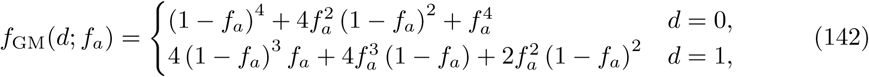

where *f*_*a*_ is the probability of a minor allele occurring at locus *a*.

The mean and variance of this GM diff distribution can easily be derived using this newly determined density function (Eq. 142). The average GM diff is given by the following

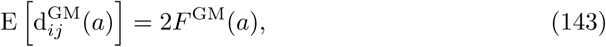

where 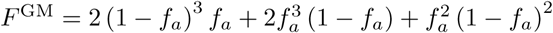 and *f*_*a*_ is the probability of a minor allele occurring at locus *a*.

The variance of the GM diff metric is given by

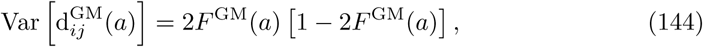

where 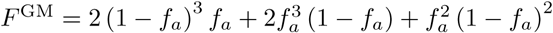 and *f*_*a*_ is the probability of a minor allele occurring at locus *a*.

The support of the AM metric (Eq. 108) is {0, 1/2, 1}. Beginning with the probability of the AM diff being equal to 0, we have the following probability

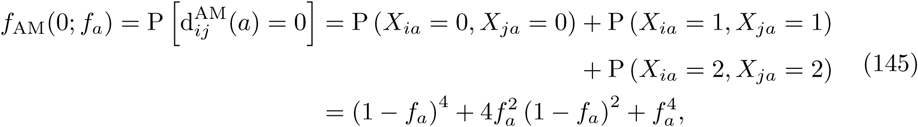

where *f*_*a*_ is the probability of a minor allele occurring at locus *a*.

The probability of the AM diff metric being equal to 1/2 is computed similarly as follows

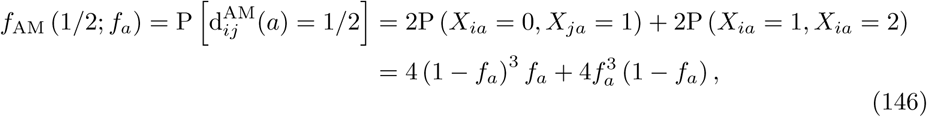

where *f*_*a*_ the probability of a minor allele occurring at locus *a*.

Finally, the probability of the AM diff metric being equal to 1 is given by the following

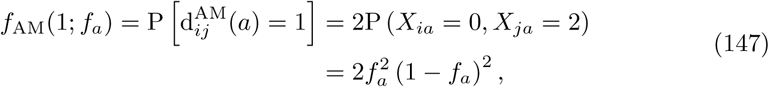

where *f*_*a*_ is the probability of a minor allele occurring at locus *a*.

As in the case of the GM diff metric, we now have the probability distribution of the AM diff metric. This also serves as the distribution of the one-dimensional AM distance projected onto a single SNP, and is given by the following

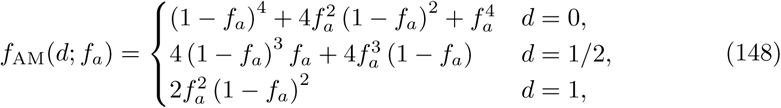

where *f*_*a*_ is the probability of a minor allele occurring at locus *a*.

The mean and variance of this AM diff distribution is derived using the corresponding density function (Eq. 148). The average AM diff is given by

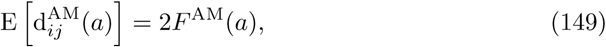

where 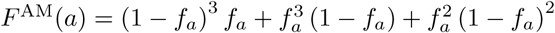 and *f*_*a*_ is the probability of a minor allele occurring at locus *a*.

The variance of the AM diff metric is given by

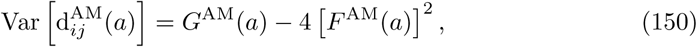

where 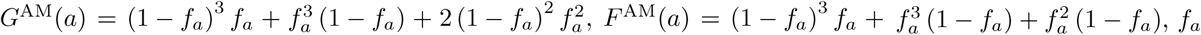 is the probability of a minor allele occurring at locus *a*.

For the TiTv diff metric (Eq. 109), the support is {0, 1/4, 1/2, 3/4, 1}. We have already derived the probability that the TiTv diff assumes each of the values of its support (Eqs. 130-134). Therefore, we have the following distribution of the TiTv diff metric

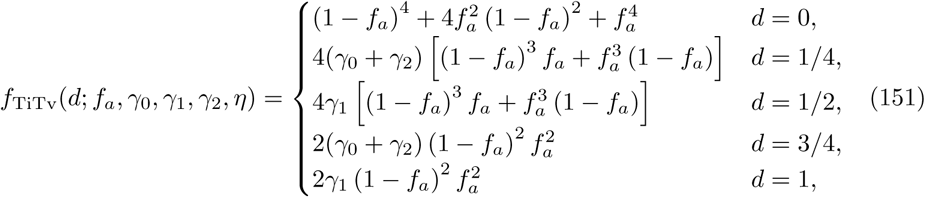

where *f*_*a*_ is the probability of a minor allele occurring at locus *a, γ*_0_ is the probability of PuPu at locus *a, γ*_1_ is the probability of PuPy at locus *a, γ*_2_ is the probability of PyPy at locus *a*, and *η* is the Ti/Tv ratio (Eq. 124).

The mean and variance of this TiTv diff distribution is derived using the corresponding density function (Eq. 151). The average TiTv diff is given by

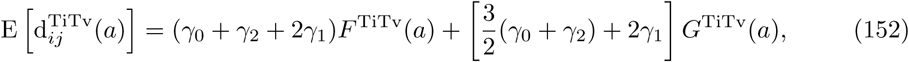

where 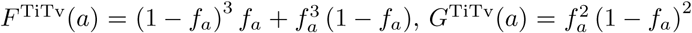, *f_a_* is the probability of a minor allele occurring at locus *a, γ*_0_ is the probability of PuPu at locus *a, γ*_1_ is the probability of PuPy at locus *a*, and *γ*_2_ is the probability of PyPy at locus *a*.

The variance of the TiTv diff metric is given by

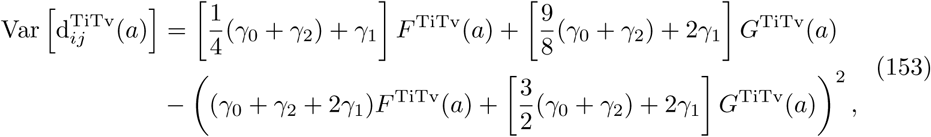

where 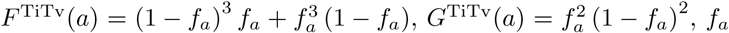 is the probability of a minor allele occurring at locus *a, γ*_0_ is the probability of PuPu at locus *a, γ*_1_ is the probability of PuPy at locus *a*, and *γ*_2_ is the probability of PyPy at locus *a*.

These novel distribution results for the projection of pairwise GWAS distances onto a single genetic variant, as well as results for the full space of *p* variants, can inform NPDR and other nearest-neighbor distance-based feature selection algorithms. Next we introduce our new diff metric and distribution results for time series derived correlation-based data, with a particular application to resting-state fMRI.

## 4 Time series correlation-based distance distribution

For time series correlation-based data, we consider the case where there are *m* correlation matrices *A*^(*p*×*p*)^ (one matrix for each subject). In particular, we have in mind the application of resting-state fMRI (rs-fMRI) data. The derivations that follow, however, are relevant to all correlation-based data with the assumptions we note. The attributes in rs-fMRI are commonly Regions of Interest (ROIs), which are collections of spatially proximal voxels [22]. Correlation in their time-series activity arise between different ROIs at the voxel level or for a given brain atlas [23]. Because the attributes of interest (*a*) are the ROIs themselves, we propose the following attribute projection (diff)

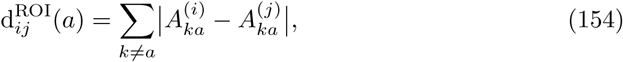

where 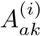 and 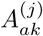 are the correlations between ROI *a* and ROI *k* for instances *i, j* ∈ ℐ, respectively. With this rs-fMRI diff, we define the pairwise distance between two instances *i, j* ∈ ℐ as follows

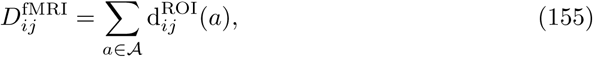

which is based on Manhattan (*q* = 1). This metric may be expanded to general *q*, but we only consider *q* = 1.

In order for comparisons between different correlations to be possible, we first perform a Fisher r-to-z transform on the correlations. This transformation makes the data approximately normally distributed with stabilized variance across different samples. After this transformation, we then load all of the transformed correlations into a *p*(*p* − 1) × *m* matrix *X* (Fig. 5). Each column of *X* represents a single instance (or subject) in rs-fMRI data. Contrary to a typical *p* × *m* data set, each row does not represent a single attribute. Rather, each attribute (or ROI) is represented by *p* − 1 consecutive rows. The first *p* − 1 rows represent ROI_1_, the next *p* − 1 rows represent ROI_2_, and so on until the last *p* − 1 rows that represent ROI_*p*_. For a given column of *X*, we exclude pairwise correlations between an ROI and itself. Therefore, the matrix does not contain 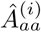 for any *i* ∈ ℐ or *a* ∈ 𝒜. Furthermore, symmetry of correlation matrices means that each column contains exactly two of each element of the upper triangle of an instance’s transformed correlation matrix. For example, 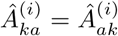 for *k* ≠ *a* and both will be contained in a given column of *X* for each *a* ∈ 𝒜. Based on our rs-fMRI diff (Eq. 154), the organization of *X* makes computation of each value of the diff very simple. In order to compute each value of the rs-fMRI diff, we just need to know the starting and ending row indices for a given ROI. Starting indices are given by

**Fig 5.**
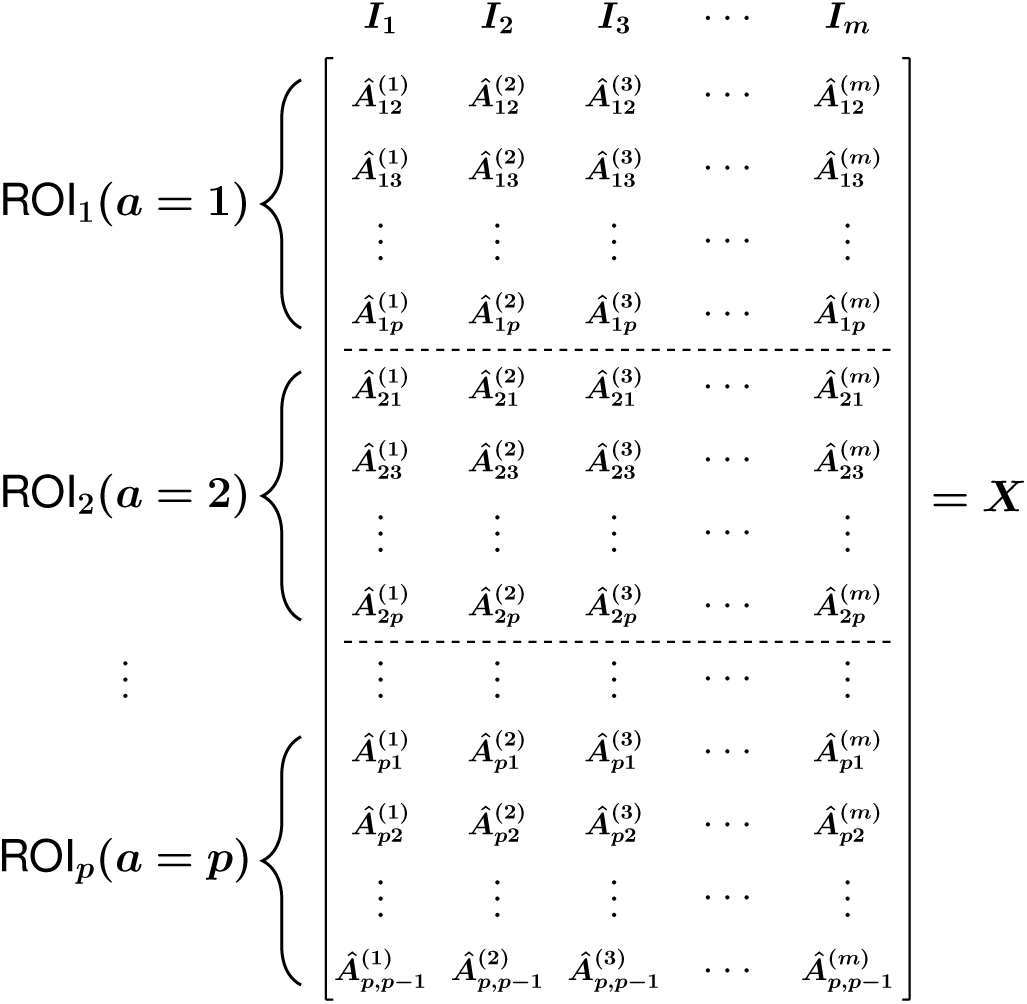
Organization based on brain regions of interest (ROIs) of resting-state fMRI correlation dataset consisting of transformed correlation matrices for *m* subjects. Each column corresponds to an instance (or subject) *I*_*j*_ and each subset of rows corresponds to the correlations for an ROI attribute (*p* sets). The notation 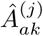 represents the r-to-z transformed correlation between attributes (ROIs) *a* and *k* ≠ *a* for instance *j*.

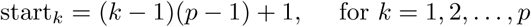

and ending indices are given by

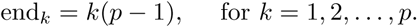

These indices allow us to extract just the rows necessary to compute the rs-fMRI diff for a fixed ROI.

We further transform the data matrix *X* by standardizing so that each of the *m* columns has zero mean and unit variance. Therefore, the data in matrix *X* are approximately standard normal. Recall that the mean (Eq. 38) and variance (Eq. 39) of the Manhattan (*L*_1_) distance distribution for standard normal data are 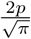 and 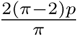, respectively. This allows us to easily derive the expected pairwise distance between instances *i, j* ∈ ℐ in rs-fMRI data as follows

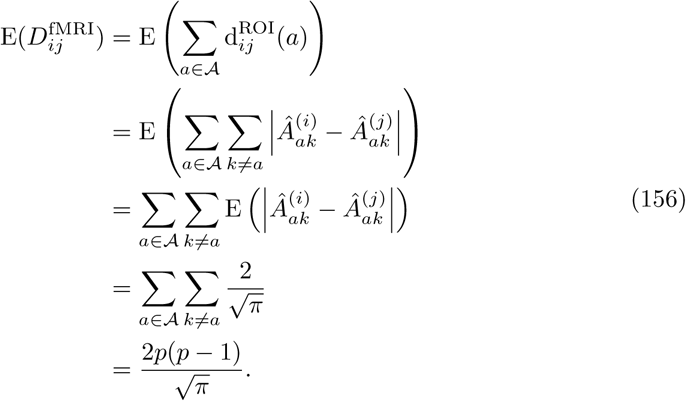

The expected pairwise rs-fMRI distance (Eq. 156) grows on the order of *p*(*p* − 1), which is the total number of transformed pairwise correlations in each column of *X* (Fig. 5). This is similar to the case of a typical *m × p* data matrix in which the data is standard normal and Manhattan distances are computed between instances.

We first derive the variance of the rs-fMRI distance by making an independence assumption with respect to the magnitude differences 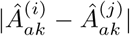 for all *k* ≠ *a* ∈ 𝒜. We observe empirically that this assumption gives a reasonable estimate of the actual variance of rs-fMRI distances in simulated data, but there is a consistent discrepancy between predicted and simulated variances. We begin our derivation of the variance of rs-fMRI distances by assuming that cross-covariances between the diffs of different pairs of ROIs are negligible. This allows us to determine the relationship between the predicted variance under the independence assumption and the simulated variance. We proceed by applying the variance operator linearly as follows

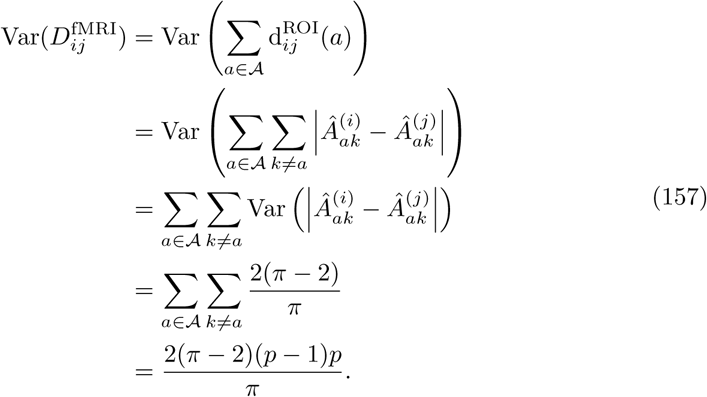

Similar to the case of an *m* × *p* data matrix containing standard normal data, we have an rs-fMRI distance variance that grows on the order of *p*(*p* −1), which is the total number of pairwise associations in a column of data matrix *X* (Fig. 5). Therefore, the expected rs-fMRI distance (Eq. 156) and the variance of the rs-fMRI distance (Eq. 157) increase on the same order.

The independence assumption used to derive the variance of our rs-fMRI distance metric (Eq. 157) is not satisfied because a single value of the diff (Eq. 154) includes the same fixed ROI, *a*, for each term in the sum for all *k* ≠ *a*. Therefore, the linear application of the variance operator we have previously employed does not account for the additional cross-covariance that exists. However, we have seen empirically that the theoretical variance of the distance we computed for the rs-fMRI distance metric (Eq. 157) still reasonably approximates the sample variance, there is a slight discrepancy between our theoretical rs-fMRI distance metric variance (Eq. 157) and the sample variance. More precisely, the formula we have given for the variance (Eq. 157) consistently underestimates the sample variance of the rs-fMRI distance. To adjust for this discrepancy, we determine a corrected formula by assuming that there is dependence between the terms of the rs-fMRI diff and estimate the cross-covariance between rs-fMRI diffs of different pairs of ROIs.

We begin the derivation of our corrected formula by writing the variance as a two-part sum, where the first term in the sum involves the variance of the magnitude difference 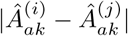 and then second term involves the cross-covariance of the rs-fMRI diff for distinct pairwise ROI-ROI associations. This formulation is implied in our previous derivation of the variance, but our independence assumption allowed us to assume that all terms in the second part of the two-part sum were zero. Our formulation of the variance is given by the following

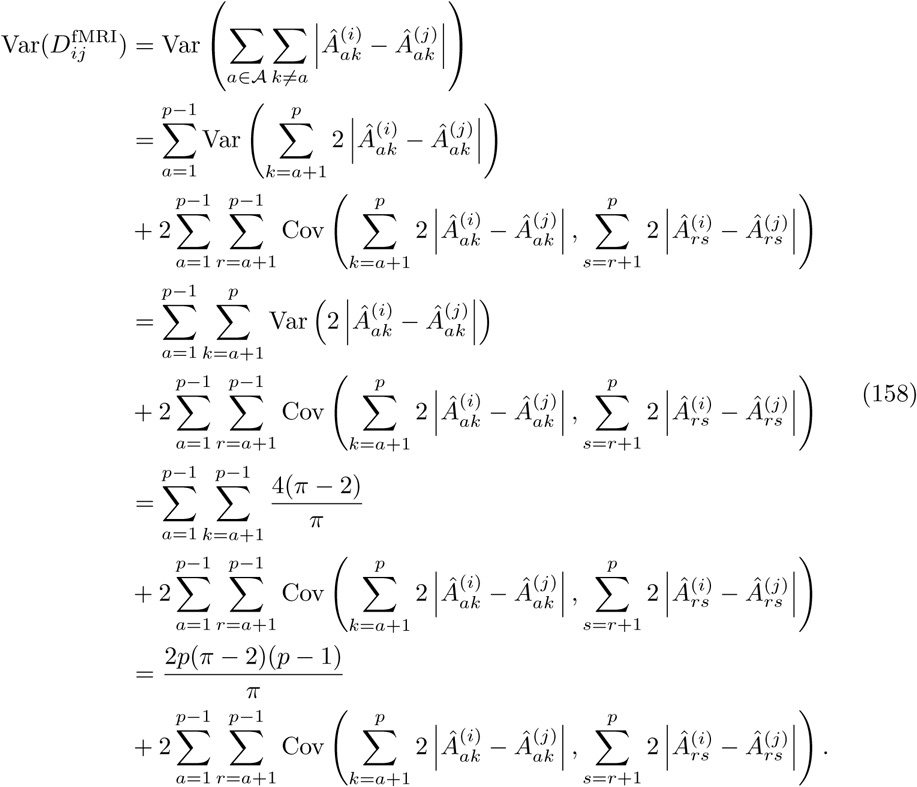

In order to have a formula in terms of the number of ROIs *p* only, we estimate the double sum on the right-hand side of the equation of rs-fMRI distance variance (Eq. 158). Through simulation, it can be seen that the difference between sample variance 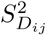 and 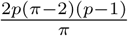 has a quadratic relationship with *p*. More explicitly, we have the following relationship

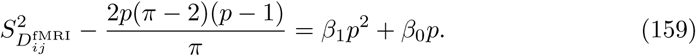

where *β*_0_ and *β*_1_ are the coefficients we must estimate in order to approximate the cross-covariance term in the right-hand side of the rs-fMRI distance variance equation (Eq. 158).

The coefficient estimates found through least squares fitting are *β*_1_ = − *β*_0_ ≈0.08. These estimates allow us to arrive at a functional form for the double sum in the right-hand side of the rs-fMRI distance variance equation (Eq. 158) that is proportional to 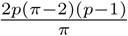. That is, we have the following formula for approximating the double sum

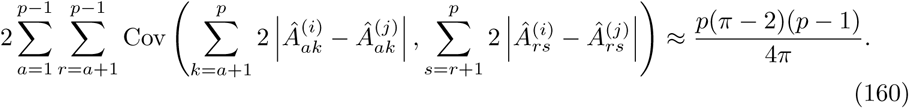

Therefore, the variance of the rs-fMRI distances is approximated well by the following

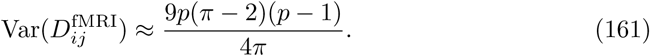

With the mean (Eq. 156) and variance (Eq. 161) estimates, we have the following asymptotic distribution for rs-fMRI distances

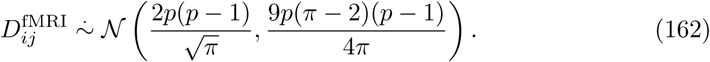

### 4.1 Max-min normalized time series correlation-based distance distribution

Previously (Section 2.4) we determined the asymptotic distribution of the sample maximum of size *m* from a standard normal distribution. We can naturally extend these results to our transformed rs-fMRI data because *X* (Fig. 5) is approximately standard normal. We proceed with the definition of the max-min normalized rs-fMRI pairwise distance.

Consider the max-min normalized rs-fMRI distance given by the following equation

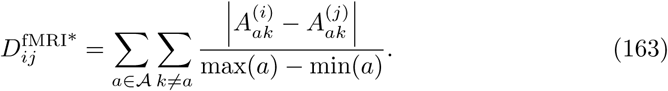

Assuming that the data *X* has been r-to-z transformed and standardized, we can easily compute the expected attribute range and variance of the attribute range. The expected maximum of a given attribute in data matrix *X* is estimated by the following

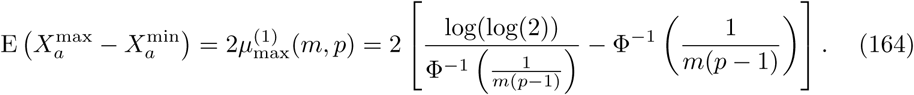

The variance can be esimated with the following

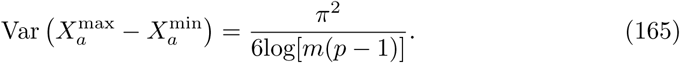

Let 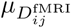 and 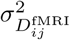 denote the mean and variance of the rs-fMRI distance distribution given by Eqs. 156 and 161. Using the formulas for the mean and variance of the max-min normalized distance distribution given in Eq. 89, we have the following asymptotic distribution for the max-min normalized rs-fMRI distances

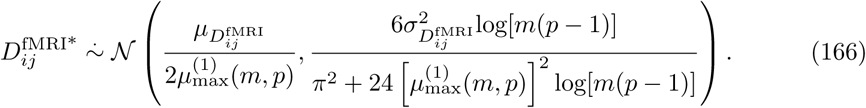

### 4.2 One-dimensional projection of rs-fMRI distance onto a single ROI

Just as in previous sections (Sections. 2.2.3 and 3.4), we now derive the distribution of our rs-fMRI diff metric (Eq. 154). Unlike what we have seen in previous sections, we do not derive the exact distribution for this diff metric. We have determined empirically that the rs-fMRI diff is approximately normal. Although the rs-fMRI diff is a sum of *p* − 1 magnitude differences, the Classical Central Limit Theorem does not apply because of the dependencies that exist between the terms of the sum. Examination of histograms and quantile-quantile plots of simulated values of the rs-fMRI diff easily indicate that the normality assumption is safe. Therefore, we derive the mean and variance of the approximately normal distribution of the rs-fMRI diff. As we have seen previously, this normality assumption is reasonable even for small values of *p*.

The mean of the rs-fMRI diff is derived by fixing a single ROI *a* and considering all pairwise associations with other ROIs *k* ≠ 𝒜. This is done as follows

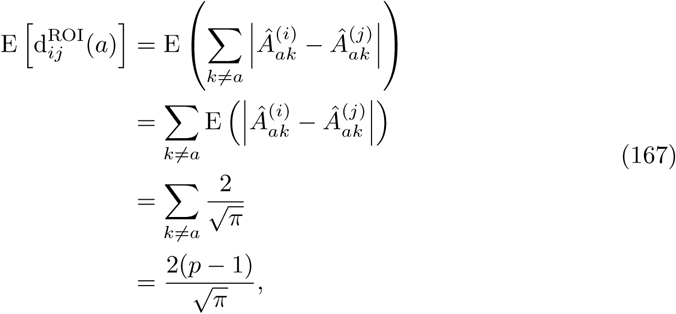

where *a* is a single fixed ROI.

Considering the variance of the rs-fMRI diff metric, we have two estimates. The first estimate uses the variance operator in a linear fashion, while the second will simply be a direct implication of the corrected formula of the variance of rs-fMRI pairwise distances (Eq. 161). Our first estimate is derived as follows

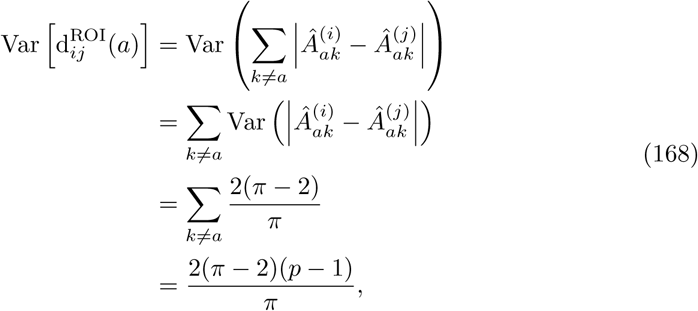

where *a* is a single fixed ROI.

Using the corrected rs-fMRI distance variance formula (Eq. 161), our second estimate of the rs-fMRI diff variance is given directly by the following

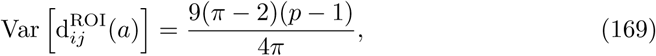

where *a* is a single fixed ROI.

Empirically, the first estimate (Eq. 168) of the variance of our rs-fMRI diff is closer to the sample variance than the second estimate (Eq. 169). This is due to fact that we are considering only a fixed ROI *a* ∈ 𝒜, so the cross-covariance between the magnitude differences 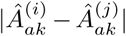 for different pairs of ROIs (*a* and *k* ≠ *a*) is negligible here. When considering all ROIs *a* ∈ 𝒜, these cross-covariances are no longer negligible. Using the first variance estimate (Eq. 168) and the estimate of the mean (Eq. 167), we have the following asymptotic distribution of the rs-fMRI diff

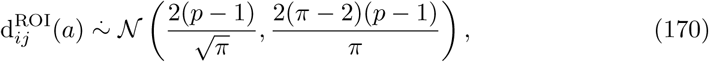

where *a* is a single fixed ROI.

### 4.3 Normalized Manhattan (*q* = 1) for rs-fMRI

Substituting the non-normalized mean (Eq. 156) into the equation for the mean of the max-min normalized rs-fMRI metric (Eq. 166), we have the following

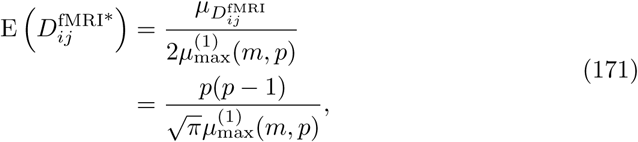

where 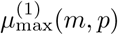 (Eq. 164) is the expected maximum of a single ROI in a data set with *m* instances and *p* ROIs.

Similarly, the variance of 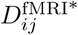 is given by

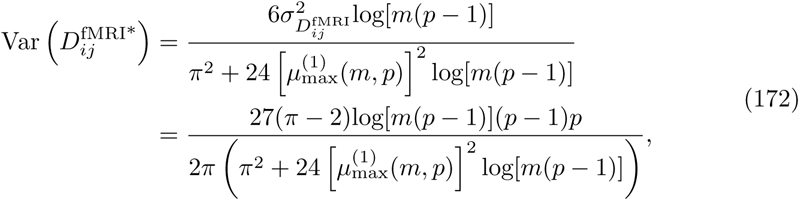

where 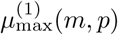 (Eq. 164) is the expected maximum of a single ROI in a data set with *m* instances and *p* ROIs.

We summarize the moment estimates for the rs-fMRI metrics for correlation-based data derived from time series (Table 5). We organize this summary by standard and attribute range-normalized rs-fMRI distance metric, statistic (mean or variance), and asymptotic formula.

**Table 5.** Aymptotic means and variances for the new standard (Eq. 155) and max-min normalized (Eq. 163) rs-fMRI distance metrics.

| rs-fMRI - Metric | Stat | Formula (Eq. #) |
| --- | --- | --- |
| standard<br>(Eq. 155) | mean | $\frac{2p(p-1)}{\sqrt{\pi}} \quad (156)$ |
| | variance | $\frac{9p(\pi-2)(p-1)}{4\pi} \quad (158)$ |
| max-min<br>normalized<br>(Eq. 163) | mean | $\frac{\mu_{D_{ij}}}{2\mu_{\max}^{(1)}(m, p)} \quad (171)$<br>where $\mu_{D_{ij}}$ and $\mu_{\max}^{(1)}(m, p)$ are given by Eqs. 156 and 164 |
| | variance | $\frac{6\sigma_{D_{ij}}^2 \log[m(p-1)]}{\pi^2 + 24 \left[ \mu_{\max}^{(1)}(m, p) \right]^2 \log[m(p-1)]} \quad (172)$<br>where $\sigma_{D_{ij}}^2$ and $\mu_{\max}^{(1)}(m, p)$ are given by Eqs. 156 and 164 |

## 5 Comparison of theoretical and sample moments

We compare our analytical asymptotic estimates of sample moments for distributions of pairwise distances in high attribute dimension by generating random data for various dimensions *m* and *p* (Fig. 6). We fix *m* = 100 samples and compute Manhattan (Eq. 1) distance matrices from standard normal data for *p* = 1000, 2000, 3000, 4000, and 5000 attributes. For each value of *p*, we generate 20 random datasets and compute the mean and standard deviation of pairwise distances. We then average these 20 simulated means and standard deviations. For comparison, we compute the theoretical moments (Eqs. 38 and 39) for each value of *p* and fixed *m* = 100 from the theoretical formulas. Scatter plots of theoretical versus simulated mean (Fig. 6A) and theoretical versus simulated standard deviation (Fig. 6B) indicate that our theoretical asymptotic formulas for sample moments are reliable for both large and relatively small numbers of attributes. For other combinations of data type, distance metric, sample size *m*, and number of attributes *p*, we find similar agreement between theoretical formulas and simulated moments (not shown).

**Fig 6.**
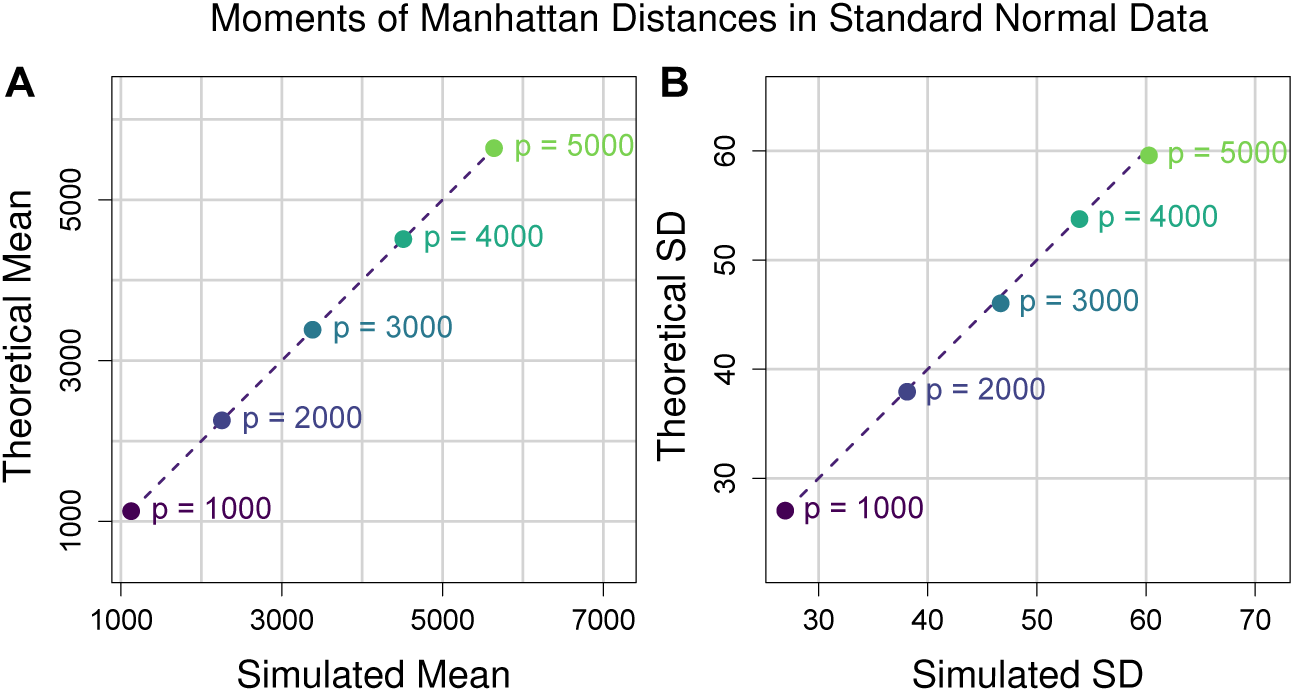
Comparison of theoretical and sample moments of Manhattan (Eq. 1) distances in standard normal data. (**A**) Scatter plot of theoretical vs simulated mean Manhattan distance (Eq. 38). Each point represents a different number of attributes *p*. For each value of *p* we fixed *m* = 100 and generated 20 distance matrices from standard normal data and computed the average simulated pairwise distance from the 20 iterations. The corresponding theoretical mean was then computed for each value of *p* for comparison. The dashed line represents the identity (or *y* = *x*) line for reference. (**B**) Scatter plot of theoretical vs simulated standard deviation of Manhattan (Eq. 1) distance (Eq. 39). These standard deviations come from the same random distance matrices for which mean distance was computed for **A**. Both theoretical mean and standard deviation approximate the simulated moments quite well.

## 6 Effects of correlation on distances

All of the derivations presented in previous sections were for the cases where there is no correlation between instances or between attributes. We assumed that any pair (*X*_*ia*_, *X*_*ja*_) of data points for instances *i* and *j* and fixed attribute *a* were independent and identically distributed. This was assumed in order to determine asymptotic estimates in null data. That is, data with no main effects, interaction effects, or pairwise correlations between attributes. Within this simplified context, our asymptotic formulas for distributional moments are reliable. However, in real data are numerous statistical effects that impact distance distributional properties. We find that deviation from normality is caused primarily by large magnitude pairwise correlation between attributes. Pairwise attribute correlation can be the result of main effects, where attributes have different within-group means. On the other hand, there could be an underlying interaction network in which there are strong associations between attributes. If attributes are differentially correlated between phenotype groups, then interactions exist that change the distance distribution. In the following few sections, we consider particular cases of the *L*_*q*_ metric for continuous and discrete data under the effects of pairwise attribute correlation.

### 6.1 Continuous data

Without loss of generality, suppose we have *X*^(*m*×*p*)^ where *X*_*ia*_ ∼ 𝒩 (0, 1) for all *i* = 1, 2, …, *m* and *a* = 1, 2, …, *p*, and let *m* = *p* = 100. We consider only the *L*_2_ (Euclidean) metric (Eq. 1, *q* = 2). We explore the effects of correlation on these distances by generating simulated data sets with increasing strength of pairwise attribute correlation and then plotting the density curve of the induced distances (Fig. 7A). Deviation from normality in the distance distribution is directly related to the average absolute pairwise correlation that exists in the simulated data. This measure is given by

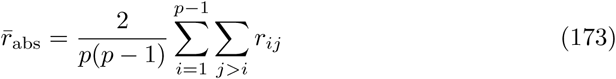

where *r*_*ak*_ is the correlation between attributes *a, k* ∈ 𝒜 across all instances *m*. Distances generated on data without correlation closely approximate a Gaussian. The mean (Eq. 50) and variance (Eq. 49) of the uncorrelated distance distribution are given by substituting *p* = 100 for the mean. As 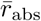 increases, positive skewness and increased variability in distances emerges. The predicted and sample means, however, are approximately the same between correlated and uncorrelated distances due to linearity of the expectation operator. Because of the dependencies between attributes, the predicted variance of 1 for *L*_2_ on standard normal data obviously no longer holds.

**Fig 7.**
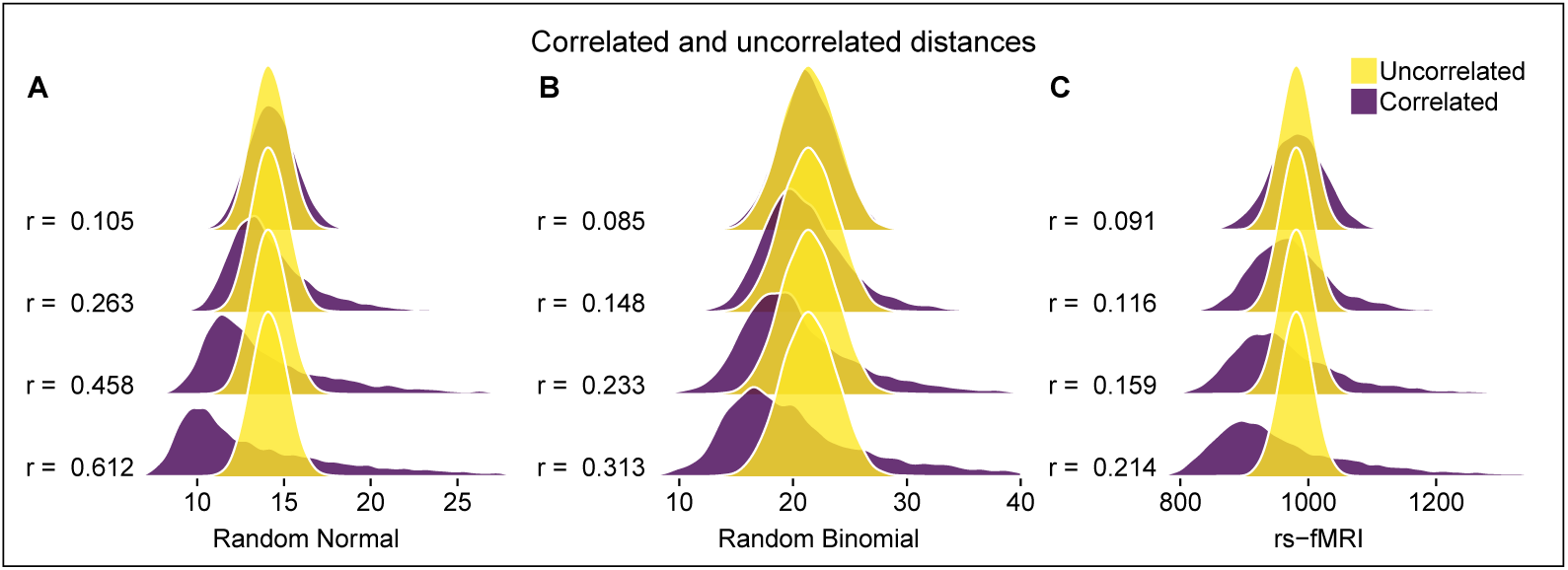
Distance densities from uncorrelated vs correlated bioinformatics data. (**A**) Euclidean distance densities for random normal data with and without correlation. Correlated data was created by multiplying random normal data by upper-triangular Cholesky factor from randomly generated correlation matrix. We created correlated data for average absolute pairwise correlation (Eq. 173) 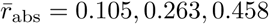, and 0.612. (**B**) TiTv distance densities for random binomial data with and without correlation. Correlated data was created by first generating correlated standard normal data using the Cholesky method from (A). Then we applied the standard normal CDF to create correlated uniformly distributed data, which was then transformed by the inverse binomial CDF with *n* = 2 trials and success probabilites *f*_*a*_ for all *a* ∈ 𝒜. (**C**) Time series correlation-based distance densities for random rs-fMRI data (Fig. 5) with and without additional pairwise feature correlation. Correlation was added to the transformed rs-fMRI data matrix (Fig. 5) using the Cholesky algorithm from (A).

In order to introduce a controlled level of correlation between attributes, we created correlation matrices based on a random graph with specified connection probability, where attributes correspond to the vertices in each graph. We assigned high correlations to connected attributes from the random graph and low correlations to all non-connections. Using the upper-triangular Cholesky factor *U* for uncorrelated data matrix *X*, we computed the following product to create correlated data matrix *X*^corr^

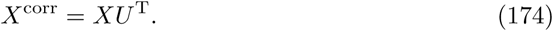

The new data matrix *X*^corr^ has approximately the same correlation structure as the randomly generated correlation matrix created from a random graph.

### 6.2 GWAS data

Analogous to the previous section, we explore the effects of pairwise attribute correlation in the context of GWAS data. Without loss of generality, we let *m* = *p* = 100 and consider only the TiTv metric (Eq. 112). To create correlated GWAS data, we first generated standard normal data with random correlation structure, just as in the previous section. We then applied the standard normal cumulative distribution function (CDF) to this correlated data in order transform the correlated standard normal variates into uniform data with preserved correlation structure. We then subsequently applied the inverse binomial CDF to the correlated uniform data with random success probabilities *f*_*a*_ for all *a* ∈ 𝒜. Each attribute *a* ∈ 𝒜 corresponds to an individual SNP in the data matrix. The resulting GWAS data set is binomial with *n* = 2 trials and has roughly the same correlation matrix as the original correlated standard normal data with which we started. Average absolute pairwise correlation 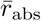 induces positive skewness in GWAS data at lower levels than in correlated standard normal data (Fig. 7B). This could have important implications in nearest neighborhoods in NPDR and similar methods.

### 6.3 Time-series derived correlation-based datasets

For our correlation data-based metric (Eq. 155), we consider additional effects of correlation between features. Without loss of generality, we let *m* = 100 and *p* = 30. We show an illustration of the effects of correlated features in this context (Fig. 7C). Based on the density estimates, it appears that correlation between features introduces positive skewness at low values of 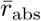. We introduced correlation to the transformed data matrix (Fig. 5) with the cholesky method used previously.

## 7 Discussion

Nearest-neighbor distance-based feature selection is a class of methods that are relatively simple to implement, and they perform well at detecting interaction effects in high dimensional data. Theoretical analysis of the limiting behavior of distance distributions for various data types and dimensions may lead to improved hyperparameter estimates of these feature selection methods. Furthermore, these theoretical results may help guide the choice of distance metric for a given dataset. Most often, distance-based feature selection methods use the *L*_*q*_ metric (Eq. 1) with *q* = 1 or *q* = 2. However, these two realizations of the *L*_*q*_ metric have considerably different expressions for the mean and variance of their respective limiting distributions. For instance, the expected distance for *L*_1_ and *L*_2_ for standard normal data is on the order of *p* (Eq. 38 and Table 2) and 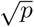 (Eq. 48 and Table 2), respectively. In addition, *L*_1_ and *L*_2_ on standard normal data have asymptotic variances on the order of *p* and 1, respectively (Eqs. 39 and 49).

These results can inform the choice of *L*_1_ or *L*_2_ depending context. For instance, distances become harder to distinguish from one another in high dimensions, which is one of the curses of dimensionality. In the case of *L*_2_, the asymptotic distribution 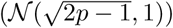 indicates that the limiting *L*_2_ distribution can be thought of simply as a positive translation of the standard normal distribution (𝒩 (0, 1)). The *L*_2_ distribution 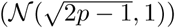 also indicates that most neighbors are contained in a thin shell far from the instance in high dimension (*p* ≫ 1). On the other hand, the *L*_1_ distances become more dispersed due to the fact that the variance of the limiting distribution is proportional to the attribute dimension *p*. This *L*_1_ dispersion could be more desirable when determining nearest neighbors because instances may be easier to distinguish with this metric. If using *L*_1_, then it may be best to use a fixed-k algorithm instead of fixed-radius. This is because fixed-radius neighborhood order could vary quite a bit considering the *L*_1_ variance is proportional to attribute dimension *p*, which in turn could affect the quality of selected attributes. If *L*_2_ is being used, then perhaps either fixed-k or fixed-radius may perform equally well because most distances will be within 1 standard deviation away from the mean.

In our analysis, we derived distance asymptotics for some of the most commonly used metrics in nearest-neighbor distance-based feature selection, as well as two new metrics for GWAS (Eq. 112) and time series correlation-based data (Eqs. 155 and 163) like resting-state fMRI. We also extended the asymptotic results for the mean and variance of the attribute range-normalized *L*_*q*_ (max-min) distance for standard normal (Eq. 89) and standard uniform (Eq. 97) data using extreme value theory. Our derivations provide an important reference for those using nearest-neighbor feature selection or classification methods in common bioinformatics data.

In this work, we expanded nearest-neighbor distance-based feature selection into the context of time series correlation-based data. Our motivation for this is partly based on the application to resting-state fMRI data. In order for this to be possible, we created a new metric (Eq. 154) that allows us to use regions of interest (ROIs) as attributes. Not all ROIs will be relevant to a particular phenotype in case-control studies, and nearest-neighbor feature selection would be a useful to tool to determine important ROIs due to interactions and to help elucidate the network structure of the brain as it relates to the phenotype of interest.

The recently introduced transition-transversion metric (Eq. 109) provides an additional dimension to the commonly used discrete metrics in GWAS nearest-neighbor distance-based feature selection. In this work, we have provided the asymptotic mean and variance of the limiting TiTv distance distribution. This novel result, as well as asymptotic estimates for the GM (Eq. 107) and AM (Eq. 108) metrics, provides an important reference to aid in neighborhood parameter selection in this context. We have also shown how the Ti/TV ratio *η* (Eq. 124) and minor allele frequency (or success probability) *f*_*a*_ affects these discrete distances. For the GM and AM metrics, the distance is solely determined by the minor allele frequencies because the genotype encoding is not taken into account. We showed how both minor allele frequency and Ti/Tv ratio uniquely affects the TiTv distance (Figs. 4A and 4C). Because transversions are more drastic forms of mutation than transitions, this additional dimension of information is important to consider, which is why we have provided asymptotic results for this metric. In addition to asymptotic *L*_*q*_ distance distributions, we have also provided the exact distributions for the one-dimensional projection of the *L*_*q*_ distance onto individual attributes (Sections. 2.2.3, 3.4, and 4.2). These distributions are important for all nearest-neighbor distance-based feature selection algorithms, such as Relief or NPDR, because the *L*_*q*_ distance is a function of the one-dimensional attribute projection (diff). In particular, these projected distance distributions are important for improving inference for predictors in NPDR, which are one-dimensional attribute projections.

Correlations between attributes and instances can cause significant deviations from the asymptotic results for uncorrelated data we have derived in this work. To illustrate this behavior, we showed how strong correlations lead to positive skewness in the distance distribution of random normal, binomial, and rs-fMRI data (Figs. 7A, 7B, and 7C). Pairwise correlation between attributes does not change the average distance, so our asymptotic results for uncorrelated data also apply when attributes are not independent. In contrast, the sample variance of distances deviates from the uncorrelated case substantially as the average absolute pairwise attribute correlation increases (Eq. 173). For fixed-radius neighborhood methods, this deviation increases the probability of including neighbors for a given instance and may reduce the power to detect interactions. The increased variability for distances with correlated data may inform the choice of metric and optimization of neighborhoods in nearest-neighbor feature selection.

