## Supplementary Figures for "Theoretical properties of nearest-neighbor distance distributions and novel metrics for high dimensional bioinformatics data"

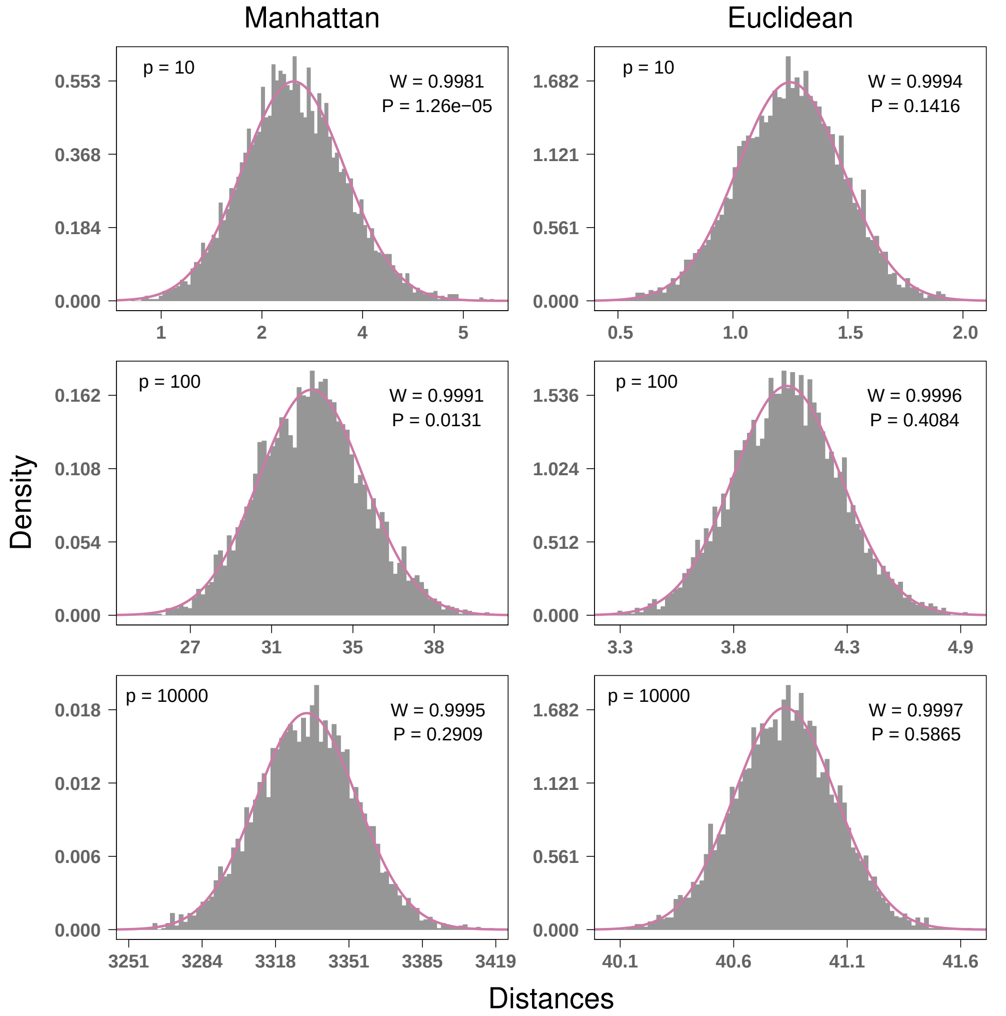


**Figure S1.** Convergence to Gaussian for Manhattan and Euclidean distances for simulated standard uniform data with $m = 100$ instances and $p = 10, 100$, and 10000 attributes. Convergence to Gaussian occurs rapidly with increasing $p$, and Gaussian is a good approximation for $p$ as low as 10 attributes. The number of attributes in bioinformatics data is typically much larger, at least on the order of $10^3$. The Euclidean metric has stronger convergence to normal than Manhattan. P values from Shapiro-Wilk test, where the null hypothesis is a Gaussian distribution.

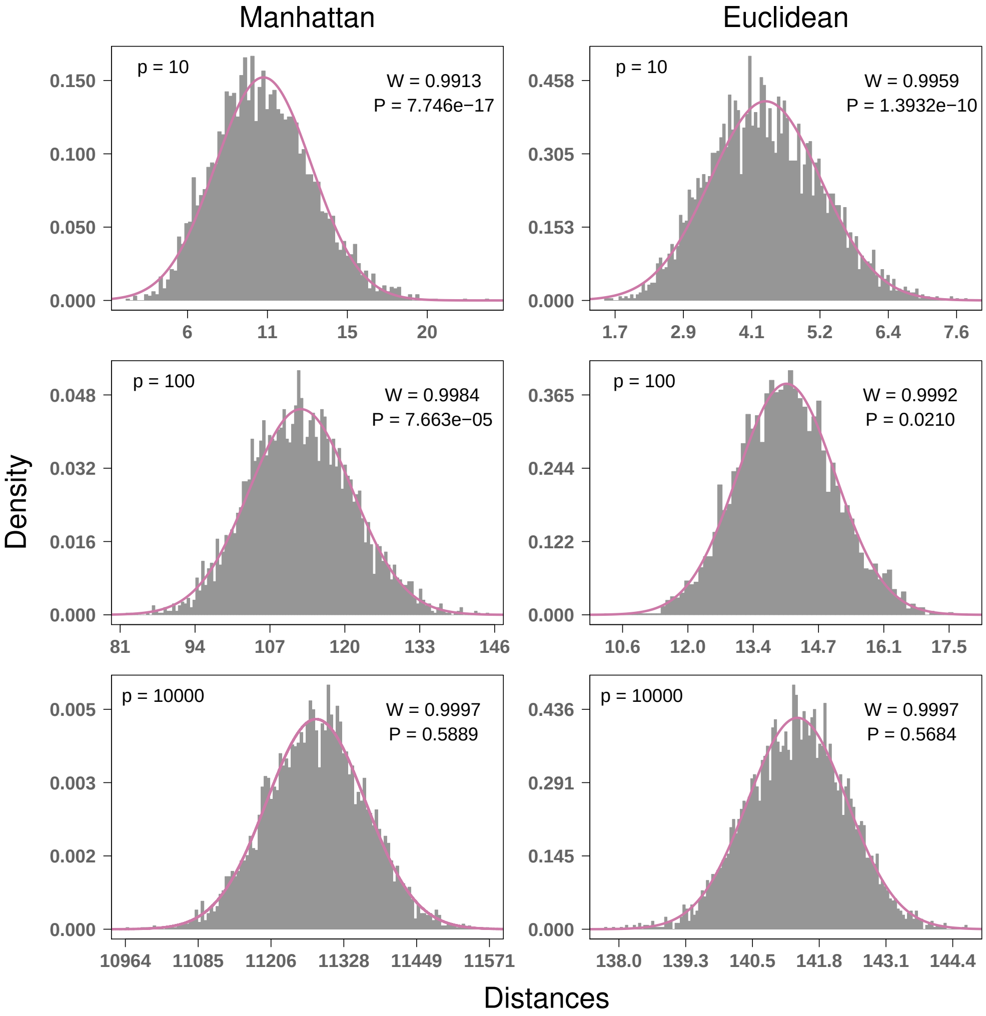


**Figure S2.** Convergence to Gaussian for Manhattan and Euclidean distances for simulated standard normal data with $m = 100$ instances and $p = 10, 100$, and 10000 attributes. Convergence to Gaussian occurs rapidly with increasing $p$, and Gaussian is a good approximation for $p$ as low as 10 attributes. The number of attributes in bioinformatics data is typically much larger, at least on the order of $10^3$. The Euclidean metric has stronger convergence to normal than Manhattan. P values from Shapiro-Wilk test, where the null hypothesis is a Gaussian distribution.

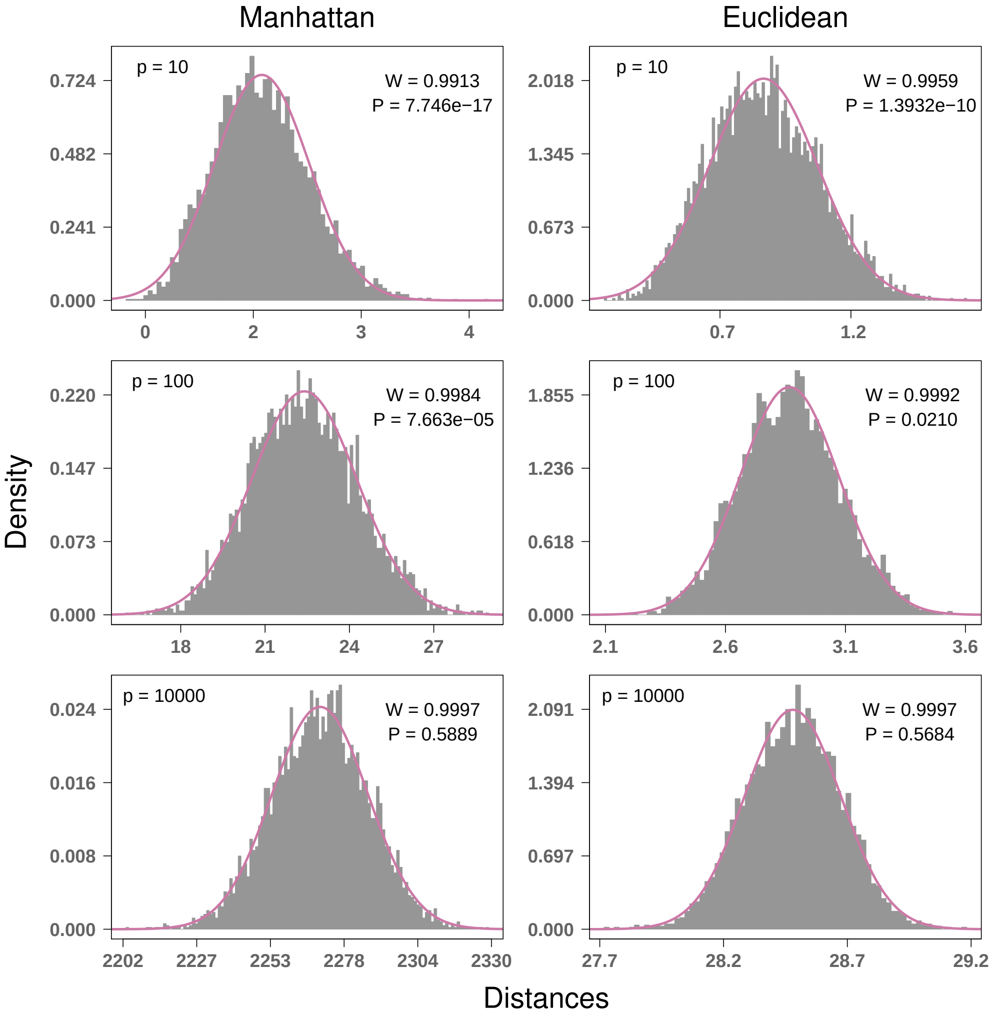


**Figure S3.** Convergence to Gaussian for max-min normalized Manhattan and Euclidean distances for simulated standard normal data with $m = 100$ instances and $p = 10, 100$, and 10000 attributes. Convergence to Gaussian occurs rapidly with increasing $p$, and Gaussian is a good approximation for $p$ as low as 10 attributes. The number of attributes in bioinformatics data is typically much larger, at least on the order of $10^3$. The Euclidean metric has stronger convergence to normal than Manhattan. P values from Shapiro-Wilk test, where the null hypothesis is a Gaussian distribution.

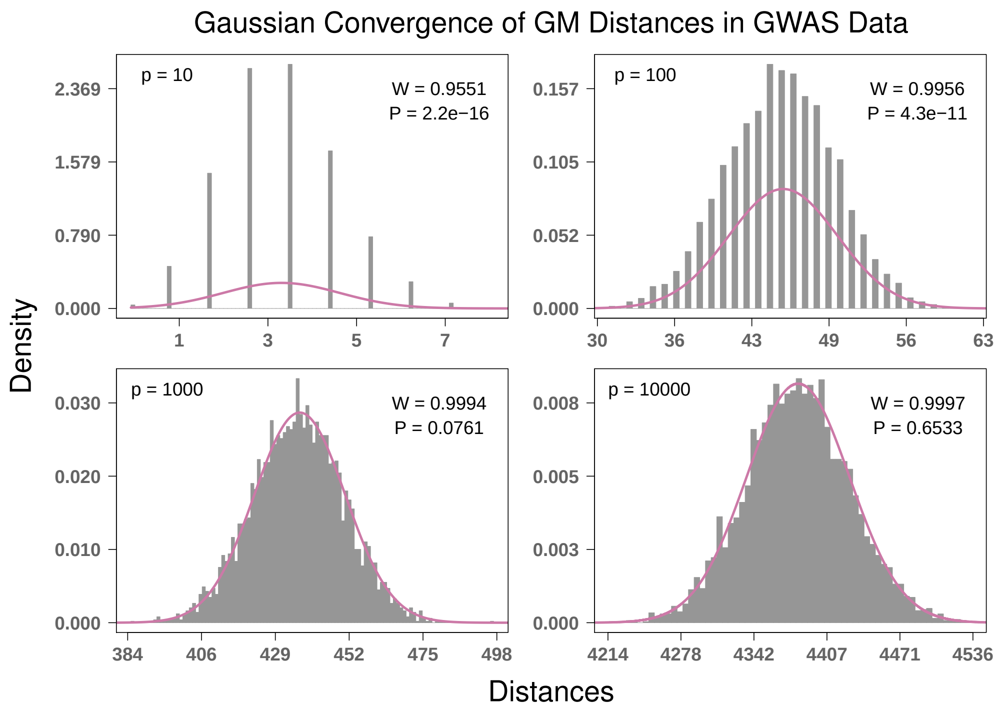


**Figure S4.** Convergence to Gaussian for GM distances for simulated binomial GWAS data with $m = 100$ instances and $p = 10, 100, 1000$, and 10000 attributes. The average MAF was set to 0.205 for all simulations. Convergence to Gaussian occurs more gradually with increasing $p$ than in continuous data. Significant convergence seems to occur when $p \geq 1000$, however, this is actually a relatively small number of features in the context of GWAS. Considering a realistic number of features for GWAS, the normality assumption of GM distances holds. This metric has the slowest convergence to Gaussian among all we have considered. P values from Shapiro-Wilk test, where the null hypothesis is a Gaussian distribution.

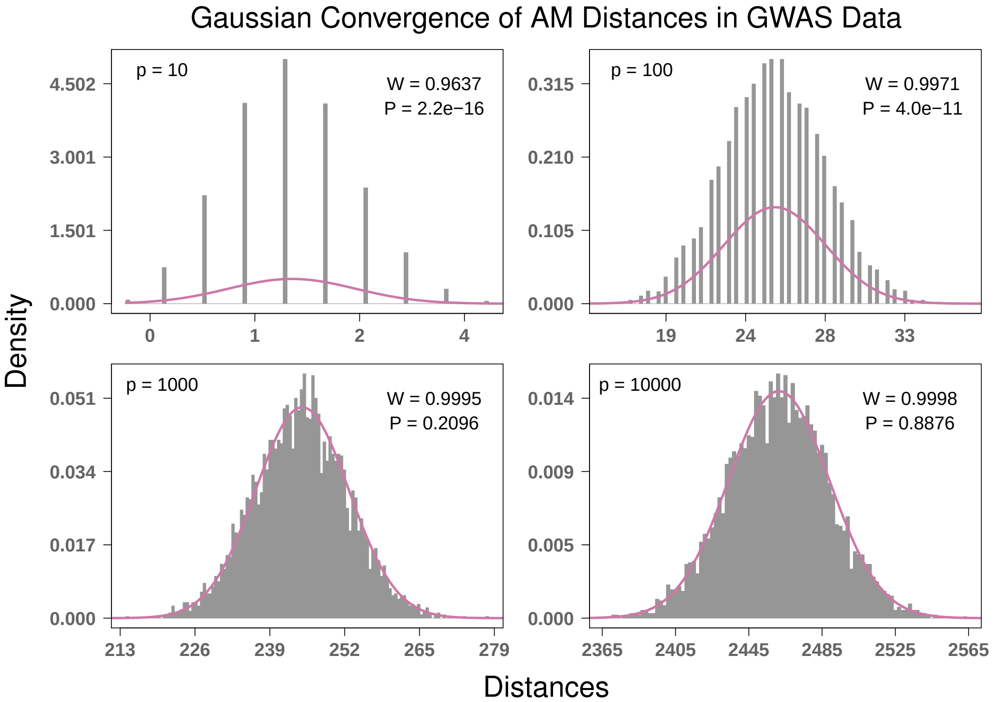


**Figure S5.** Convergence to Gaussian for AM distances for simulated binomial GWAS data with $m = 100$ instances and $p = 10, 100, 1000$, and 10000 attributes. The average MAF was set to 0.205 for all simulations. Convergence to Gaussian occurs more gradually with increasing $p$ than in continuous data. Significant convergence seems to occur when $p \geq 1000$, however, this is actually a relatively small number of features in the context of GWAS. Considering a realistic number of features for GWAS, the normality assumption of AM distances holds. This metric has the slightly faster convergence to Gaussian than the GM metric, which is probably due to the fact that the AM metric has one more value in its range (e.g., 1/2). P values from Shapiro-Wilk test, where the null hypothesis is a Gaussian distribution.

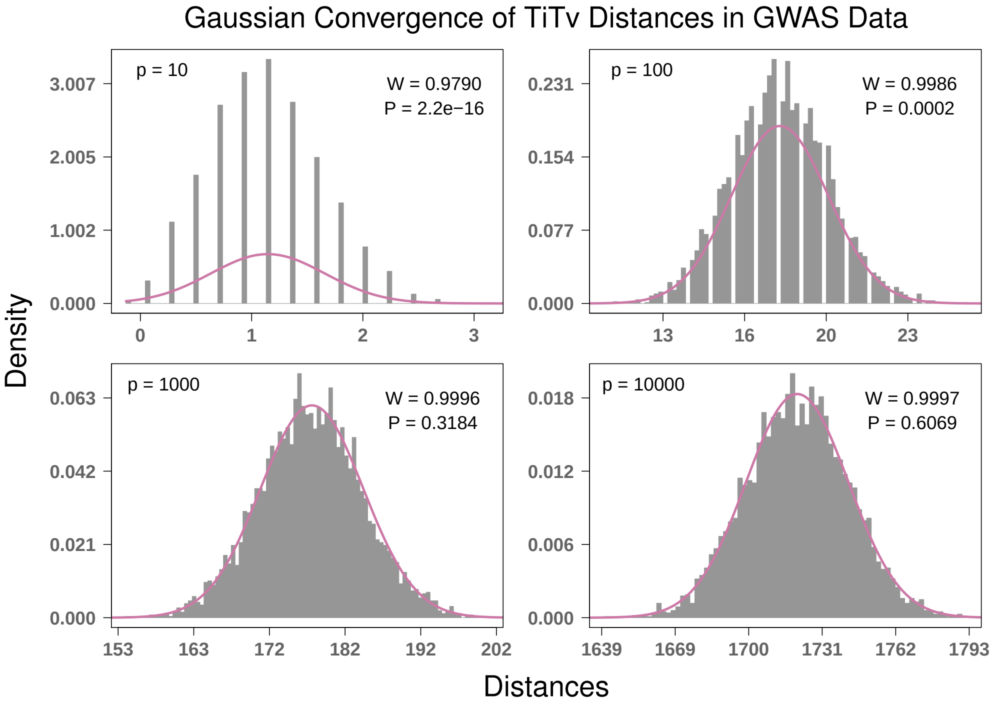


**Figure S6.** Convergence to Gaussian for TiTv distances for simulated binomial GWAS data with $m = 100$ instances and $p = 10, 100, 1000$, and 10000 attributes. The average MAF was set to 0.205 for all simulations and the Ti/Tv ratio ($\eta$) was set to 2. Convergence to Gaussian occurs more gradually with increasing $p$ than in continuous data. Significant convergence seems to occur when $p \geq 1000$, however, this is actually a relatively small number of features in the context of GWAS. Considering a realistic number of features for GWAS, the normality assumption of TiTv distances holds. This metric has the significantly faster convergence to Gaussian than the AM metric, which is probably due to the fact that the TiTv metric contains 2 more values in its range (e.g., 1/4 & 3/4). P values from Shapiro-Wilk test, where the null hypothesis is a Gaussian distribution.

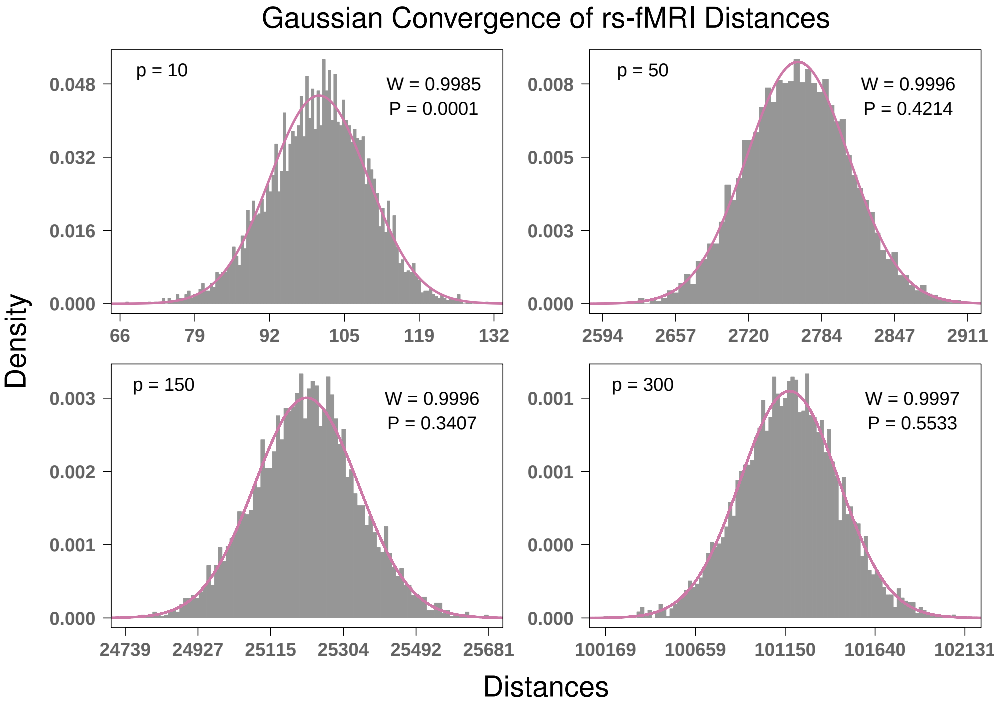


**Figure S7.** Convergence to Gaussian for rs-fMRI distances for simulated correlation matrices with $m = 100$ instances and $p = 10, 50, 150$, and 300 attributes (or ROIs). Correlation matrices were generated for each instance from random normal $m \times p$ data matrices. Each correlation matrix was then stretched out into a long vector, Fisher r-to-z transformed, stored in a $p(p-1) \times m$ matrix, and standardized so that the $m$ columns are mean 0 and unit variance. Convergence to Gaussian occurs very rapidly for this data because the dimensions are larger than a typical $m \times p$ data set. The large attribute dimension $p(p-1)$ means that there are significantly more terms in each sum to compute pairwise distances. Therefore, Classical Central Limit Theorem dictates that distances in this context will be closer to Gaussian. P values from Shapiro-Wilk test, where the null hypothesis is a Gaussian distribution.

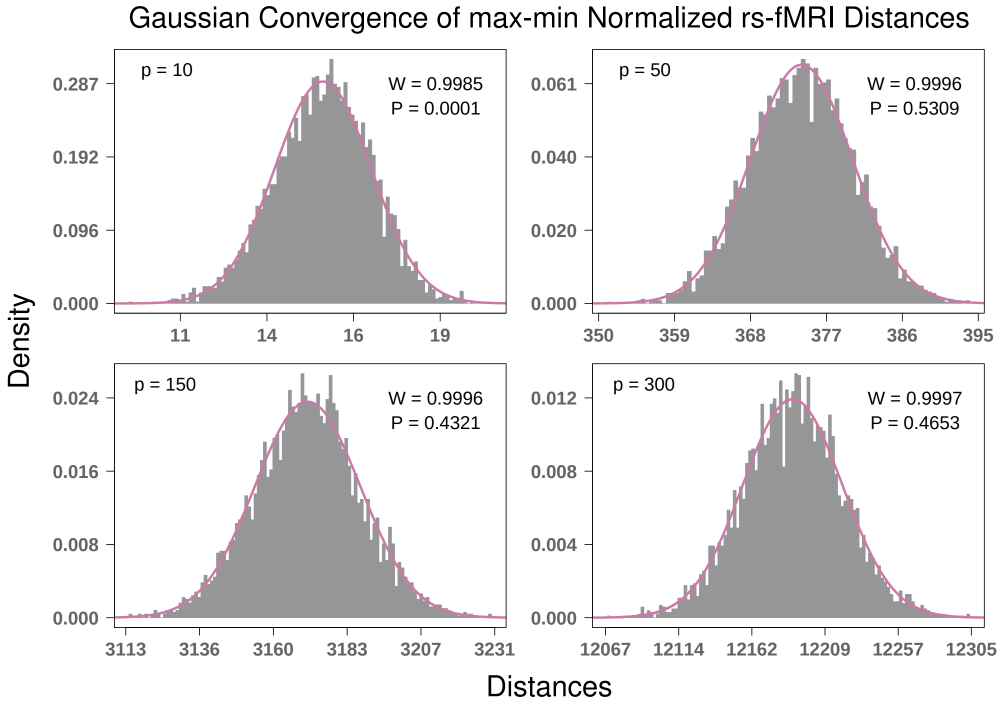


**Figure S8.** Convergence to Gaussian for max-min normalized rs-fMRI distances for simulated correlation matrices with $m = 100$ instances and $p = 10, 50, 150$, and 300 attributes (or ROIs). Correlation matrices were generated for each instance from random normal $m \times p$ data matrices. Each correlation matrix was then stretched out into a long vector, Fisher r-to-z transformed, stored in a $p(p-1) \times m$ matrix, and standardized so that the $m$ columns are mean 0 and unit variance. Convergence to Gaussian occurs approximately as rapidly as the standard rs-fMRI metric. Just as in the standard rs-fMRI metric, the large attribute dimension $p(p-1)$ means that there are significantly more terms in each sum to compute pairwise distances. Therefore, Classical Central Limit Theorem dictates that distances in this context will be closer to Gaussian. P values from Shapiro-Wilk test, where the null hypothesis is a Gaussian distribution.

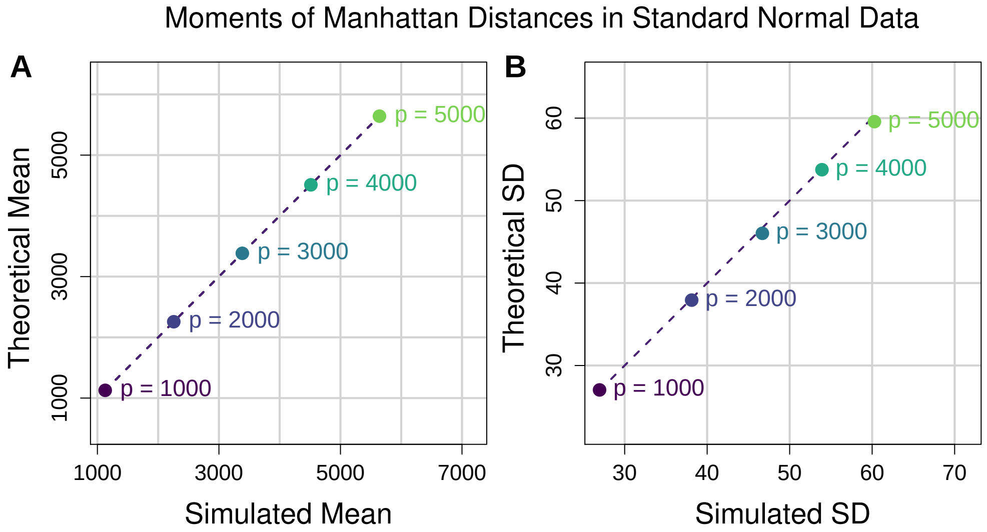


**Figure S9.** Comparison of theoretical and simulated moments of Manhattan distances in standard normal data. (**A**) Scatter plot of theoretical vs simulated mean Manhattan distance. Each point represents a different number of attributes $p$. For each value of $p$ we fixed $m = 100$ and generated 20 distance matrices from standard normal data and computed the average simulated pairwise distance from the 20 iterations. The corresponding theoretical mean was then computed for each value of $p$ for comparison. The dashed line represents the identity (or $y = x$) line for reference. (**B**) Scatter plot of theoretical vs simulated standard deviation of Manhattan distance. These standard deviations come from the same random distance matrices for which mean distance was computed for **A**. Both theoretical mean and standard deviation approximate the simulated moments quite well.

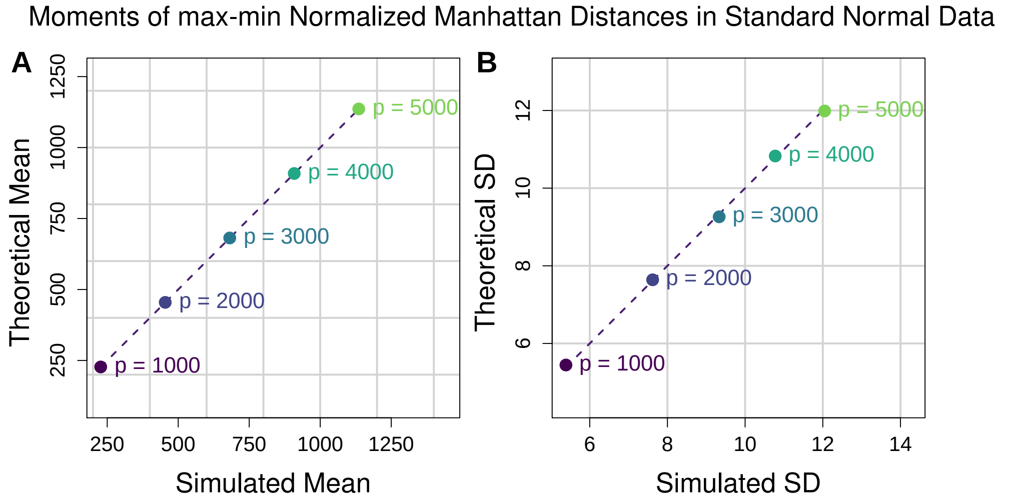


**Figure S10.** Comparison of theoretical and simulated moments of max-min normalized Manhattan distances in standard normal data. (**A**) Scatter plot of theoretical vs simulated mean max-min normalized Manhattan distance. Each point represents a different number of attributes $p$. For each value of $p$ we fixed $m = 100$ and generated 20 distance matrices from standard normal data and computed the average simulated pairwise distance from the 20 iterations. The corresponding theoretical mean was then computed for each value of $p$ for comparison. The dashed line represents the identity (or $y = x$) line for reference. (**B**) Scatter plot of theoretical vs simulated standard deviation of max-min normalized Manhattan distance. These standard deviations come from the same random distance matrices for which mean distance was computed for **A**. Both theoretical mean and standard deviation approximate the simulated moments quite well.

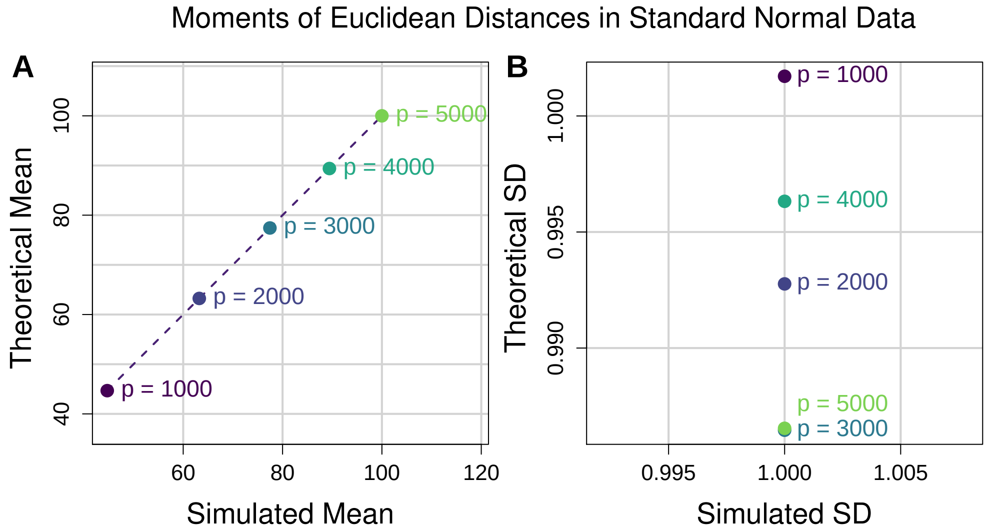


**Figure S11.** Comparison of theoretical and simulated moments of Euclidean distances in standard normal data. (**A**) Scatter plot of theoretical vs simulated mean Euclidean distance. Each point represents a different number of attributes $p$. For each value of $p$ we fixed $m = 100$ and generated 20 distance matrices from standard normal data and computed the average simulated pairwise distance from the 20 iterations. The corresponding theoretical mean was then computed for each value of $p$ for comparison. The dashed line represents the identity (or $y = x$) line for reference. (**B**) Scatter plot of theoretical vs simulated standard deviation of Euclidean distance. These standard deviations come from the same random distance matrices for which mean distance was computed for **A**. Theoretical and simulated means lie approximately on the identity line because the mean is proportional to attribute dimension $p$. Theoretical standard deviation is constant, which is why each horizontal coordinate is the same for $p = 1000, 2000, 3000, 4000,$ and 5000. The variation in sample standard deviation of Euclidean distance is quite small, so each simulated moment is clustered about 1.

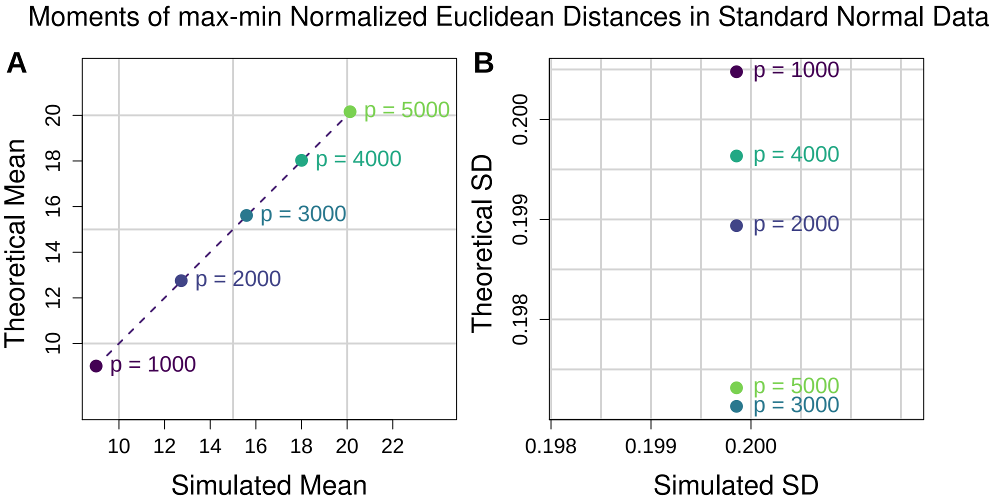


**Figure S12.** Comparison of theoretical and simulated moments of max-min normalized Euclidean distances in standard normal data. (**A**) Scatter plot of theoretical vs simulated mean max-min normalized Euclidean distance. Each point represents a different number of attributes $p$. For each value of $p$ we fixed $m = 100$ and generated 20 distance matrices from standard normal data and computed the average simulated pairwise distance from the 20 iterations. The corresponding theoretical mean was then computed for each value of $p$ for comparison. The dashed line represents the identity (or $y = x$) line for reference. (**B**) Scatter plot of theoretical vs simulated standard deviation of max-min normalized Euclidean distance. These standard deviations come from the same random distance matrices for which mean distance was computed for **A**. Theoretical and simulated means lie approximately on the identity line because the mean is proportional to $\sqrt{p}$. Theoretical standard deviation is a function of the fixed attribute dimension $m$, which is why each horizontal coordinate is the same for $p = 1000, 2000, 3000, 4000,$ and $5000$. The variation in sample standard deviation of max-min normalized Euclidean distance is quite small, so each simulated moment is clustered about the theoretical value that depends on $m$.

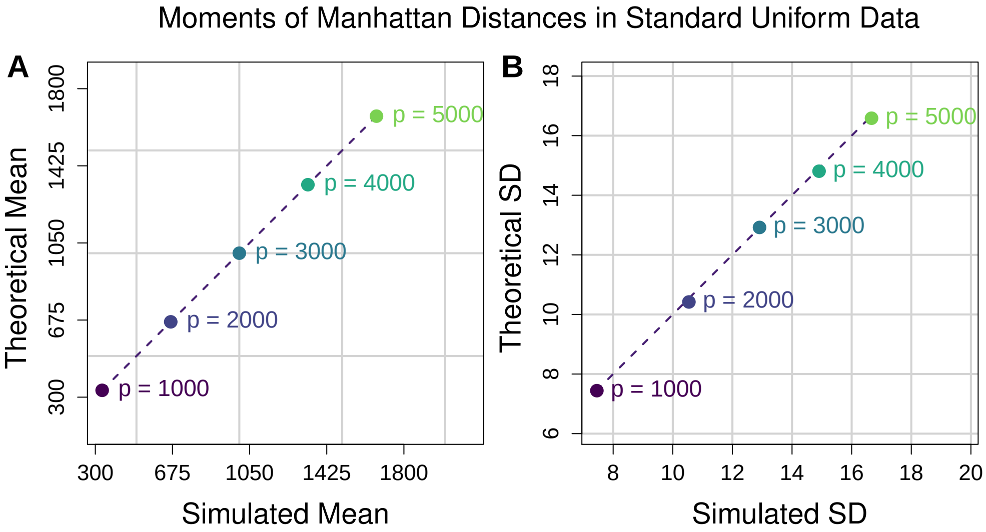


**Figure S13.** Comparison of theoretical and simulated moments of Manhattan distances in standard uniform data. (**A**) Scatter plot of theoretical vs simulated mean Manhattan distance. Each point represents a different number of attributes $p$. For each value of $p$ we fixed $m = 100$ and generated 20 distance matrices from standard uniform data and computed the average simulated pairwise distance from the 20 iterations. The corresponding theoretical mean was then computed for each value of $p$ for comparison. The dashed line represents the identity (or $y = x$) line for reference. (**B**) Scatter plot of theoretical vs simulated standard deviation of Manhattan distance. These standard deviations come from the same random distance matrices for which mean distance was computed for **A**. Both theoretical mean and standard deviation approximate the simulated moments quite well.

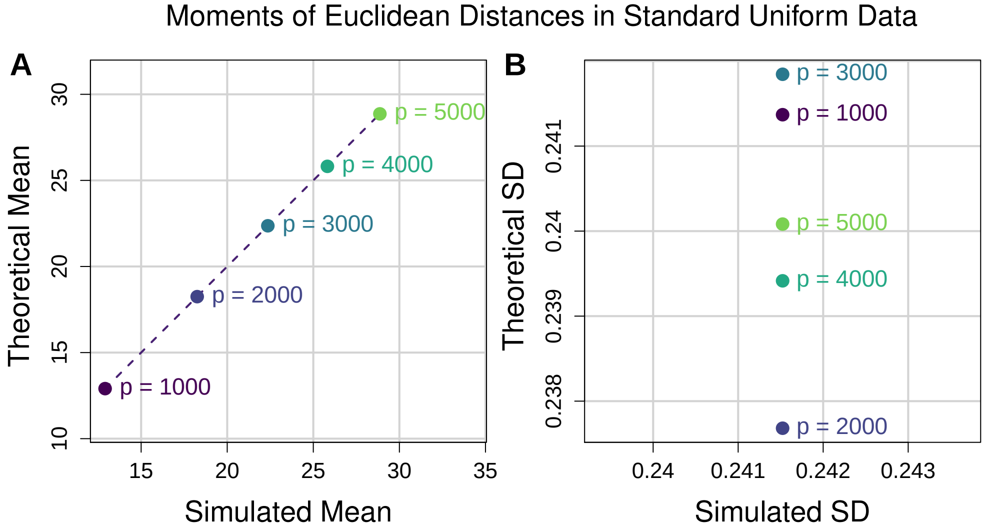


**Figure S14.** Comparison of theoretical and simulated moments of Euclidean distances in standard uniform data. (**A**) Scatter plot of theoretical vs simulated mean Euclidean distance. Each point represents a different number of attributes $p$. For each value of $p$ we fixed $m = 100$ and generated 20 distance matrices from standard uniform data and computed the average simulated pairwise distance from the 20 iterations. The corresponding theoretical mean was then computed for each value of $p$ for comparison. The dashed line represents the identity (or $y = x$) line for reference. (**B**) Scatter plot of theoretical vs simulated standard deviation of Euclidean distance. These standard deviations come from the same random distance matrices for which mean distance was computed for **A**. Theoretical and simulated means lie approximately on the identity line because the mean is proportional to attribute dimension $p$. Theoretical standard deviation is constant, which is why each horizontal coordinate is the same for $p = 1000, 2000, 3000, 4000,$ and $5000$. The variation in sample standard deviation of Euclidean distance is quite small, so each simulated moment is clustered about $7/120$.

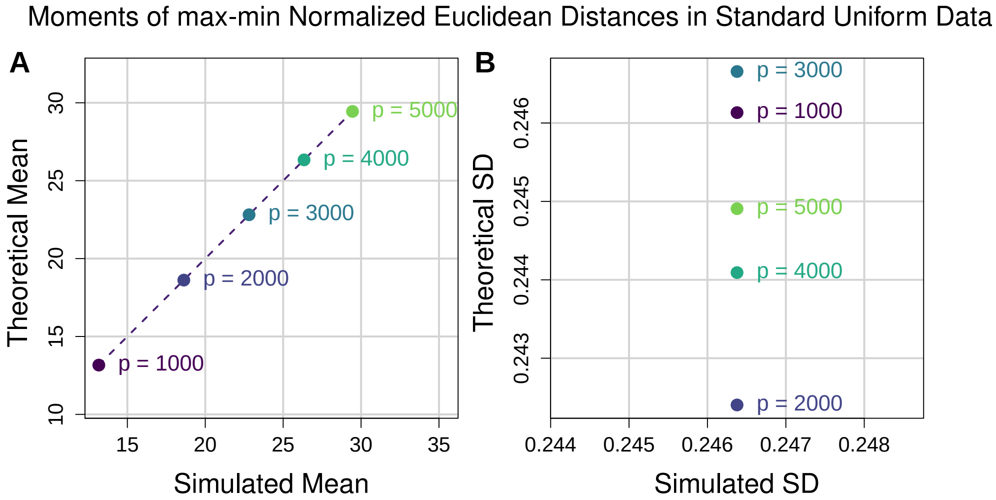


**Figure S15.** Comparison of theoretical and simulated moments of max-min normalized Euclidean distances in standard uniform data. (**A**) Scatter plot of theoretical vs simulated mean max-min normalized Euclidean distance. Each point represents a different number of attributes $p$. For each value of $p$ we fixed $m = 100$ and generated 20 distance matrices from standard uniform data and computed the average simulated pairwise distance from the 20 iterations. The corresponding theoretical mean was then computed for each value of $p$ for comparison. The dashed line represents the identity (or $y = x$) line for reference. (**B**) Scatter plot of theoretical vs simulated standard deviation of max-min normalized Euclidean distance. These standard deviations come from the same random distance matrices for which mean distance was computed for **A**. Theoretical and simulated means lie approximately on the identity line because the mean is proportional to $\sqrt{p}$. Theoretical standard deviation is a function of the fixed attribute dimension $m$, which is why each horizontal coordinate is the same for $p = 1000, 2000, 3000, 4000,$ and $5000$. The variation in sample standard deviation of max-min normalized Euclidean distance is quite small, so each simulated moment is clustered about the theoretical value that depends on $m$.

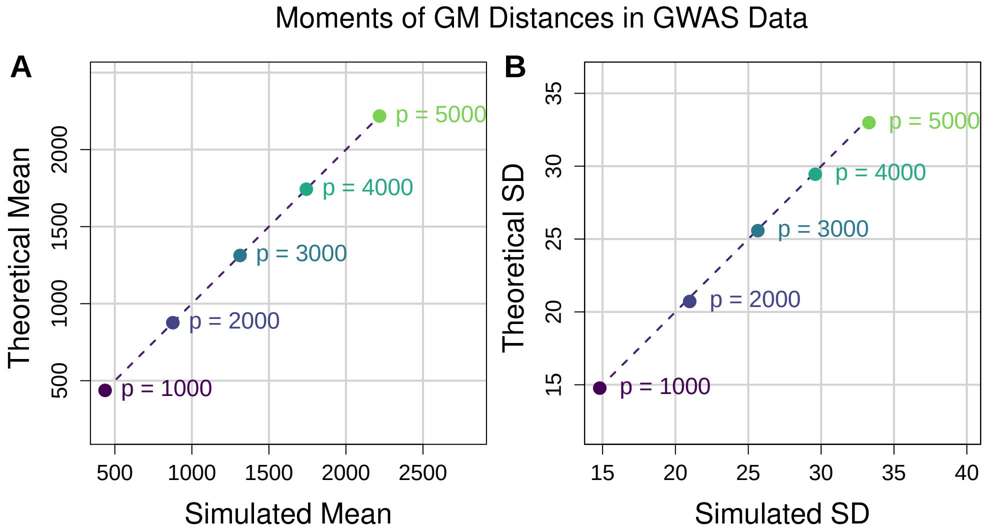


**Figure S16.** Comparison of theoretical and simulated moments of GM distances in binomial GWAS data. (**A**) Scatter plot of theoretical vs simulated mean GM distance. Each point represents a different number of attributes $p$. For each value of $p$ we fixed $m = 100$ and generated 20 distance matrices from binomial GWAS data and computed the average simulated pairwise distance from the 20 iterations. The corresponding theoretical mean was then computed for each value of $p$ for comparison. The dashed line represents the identity (or $y = x$) line for reference. (**B**) Scatter plot of theoretical vs simulated standard deviation of GM distance. These standard deviations come from the same random distance matrices for which mean distance was computed for **A**. Both theoretical mean and standard deviation approximate the simulated moments quite well.

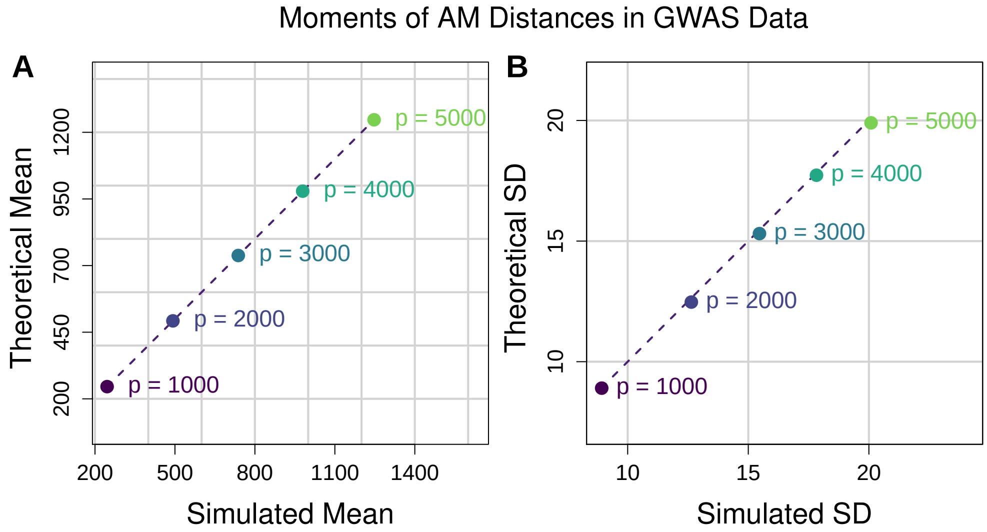


**Figure S17.** Comparison of theoretical and simulated moments of AM distances in binomial GWAS data. (**A**) Scatter plot of theoretical vs simulated mean AM distance. Each point represents a different number of attributes $p$. For each value of $p$ we fixed $m = 100$ and generated 20 distance matrices from binomial GWAS data and computed the average simulated pairwise distance from the 20 iterations. The corresponding theoretical mean was then computed for each value of $p$ for comparison. The dashed line represents the identity (or $y = x$) line for reference. (**B**) Scatter plot of theoretical vs simulated standard deviation of AM distance. These standard deviations come from the same random distance matrices for which mean distance was computed for **A**. Both theoretical mean and standard deviation approximate the simulated moments quite well.

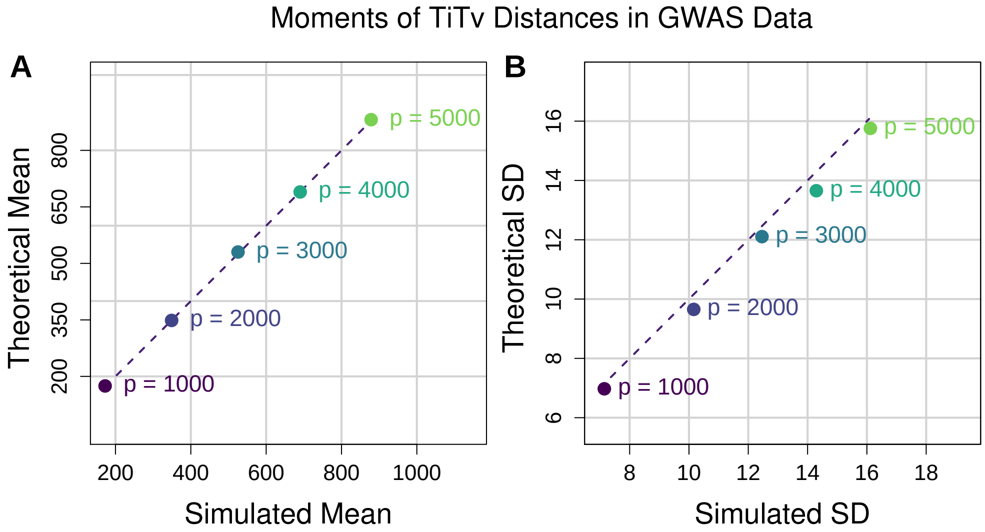


**Figure S18.** Comparison of theoretical and simulated moments of TiTv distances in binomial GWAS data. For each simulated data set, the average MAF was set to be 0.205 and the Ti/Tv ratio ($\eta$) was fixed to be 2. (**A**) Scatter plot of theoretical vs simulated mean TiTv distance. Each point represents a different number of attributes $p$. For each value of $p$ we fixed $m = 100$ and generated 20 distance matrices from binomial GWAS data and computed the average simulated pairwise distance from the 20 iterations. The corresponding theoretical mean was then computed for each value of $p$ for comparison. The dashed line represents the identity (or $y = x$) line for reference. (**B**) Scatter plot of theoretical vs simulated standard deviation of TiTv distance. These standard deviations come from the same random distance matrices for which mean distance was computed for **A**. Both theoretical mean and standard deviation approximate the simulated moments quite well.

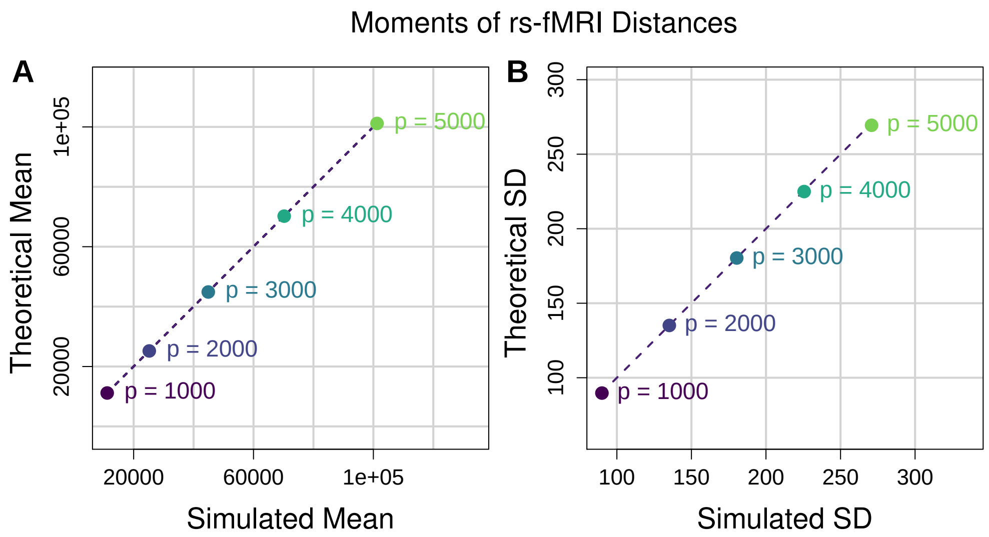


**Figure S19.** Comparison of theoretical and simulated moments of rs-fMRI distances from random correlation matrices. For each instance, we generated a $p \times p$ correlation matrix from a random $m \times p$ standard normal data set. We then stretched out each correlation matrix into a long vector, Fisher r-to-z transformed the correlations, stored the vector in a column of a large $p(p-1) \times m$ matrix, and then standardized columns to be mean 0 and unit variance. (**A**) Scatter plot of theoretical vs simulated mean rs-fMRI distance. Each point represents a different number of attributes $p$. For each value of $p$ we fixed $m = 100$ and generated 20 distance matrices from rs-fMRI data and computed the average simulated pairwise distance from the 20 iterations. The corresponding theoretical mean was then computed for each value of $p$ for comparison. The dashed line represents the identity (or $y = x$) line for reference. (**B**) Scatter plot of theoretical vs simulated standard deviation of rs-fMRI distance. These standard deviations come from the same random distance matrices for which mean distance was computed for **A**. Both theoretical mean and standard deviation approximate the simulated moments quite well.

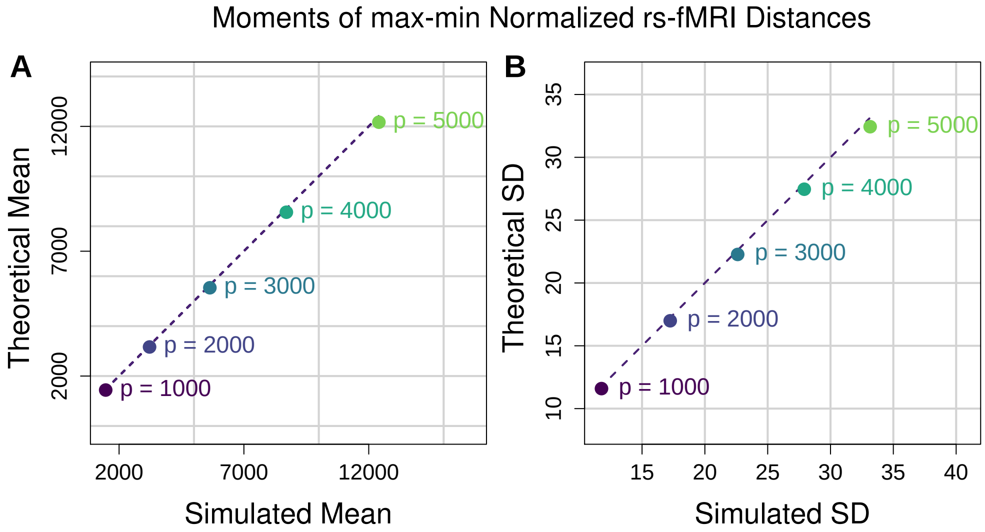


**Figure S20.** Comparison of theoretical and simulated moments of max-min normalized rs-fMRI distances from random correlation matrices. For each instance, we generated a $p \times p$ correlation matrix from a random $m \times p$ standard normal data set. We then stretched out each correlation matrix into a long vector, Fisher r-to-z transformed the correlations, stored the vector in a column of a large $p(p-1) \times m$ matrix, and then standardized columns to be mean 0 and unit variance. (**A**) Scatter plot of theoretical vs simulated mean max-min normalized rs-fMRI distance. Each point represents a different number of attributes $p$. For each value of $p$ we fixed $m = 100$ and generated 20 distance matrices from rs-fMRI data and computed the average simulated pairwise distance from the 20 iterations. The corresponding theoretical mean was then computed for each value of $p$ for comparison. The dashed line represents the identity (or $y = x$) line for reference. (**B**) Scatter plot of theoretical vs simulated standard deviation of max-min normalized rs-fMRI distance. These standard deviations come from the same random distance matrices for which mean distance was computed for **A**. Both theoretical mean and standard deviation approximate the simulated moments quite well.

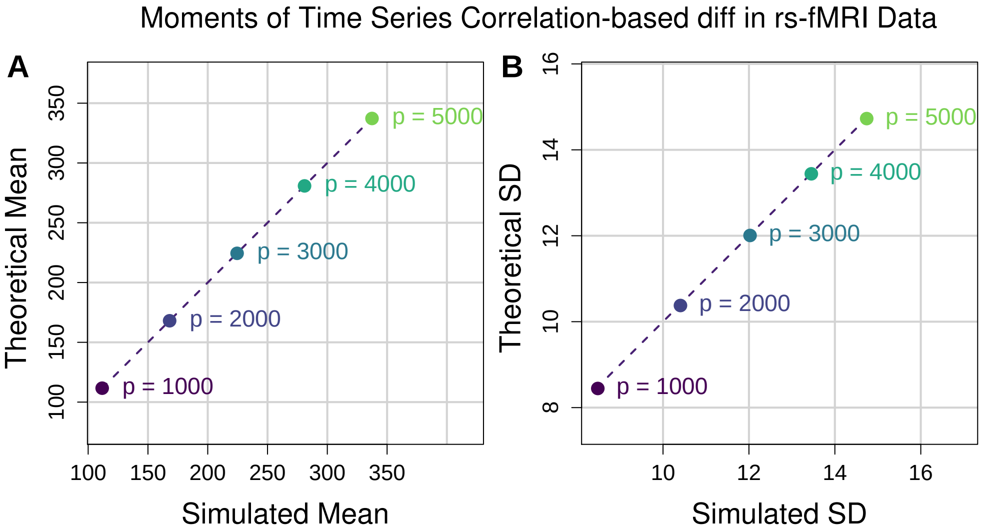


**Figure S21.** Comparison of theoretical and simulated moments of rs-fMRI diff metric from random correlation matrices. For each instance, we generated a $p \times p$ correlation matrix from a random $m \times p$ standard normal data set. We then stretched out each correlation matrix into a long vector, Fisher r-to-z transformed the correlations, stored the vector in a column of a large $p(p-1) \times m$ matrix, and then standardized columns to be mean 0 and unit variance. (**A**) Scatter plot of theoretical vs simulated mean rs-fMRI diff. Each point represents a different number of attributes $p$. For each value of $p$ we fixed $m = 100$ and generated 20 diff metric values from the rs-fMRI data and computed the average simulated diff from the 20 iterations. The corresponding theoretical mean was then computed for each value of $p$ for comparison. The dashed line represents the identity (or $y = x$) line for reference. (**B**) Scatter plot of theoretical vs simulated standard deviation of rs-fMRI diff. These standard deviations come from the same random diff values from which mean diffs were computed for **A**. Both theoretical mean and standard deviation approximate the simulated moments quite well.

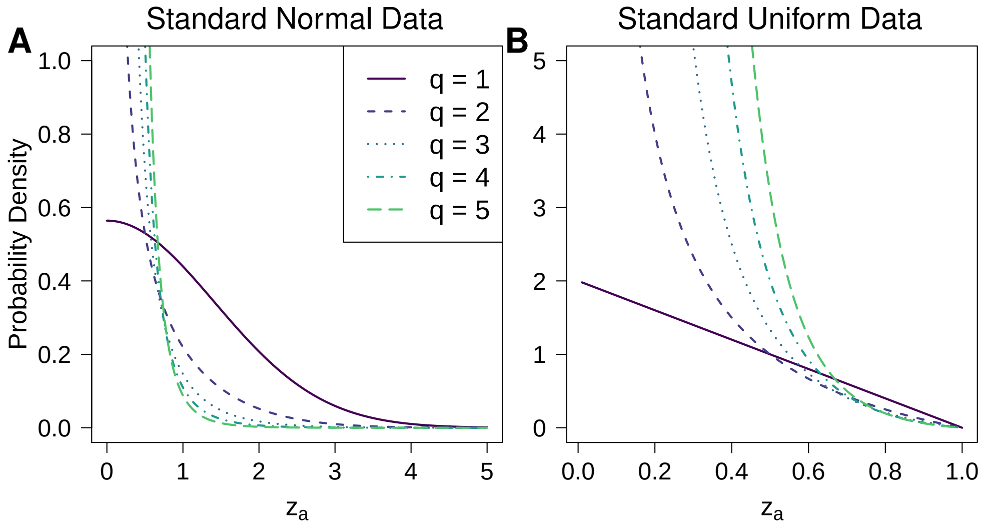


**Figure S22.** Density curves for one-dimensional projected distances (diffs) onto a fixed attribute $a$ for standard normal and standard uniform data. (**A**) Density curves for the distribution of attribute diff in standard normal data for $q = 1, 2, 3, 4$, and 5. This density is that of a Generalized Gamma distribution. For $q = 1$, this is also known as a half-normal distribution. (**B**) Density curves for the distribution of attribute diff in standard uniform data for $q = 1, 2, 3, 4$, and 5. This density is that of a Kumaraswamy distribution. For $q = 1$, this is also known as a triangular distribution.

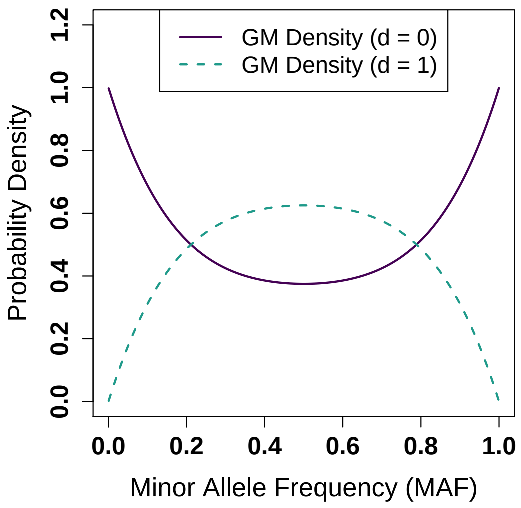


**Figure S23.** One-dimensional projected GM distance (diff) onto an attribute vs minor allele frequency (MAF). For each possible value of the GM diff (d $= 0, 1$), the exact density of the GM diff is plotted for all possible values of MAF. The expected value of MAF at a particular locus $a$ is $f_a$ for all $a \in \mathcal{A}$, where $f_a$ is the probability of a minor allele occurring at locus $a$. For each element $X_{ia}$ of the data matrix for a fixed attribute $a$, we have $X_{ia} \sim \mathcal{B}(2, f_a)$. Depending on the MAF, the frequency of GM diff taking on a value of 0 or 1 changes. For small MAF, the GM diff will be 0 most often. As MAF increases beyond 0.5, the minor allele switches.

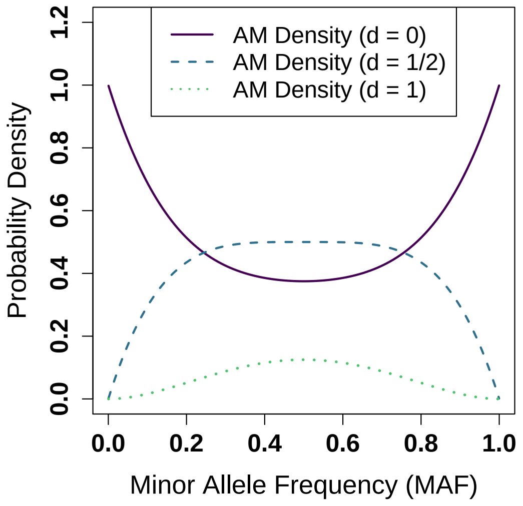


**Figure S24.** One-dimensional projected AM distance (diff) onto an attribute vs minor allele frequency (MAF). For each possible value of the AM diff (d $= 0, 1/2, 1$), the exact density of the AM diff is plotted for all possible values of MAF. The expected value of MAF at a particular locus $a$ is $f_a$ for all $a \in \mathcal{A}$, where $f_a$ is the probability of a minor allele occurring at locus $a$. For each element $X_{ia}$ of the data matrix for a fixed attribute $a$, we have $X_{ia} \sim \mathcal{B}(2, f_a)$. Depending on the MAF, the frequency of AM diff taking on a value of 0, 1/2, or 1 changes. For small MAF, the AM diff will be 0 most often. For large MAF ($\approx$0.5), the AM diff will be mostly 1/2 with 1 being the second most common value. The least common value of the AM diff is 1. As MAF increases beyond 0.5, the minor allele switches.

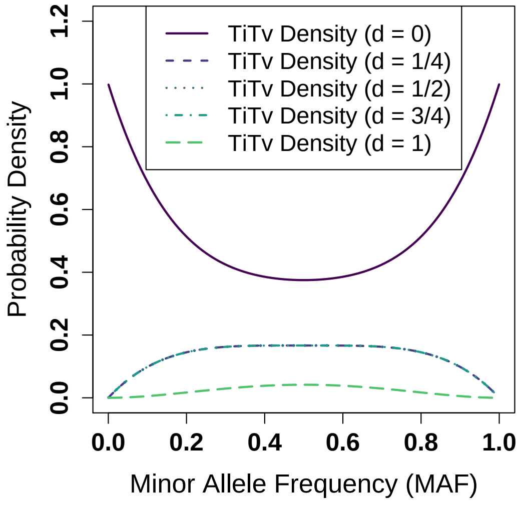


**Figure S25.** One-dimensional projected TiTv distance (diff) onto an attribute vs minor allele frequency (MAF). For each possible value of the TiTv diff (d $= 0, 1/4, 1/2, 3/4, 1$), the exact density of the TiTv diff is plotted for all possible values of MAF. The Ti/Tv ratio $\eta$ was fixed to be 2. The expected value of MAF at a particular locus $a$ is $f_a$ for all $a \in \mathcal{A}$, where $f_a$ is the probability of a minor allele occurring at locus $a$. For each element $X_{ia}$ of the data matrix for a fixed attribute $a$, we have $X_{ia} \sim \mathcal{B}(2, f_a)$. Depending on the MAF, the frequency of TiTv diff taking on a value of 0, 1/4, 1/2, 3/4, or 1 changes. The density of the TiTv diff for d $= 1/4, 1/2,$ and $3/4$ has the same resulting curve as a function of MAF. The most common value of TiTv diff at any MAF is 0, the second most common is 1/4, 1/2, or 3/4, and the least common is 1. As MAF increases beyond 0.5, the minor allele switches.
